# Late Cretaceous origins for major nightshade lineages from total evidence timetree analysis

**DOI:** 10.1101/2025.07.22.666174

**Authors:** Ixchel S. González Ramírez, Rocío Deanna, Stacey D. Smith

## Abstract

**Background and Aims:** The timing of the radiation of nightshades (Solanaceae) has been contentious in the literature, with estimates of the crown age ranging from ca. 30 to 70 Mya (mid-Oligocene to late Cretaceous). The tempo of diversification of major lineages within the family (e.g., berries, tobaccos) has been equally challenging to resolve, in large part because of the paucity of fossil information. Recently described fossils present an opportunity to revisit the timing of nightshade diversification using more powerful model-based methods. Here, we simultaneously infer divergence times within Solanaceae and the placement of a select set of well-preserved and morphologically diverse fruit and seed fossils.

**Methods:** We assembled a family-wide morphological dataset, including 17 categorical and eight continuous characters, for 134 living and 14 fossil Solanaceae taxa, as well as sequence data for the extant taxa. We implemented a Bayesian total evidence dating analysis in RevBayes using (a time continuous and a time heterogeneous) fossilized birth-death model and models of character evolution for each type of data.

**Key Results:** The origin of Solanaceae was ∼98 Mya, and the major splits were roughly three-fold older than previously estimated. Although the 14 fossil taxa were phylogenetically placed with different degrees of confidence, we identified a fruit fossil and a seed fossil whose affinities were strongly supported. Moreover, most of the fossils lacking a precise placement were nevertheless confidently inferred to belong to the large berry clade.

**Conclusions:** Our study provides an example of how a sophisticated model used on a carefully assembled dataset can shed light on the timing of the evolution of a group, while accounting for phylogenetic uncertainty. The timetree we present here provides a temporal framework for further research, from comparative genomics and patterns of diversification to trait evolution and biogeography.

## INTRODUCTION

Bursts of species diversification are a hallmark of evolutionary history. These radiations often coincide with major morphological innovations (e.g., Troyer *et al*., 2024; Liu *et al.,* 2024), but they may equally be driven by extrinsic factors such as the expansion of ecological opportunities or changing climate (Blaimer *et al*., 2023; Dellinger *et al*., 2024). While examples from the animal literature are among the best known, such as the radiation of placental mammals or metazoan phyla (e.g., Lee *et al*., 2013; Foley *et al*., 2023), radiation events have also shaped much of plant evolutionary history (Wang *et al*., 2009; Rothfels *et al*., 2012; Zuntini *et al*., 2024). These periods of rapid diversification pose significant challenges for phylogenetic inference, in terms of both estimating relationships among lineages and the timing of lineage-splitting events (Whitfield and Lockhart, 2007; Léveillé-Bourret *et al*., 2017).

The explicit incorporation of fossil lineages into total-evidence phylogenetic analyses presents a new approach for simultaneously estimating the timing and order of evolutionary events. While fossils have long been used to calibrate nodes of the tree (e.g., Sanderson and Doyle, 2001; Magallón, 2004; Won *et al*., 2006), they can also be treated as sampled taxa, allowing their age and placement to inform the shape of the tree. Specifically, when using models that include the fossil generating process, such as the fossilized birth-death (FBD) model (Stader, 2010; Heath *et al*., 2014; Stadler *et al*., 2018), the timing of fossil occurrences will make some areas of tree space highly improbable (e.g., those that place the youngest and oldest fossils as sister taxa) while favoring others (Barido-Sottani *et al*., 2020; Mongiardino Koch *et al*., 2021). Additionally, the joint inference of topology and divergence times is a natural framework to account for the effect that topological uncertainty might have on divergence times. This is a particularly important feature when working with fossils where uncertainty in placement is expected. Nevertheless, compared to node-dating approaches (reviewed in Forest, 2009), fully model-based total evidence methods have been applied relatively rarely to empirical systems (reviewed in Mulvey *et al*., 2025), and even more rarely in plant clades (e.g., Renner *et al*., 2016; May *et al*., 2021; Coiro *et al*., 2023). The paucity of these studies likely reflects not only the limited fossil record for many plant families (e.g., Ramírez *et al*., 2007; Huang *et al*., 2024) but also the limited trait data for extant taxa, particularly for the micro-morphological features commonly examined in paleobotanical studies but rarely documented in typical taxonomic treatments (e.g., pollen and seed ultrastructure, wood anatomy, venation patterns of calyces or leaves).

Here we investigate divergence times and phylogenetic relationships in the large and cosmopolitan nightshade family (Solanaceae) using a newly developed family-wide morphological dataset spanning living and fossil taxa. Several previous studies have estimated divergence times for the family and clades within, although the dates estimated have varied widely (Table S1). Studies using fossil calibrations (i.e., node dating) at the angiosperm level have suggested stem ages for the family ranging from 66.6 to 82.5 Ma (Magallón *et al*., 2015; Barba-Montoya *et al*., 2018; Li *et al*., 2019; Silvestro *et al*., 2021), and those including multiple species within the family estimate crown ages between 30 and 73.3 Ma (Särkinen *et al*., 2013; De-Silva *et al*., 2017; Huang *et al*., 2023). The minimum age of the large berry clade, comprising roughly 3/4 of the ca. 2,850 species in the family, however, has been solidified with the recent discoveries of fossil berries from the early Eocene at 52 Ma (Wilf *et al*., 2017; Deanna *et al*., 2020, 2023). Nevertheless, relationships among the major lineages of berries (e.g., the peppers, tomatillos, wolfberries, and tomatoes) and the timing of these splits remain uncertain (Särkinen *et al*., 2013; Cao *et al*., 2021; Powell *et al*., 2022; Huang *et al*., 2023).

In order to explore relationships among major nightshade lineages and the timeline of their emergence, this study incorporates both discrete and continuous morphological characters within the Bayesian fossilized birth-death (FBD) modeling framework. Like other total evidence dating approaches (e.g., Pyron, 2011; Ronquist *et al*., 2012), fossils are treated as tips and are scored for the same characters as the extant taxa in order for their position to be estimated. The FBD model has become increasingly popular as it allows for biologically realistic prior distributions based on speciation, extinction, and fossilization (Heath *et al*., 2014), although it remains challenging to apply in many cases due to the model complexity and biases in the fossil record (Mulvey *et al*., 2025). As with any Bayesian approach, a range of characters can be included so long as they are matched with an appropriate model (e.g., Brownian motion for continuous characters or Mk models for discrete characters). In this study, besides categorical traits, we also include continuous morphological data. Although morphometric data have been used for topological inference (Parins-Fukuchi, 2018) and for divergence time studies with other Bayesian frameworks (Alvarez-Carretero *et al*., 2019), our work is one of the few studies that include continuous traits in a total-evidence dating analysis (e.g., Zhang *et al*., 2024). Morphometric data have long served as important sources of taxonomic characters for plant fossils (e.g., Kara *et al*., 2024; Brightly *et al*., 2024; Wu *et al*., 2025), and Solanaceae are no exception (Franco and Brea, 2023; Deanna *et al*., 2023, 2025a). Building on decades of molecular systematic studies in Solanaceae (Olmstead & Palmer, 1992; Olmstead *et al*., 2008; Särkinen *et al*., 2013), our novel continuous and discrete morphological dataset and tailored model-based analyses provide a rich and detailed picture of the family’s diversification, the first to put fossils in their rightful place as unique products of nightshade evolutionary history.

## MATERIALS AND METHODS

### Taxon sampling

We included 134 extant species of Solanaceae, encompassing 97 genera of the 101 in the family. Several genera were represented by multiple tips, either because of the large size of the genus (e.g., *Solanum* L.) or because of their predicted close relationship to fossil taxa (e.g, *Physalis* L., Deanna *et al*., 2023). Within the family’s largest genus, *Solanum*, we sampled 32 species in order to span the major lineages that have emerged from focused studies in this clade (Tepe *et al*., 2016; Gagnon *et al*., 2022; Messeder *et al*., 2024). We also considered the availability of molecular sequence data and specimens for scoring and selecting individual tips. Although four genera were not included here (*Cataracta* Zamora-Tav., O.Vargas & M.Martínez, *Duboisia* R.Br.*, Henoonia* Griseb.*, Melananthus* Walp.), these have been included in previous studies and are confidently placed within the family (Deanna *et al*., 2025b). Thus, our sampling is designed to provide a scaffold for the family, which will allow us to estimate the timing of splits among important taxonomic groups.

Our dataset also included 14 fossil taxa based on recent discoveries and taxonomic revision (Table S2; Wilf *et al*., 2017; Deanna *et al*., 2020, 2023, 2025a). Four of these are fruit macrofossils (*Eophysaloides inflata* Martínez-A. & Deanna, *Lycianthioides calycina* Deanna & Manchester, *Physalis infinemundi* Wilf, and *P. hunickenii* Deanna Wilf & Gandolfo), all of which are berries, and the remaining ten fossils are seeds. The nightshade seed fossil record includes 11 distinct taxa, whose taxonomy has been recently revised (Deanna *et al*., 2025a). We excluded the fossil *Physalis pliocenica* Szafer because only a short description of the specimen was found, and it lacks a morphometric analysis (Szafer, 1946). For most of these fossil fruit and seed taxa, we scored the type specimen (except for *Solanum foveolatum* Negru and *Solanispermum reniforme* Chandle*r,* for which we used a different specimen due to lack of access to the holotype and lack of information in the bibliography) and used the stratigraphic uncertainty as age input for the total evidence dating analyses.

### Morphological and molecular dataset construction

We scored a total of 25 morphological characters for seeds and fruits, 17 of which were discrete and eight continuous. The discrete characters included 14 binary variables (fruiting calyx inflation, fruiting calyx venation, distinctiveness of widest veins that terminate in lobe tips, presence of secondary veins that emerge from the base and their forking, basal invagination and angle of the fruiting calyx, presence of calyx teeth, seed compression, hilum position in the seed, hilar-chalazal cavity, exotestal wall shape, seed wings, seed elaiosomes) and three categorical (fruit type, fruiting calyx lobe sinus shape, embryo shape). These characters were selected due to their variation across the fossils (Deanna *et al*., 2023, 2025a) and their utility in previous family-level classifications (Hunziker *et al*., 2001). The morphometric dataset included six fruit traits (fruit length-width ratio, fruiting pedicel length, fruiting calyx length, fruiting calyx length-width ratio, length of the longest fruiting calyx lobe, and fruiting calyx lobe length-width ratio) and two seed traits (seed length and length-width ratio). The full coding scheme for all characters is given in Table S3 and is adapted from Deanna *et al*. (2023) for fruits and Deanna *et al*. (2025a) for seeds.

For the molecular dataset, we assembled sequence data for the 134 extant taxa from 10 nuclear and plastid markers commonly used for molecular systematic studies of Solanaceae: ITS, *LEAFY*, *waxy*, *matK*, *ndhF*, the *ndhF*-*rpl32* spacer, *psbA*, *rbcL*, the *trnL-trnF* spacer, and the *trnS-trnG* spacer. These data were gathered from Genbank using the PyPHLAWD pipeline (Smith & Walker, 2018), and each region was aligned using MUSCLE (Edgar, 2004) as implemented in Mega v. 12 (Kumar *et al*., 2024).

### Preliminary analyses

Implementing a complex model where multiple processes evolve jointly–i.e., morphological evolution, molecular evolution, and the tree generating process–requires the specification of multiple submodels that could be analyzed independently. While joint modeling has a lot of advantages, like allowing different components of the model to inform each other (e.g., ages informing topology), it relies on each subpart of the model working adequately. For this reason, we conducted multiple preliminary analyses for different submodels of our final analysis to ensure that each of the components of the main analysis produced results consistent with our empirical expectations. First, we inferred ML gene trees for each of the molecular markers used in this study using IQ-Tree2 (Minh *et al*., 2020). From these runs, we identified three taxa with long branches, which we were able to remedy by correcting an erroneous sequence for one species and adding additional sequences where the pipeline had failed to recover them from Genbank (two species).

Next, we conducted phylogenetic inferences using only morphological data under an Mk + Gamma model (for the discrete traits) and a Brownian motion model (for continuous traits) using a uniform tree prior to get a sense of the amount of information present without the sequence data. Finally, we conducted a joint analysis of all the data types (i.e., DNA, categorical traits, and continuous traits) using a uniform tree prior on the topology. This model employed the same character evolution data as the final TED analysis (Mk + Gamma, Brownian motion, and GTR + Gamma) without inferring divergence times. These ‘no-time’ trees from this final, corrected dataset allowed us to observe the predicted placement of the fossils using morphology alone.

### Bayesian total evidence dating analysis

As described by Warnock and Wright (2020), a Bayesian approach to inferring the divergence times of a lineage requires the specification of three components of the general model: 1) a model of character evolution that describes the way traits evolve, 2) a clock model that describes the rates at which characters evolve, and 3) a tree model that describes the process that generates the tree. In this study, we used a GTR + G to model DNA sequence evolution. The data was partitioned by marker, and we allowed four different categories of rate variation in the Gamma distribution. For the discrete morphological data, we used the Mk model (Pagel, 1994; Lewis, 2001) with all the transition rates among states for a given character being the same. In order to accommodate variation in rates across characters, we used a discretized Gamma distribution with four rate categories, analogous to the approach for modeling sequence evolution. Finally, we modeled the evolution of continuous traits as evolving under Brownian motion, with each trait having its own global rate.

Next, we assigned the three clock models for each of the three types of trait data, i.e., sequence data, categorical morphological traits, and continuous morphological traits. First, we used an uncorrelated log-normal clock (UCLN) to model the rates of evolution of sequence data (Drummond *et al*., 2006), where the rate of each branch was drawn from a shared log-normal distribution. Relaxed-clock models, such as UCLN, accommodate variation in rates across lineages and have repeatedly outperformed strict clocks in simulated (Crisp *et al*., 2014) and empirical (e.g., Lepage *et al*., 2007) datasets. Second, since all the traits (e.g., morphological and molecular) of a lineage evolve under the same tree topology, and given the overwhelming amount of molecular data in comparison to morphological data, we linked the rates of morphological evolution to the molecular clock. This is achieved by computing the rate of morphological evolution for each branch as the product of the rate of molecular change from the UCLN molecular clock by a global morphological rate parameter. The implicit assumption of this decision is that we expect lineages with a faster molecular evolution rate to also have a faster morphological rate. Finally, we allowed each continuous character to have its own global rate of evolution, independent of the molecular and discrete data. These elements are depicted graphically in Fig. S1.

For the tree-generating process, we used the Fossilized Birth-Death (FBD) process (Heath *et al*., 2014). In this model, diversification is defined by three rates: the speciation rate (λ), the extinction rate (μ), and the fossilization rate (ψ). In the simplest case, these rates are assumed to be constant across lineages and over time. Given the likely depth of the Solanaceae phylogeny (at least to the Eocene; Wilf *et al*., 2017), we also considered a time-heterogeneous model (as in May *et al*., 2021), in which the FBD parameters were allowed to vary at fixed intervals corresponding to the geological time periods as delimited in the International Stratigraphic Chart. The intervals that were relevant to our study were Pleistocene, Pliocene, Miocene, Oligocene, Eocene, Paleocene, Upper Cretaceous, and Lower Cretaceous (detailed time intervals are provided in the data repository). Geological boundaries are considered to be correlated with periods of large geological and biological changes, and as such we might expect changes in the diversification dynamics of major clades such as Solanaceae to roughly correspond to these intervals.

### Model estimation and convergence

We estimated model parameters, including the tree topology and divergence times, using MCMC in RevBayes (Höhna *et al*., 2016). The input dataset consisted of the two morphological data matrices (discrete and continuous) for 148 taxa (134 extant + 14 fossils), the 10 molecular alignments, and the age of the fossils, expressed as intervals that reflect our uncertainty about the age estimate (Table S3). In implementing the full model, including divergence time estimation, we incorporated two sets of topological constraints to help convergence. First, we created a three-taxon constraint tree of extant species with *Petunia interior* more closely related to *Physalis angulata* L. than to *Schizanthus grahamii* Gillies. This constraint tree is confidently based on several previous family-level analyses (Särkinen *et al*., 2013; Ng and Smith, 2016; Deanna *et al*., 2025b). We also imposed two constraints on fossil taxa. First, the two fossil *Physalis* fruit fossils *(P. hunickeni* and *P. infinemundi*) were constrained to be monophyletic with the two extant representatives of *Physalis* in our dataset (*P. angulata* and *P. peruviana* L.), as supported in our ‘no-time’ tree (Fig. S2) and previous morphological analyses (Wilf *et al*., 2017, Deanna *et al*., 2020, 2023). Second, we constrained *Hyoscyamus undulatus* Deanna & S.D.Sm. to form a clade with the extant *Hyoscyamus niger* L. as they share a terminal hilum and group together closely in morphometric analyses (Deanna *et al*., 2025a) and in our no-time tree (Fig. S2).

In addition to these topological constraints, we placed an origin time constraint on the FBD process prior. Specifically, we used a uniform prior distribution from 55 to 100 mya, which represents the 95% HPD estimate for the stem age of Solanaceae according to Silvestro *et al*. (2021). We chose to specify a prior on the origin time (as opposed to the root age) to account for the possibility of fossils being inferred as originating from the stem lineage of Solanaceae. As implemented by the ‘dnFBDP()’ prior distribution in RevBayes, this setting starts the FBD process from the stem branch that gives rise to the phylogeny. With these constraints, we ran MCMC chains for the two types of analyses (*i.e.*, time homogeneous and time heterogeneous) for 8,000 generations (with 381 moves per generation) with a burning phase of 200 generations. The convergence and mixing of the parameters were visually evaluated using Tracer (Rambault *et al*., 2018), and topological convergence was assessed using the R package RWTY (Warren *et al*., 2017).

### Comparing the time-homogenous and heterogenous models

For each analysis (i.e. time homogenous and time heterogeneous) we pooled the posterior sample of trees obtained from the two independent MCMC chains, obtaining a pooled tree sample for each analysis. Next, we compared the divergence time estimates from the two models by extracting the age of seven lineages of Solanaceae from the pooled posterior distributions of extant trees for each model. The seven clades of interest were /Capsiceae, /Cestroideae, /Datureae, /Dodecachroma, /Solanoideae, as well as the genus *Solanum* and the whole family. After extracting the ages for each tree in the sample, we summarized the results using violin plots. As the results appeared very similar here and in the RWTY plots, we chose to focus on the time-heterogeneous pooled tree sample for downstream steps.

We summarized the speciation, extinction, diversification, and fossilization rates obtained from this model using RevGadgets (Tribble *et al*., 2022). For each time interval, we plotted the average and the 95% highest posterior density for each parameter.

### Exploration of fossil placements

Because computing summary trees including multiple fossils at a time conflates the uncertainty associated with each of the fossils, we took a “one at a time” approach to explore the certainty of fossil placement in the phylogeny of extant Solanaceae. For each fossil, we pruned all fossils except for the focal fossil from the time-heterogeneous pooled tree sample and re-computed the MCC tree. We refer to these as the ‘one-fossil’ trees. Additionally, to explore in detail the different positions in which a given fossil might occur, we listed all the sampled sister groups (single species or clades) in the full (unpruned) time heterogenous tree sample and recorded the frequency of each sister group. Since tree topologies are sampled proportionally to their posterior probability (PP) by the MCMC, we computed the posterior probability of each sister group by dividing the frequency of each sister group by the total number of tree samples in the pooled tree posterior, and plotted them as histograms for visualization.

### Fossil-extant summary tree

After examining the certainty of the fossil placement, we identified five fossils whose placement was extremely uncertain, i.e., not only the PP of the most likely sister group was very low, but also, many distantly related sister groups had similar values of PP. These fossils with particularly uncertain placement were *Capsicum pliocenicum* Deanna & S.D.Sm., *Solanum foveolatum* Negru, *Nephrosemen reticulatum* Manchester, *Solanispermum reniforme* M.Chandler, and *Albionites arnensis* (M.Chandler) Deanna & S.Knapp. We removed these five fossils with particularly uncertain placement from each tree in the time-heterogeneous pooled posterior tree sample, and computed a final extinct plus extant MCC tree. Effectively, this procedure averages the node ages over the uncertainty of the placement of all fossils, while facilitating the interpretation of figures by eliminating the branch instability that results from the high uncertainty in the placement of these five fossils.

### Phylogenetic signal and ancestral state reconstructions

In order to assess the phylogenetic structure in the morphological data, we estimated Blomberg’s K (Blomberg et al. 2003) for the continuous variables and Fritz and Purvis’s D (FPD, Fritz and Purvis, 2010) for the binary variables. We carried out 1000 simulations (for K) and 1000 permutations (for FPD) to test for significant phylogenetic signal on each of 100 trees sampled from the posterior distribution from the pooled time heterogeneous runs. Given the requirement of ultrametric trees for these metrics, we pruned these trees to only the extant tips. These analyses were carried out using the phytools package 2.1-1 (Revell, 2024) and caper 1.0.3 (Orme *et al*., 2023) packages for R 4.5.0 (R Core Team, 2024).

To explore the evolutionary history of morphological traits, we performed ancestral state reconstructions on the same extant-only MCC tree. Eight continuous traits were mapped using the contMap function from the phytools package (Revell, 2024), while the 17 categorical characters were reconstructed using the make.simmap and describe.simmap functions from phytools package (Revell, 2024). Bayesian stochastic character mapping was conducted under the best-fitting model of trait evolution (either all-rates-different or equal-rates) (Huelsenbeck *et al*., 2003; Nielsen, 2001), with 1,000 simulations of character history carried out on the MCC tree in R version 4.5.0 (R Core Team, 2024).

### Reproducibility

All the post-processing and figures described in this section were performed in R (R Core Team, 2024), using the following packages: ape, RevGadgets (Tribble *et al*., 2022), phangorn (Schliep, 2011), deeptime (Gearty, 2024), tidyr (Wickham *et al*., 2024), ggplot2 (Wickham, 2016), dplyr (Wickham *et al*., 2023), and rlist (Ren, 2021). The data and code necessary to run all the analyses described above is available in the GitHub repository https://github.com/ixchelgzlzr/solanaceae_repo.

## RESULTS

### Phylogenetic relationships excluding time

Here and below, we will describe relationships using Solanaceae clade names following the Phylocode (Deanna *et al*., 2025b); these rank-free taxon names are indicated with “/” before the name (Cantino and de Queiroz, 2020). Our initial runs including the three types of data (DNA sequences, categorical, and continuous traits) but ignoring age estimation resulted in topologies similar to expected relationships based on previous family-level studies (Fig. S2). This ‘no-time’ MCC tree contains all of the major clades (e.g., the X=12 clade /Dodecachroma, the berry clade /Solanoideae), although many of these have low support because of the shifting placement of many of the fossils (e.g., *Solanoides dorofeevii*). Nevertheless, several of the fossils fell into positions expected based on morphology (e.g., the *Lycianthoides* and *Physalis* fruit fossils with extant *Lycianthes* and extant *Physalis*, respectively, and the *Hyoscyamus undulatus* seed fossil with the extant *Hyoscyamus*), and for the more character-rich fruit fossils, these placements were relatively well-supported (PP 0.62-0.88, Fig. S2). These findings formed the basis for our two topological constraints (monophyly of *Physalis* and monophyly of *Hyoscyamus*) used to aid the topological convergence of the full analysis.

### Model convergence and comparison

Examination of the trace plots for the two independent MCMC chains for the two models (time-homogeneous and time-heterogeneous) suggested adequate mixing and convergence on the posterior distribution. The two independent chains of 8,000 steps for each model searched similar regions of tree space (Fig. S3), and across models, we recovered very similar maximum clade credibility trees for extant tips (Fig. S4) in terms of both the estimated relationships and their posterior probability. Indeed, only four nodes differed between the two trees (Fig. S5). We also examined the dates for individual nodes of interest and found broadly overlapping estimates for deep and shallow nodes (Fig. S6). Considering the similarity in the estimated trees and dates across the models, we focus on the results from the time-heterogeneous model. We expect that a time heterogeneous model is more adequate for this group given its age (> 90Ma), and may give some additional insight into variation in diversification dynamics over time.

### Phylogenetic signal in morphological data

The majority of the 25 scored morphological characters showed detectable phylogenetic structure based on the estimated relationships. Most of the binary variables (10 out of 14) exhibited significant phylogenetic signal, with the probability of arising under a random (non-phylogenetic) distribution being less than 0.05 across all the 100 subsampled trees (Table S5). The strongest signal among the discrete traits was inferred for calyx teeth, which are only found in a single extant clade (*Lycianthes* plus *Capsicum*) and the fossil fruit *Lycianthes calycina*. We also found a strong and consistent signal for all the continuous variables, with average P-values rejecting K equal to zero of 0.001 to 0.036 across the 100 trees (Table S4). Ancestral state reconstructions for both categorical and continuous traits reveal clear clustering of character states, for example with very large seeds concentrated in /Goetzeoideae and allied genera and flattened seeds only occurring in the /Solanoideae (Figs. S7 and S8). The characters lacking strong phylogenetic signal (Table S5) were rare across the tips and/or widely distributed across the phylogeny (e.g., seed wings, fruiting calyx base invagination). Collectively, these results support the informativeness of the morphological data as observed in the ‘no-time’ analysis described above.

### Phylogenetic relationships across the family

The extant-only MCC tree from the time-heterogeneous analysis shows strong support for well-established clades, along with surprisingly high support for contentious branches within /Physalideae. The X=12 clade (/Dodecachroma, Deanna *et al*., 2025b) and the two component clades (the tobaccos and allies, /Nicotianoideae, and the berries, /Solanoideae) all are inferred with 100% posterior probability (PP) (Fig. 1). Within the berries, the chillipepper clade (/Capsiceae) is sister to the tomatillo clade (/Physalideae) with 100% PP. Backbone relationships within the tomatillos receive much stronger support than in previous studies (Deanna *et al*., 2019) and place *Withania* and allied genera sister to *Physalis* and allies (Fig. 1). /Physalideae are the best represented berry lineage in terms of the fossil record per the relationships estimated here (three fossil fruits and one fossil seed, Fig. 2), which may have contributed to the differential support for possible backbone relationships.

**Figure 1.**
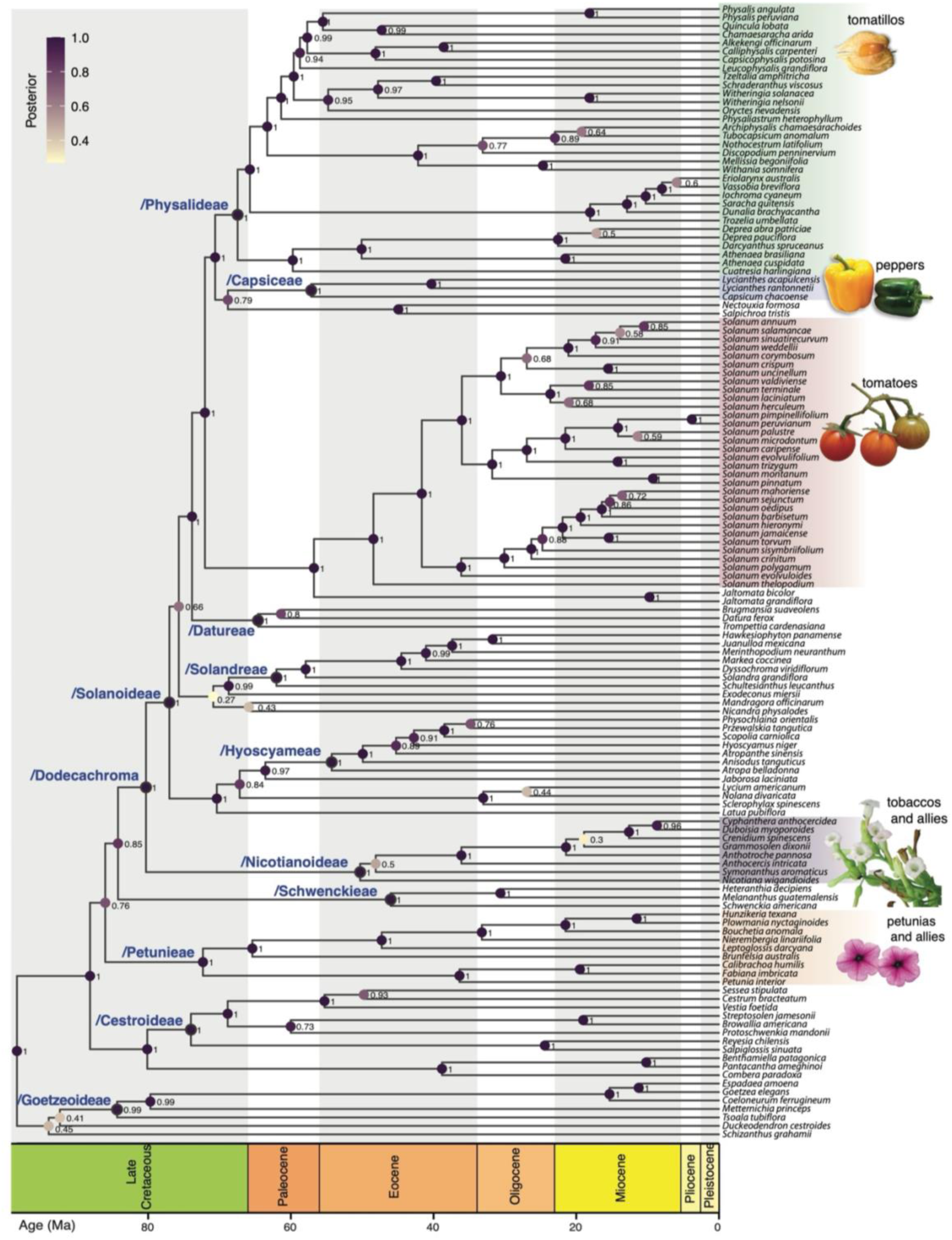
Maximum clade credibility time-calibrated tree for extant Solanaceae inferred under a time-heterogeneous model. Numbers below branches denote posterior probability. Clades are named according to the phylocode nomenclature for Solanaceae (Deanna *et al*., 2025b). Image attributions: Inat observations *Nicotiana tabaccum* #18460677 and *Solanum lycopersicum* #80608627 by user leaf0505 under CC0 license.

**Figure 2.**
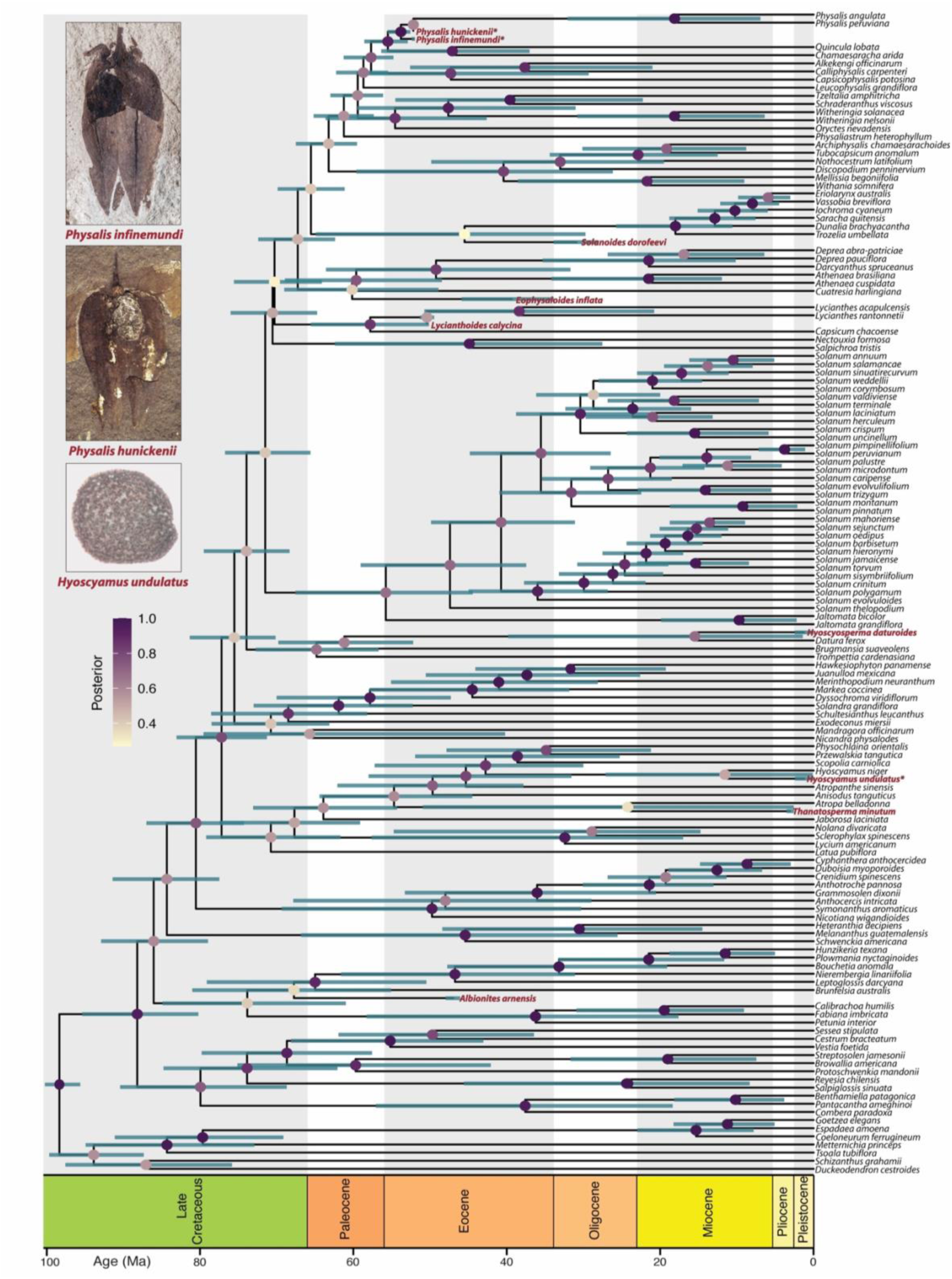
Maximum clade credibility time-calibrated tree for Solanaceae including extinct and extant tips. This tree summarizes the total posterior sample of trees, which includes 13,902 topologies obtained from two convergent and mixing runs under the time-heterogeneous model. The six fossils with particularly uncertain placement are not included in this figure. For each node, the color of the circles represents the PP value, and the bars represent the 95% HPD age. Fossils inferred as sampled ancestors are marked with a black circle around the node they occupy, and the probability displayed represents the PP of them not being a sampled ancestor (as this is the output format of RevBayes for sampled ancestors). The fossils displayed in this figure are the three fossils whose position was constrained; images adapted from Deanna *et al*. (2020, 2023, 2025b).

Nevertheless, the topology is uncertain for many regions of the tree that have been challenging in previous studies as well. For example, the small genus *Nicandra* (three species) and the monotypic *Mandragora* belong to the berry clade, but there is only 27% PP for their position as the sister group to *Exodeconus* + /Solandreae in the extant-only MCC tree (Fig. 1). Several of the earliest splits are also poorly supported. In particular, the MCC tree suggests a basal split between /Goetzeoideae + *Schizanthus* + *Duckeodendron* and the rest of the family, but the support for those taxa forming a clade is only 45%. Within that clade, the position of the enigmatic monotypic *Duckeodendron* is also equivocal, with only 42% support for a sister group relationship with /Goetzeoideae.

### Fossil placement

Summary trees including all the fossils simultaneously result in MCC trees with poor node support due to the particularly uncertain placement of a few fossils (Fig. S9). Our scrutiny of the ‘one-fossil’ tree traces and the distribution of PP for sister groups revealed wide variation in the degree of certainty in fossil placement (Fig. 3, Table S6). Among our 14 fossil taxa, three were used in topological constraints, leaving 11 fossils whose position was freely estimated by the MCMC. The highest probability nodes comprised the sister group relationships of the seed fossil *Hyoscyosperma datureoides* to the extant *Datura* and of the fruit fossil *Lycianthoides calycina* to extant *Lycianthes* (retrieved either as sister to one of the two extant representatives included in this work, or most likely as a sampled ancestor of both extant *Lycianthes*, as shown in Fig. 2) within the /Capsiceae (Figs. 2, 3). In both cases, these fossils shared unique features with those extant taxa (the elaiosomes of *Datura* and the calyx teeth of /Capsiceae). The fossil berry *Eophysalodes inflata* was estimated to belong to the tomatillo clade /Physalideae, which presents ca. 25 origins of similar inflated fruiting calyces among the extant taxa (Deanna *et al*., 2019). Nevertheless, this feature is found among other taxa (e.g., *Nicandra*, *Atropanthe*, *Anisodus*), suggesting that other characters (e.g., venation patterns, calyx size) likely contributed to its placement within /Physalideae. Notably, all of these most confidently placed fruit and seed fossils fall within the berry clade.

**Figure 3.**
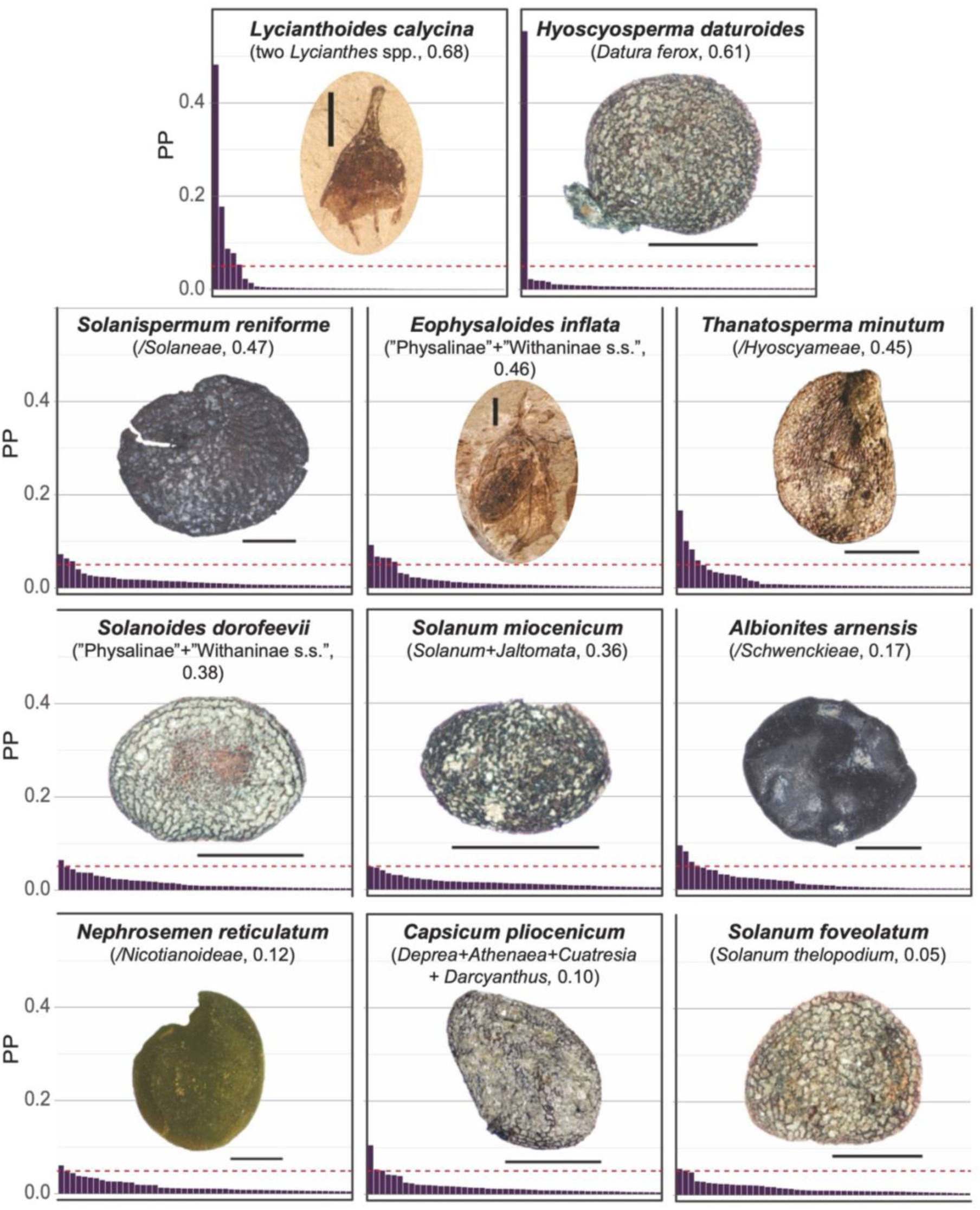
Variation in confidence of fossil position for the 11 topologically unconstrained fossils. The most probable sister group and the posterior probability for the subtending node from ‘one-fossil’ MCC trees (Fig. S10) is given in parentheses below the fossil name. The histograms show the distribution of posterior probabilities for the 50 most sampled sister groups in the ‘one fossil at a time’ tree traces (see also Table S6). The red dashed line indicates PP of 0.05.

There was less certainty in the precise placement of the rest of the fossils, which are all seeds, but in many cases, the sampled sister groups suggest their inclusion in major clades. For example, while the support for any single sister group of *Thanatosperma minutum* was less than 0.2 PP (Fig. 3), many of these fall in /Hyoscyameae (Table S6), which appears as the most probable sister group in its one-fossil tree (PP = 0.45, Fig. 3). Similarly, most probable sister groups for *Solanoides dorofeevi* largely correspond to different lineages with the tomatillo clade /Physalideae (Table S6), consistent with the presence of features commonly found in berry seeds (e.g., lateral compression, lateral hilum with a small cavity). Most of the other seeds (*Capsicum pliocenicum*, *Solanum foveolatum, Solanum miocenicum* and *Solanispermum reniforme*) are likely to be berry seeds also (Fig. 3, Table S6), but their sampled sister groups (Table S6) are spread widely across /Solanoideae. The two fossil seeds lacking strong lateral compression (the globose fossil seeds *Albionites arnensis* and *Nephrosemen reticulatum*) have their most probable placements in capsule-fruited clades (/Schwenckieae and /Nicotianoideae, respectively, Fig. 3), which also present unflattened seeds (e.g., globose or polyhedral), and their most probable sister groups do not include berry lineages (Table S6). Nevertheless, those placements in the one-fossil trees were among the lowest (PPs of 0.17 and 0.12, respectively). Ultimately, we chose to retain five fossil seeds (*S. dorofeevi*, *H. datureoides*, *A. arnensis, T. minuta,* and *H. undulatus* (a constraint)) in the extant-extinct summary tree (Fig. 2), as the position of these fossils suggests they belong to particular major lineages. These affinities observed in the one-fossil analyses were retained in the summary tree including nine fossils, with the exception of *A. arnenis*, which finds its most probable sister group in another capsule clade, /Petunieae (Fig. 2).

### Timing and patterns of diversification

Our time-heterogenous model places the crown age of the family at 98 Ma (95% HPD: 95.66-100.27 Ma) with many major splits clustered within the late Cretaceous (Fig. 2). Specifically all of the lineages previously recognized as subfamilies (/Goetzeoideae, /Cestroideae, /Nicotianoideae, /Solanoideae) appeared by ca. 70 Ma, although /Nicotianoideae did not begin to diversify until the Eocene (50 Ma, 95% HPD: 31.13-70.60 Ma). This time coincides closely with the crown age of *Solanum* (48.43 Ma, 95% HPD: 37.90-59.38 Ma, Fig. S11). Our analyses suggest that the tomatillo and chili pepper lineages began to diversify before *Solanum* with crown ages of 67.46 Ma (95% HPD: 62.70-72.59 Ma) for /Physalideae and 57.14 Ma (95% HPD: 48.51-67.53 Ma) for /Capsiceae, consistent with the appearance of fossils for these lineages by the early Eocene (Fig. 2). The mean age estimates and 95% HPD for each node are detailed in Fig. S11.

The diversification and fossilization rates estimated for Solanaceae from 100 Ma to the present suggest relatively stable dynamics over time (Fig. 4). Fossilization rates were estimated to be slightly higher in the Eocene (56 to 33.9 Ma), when all of the fossil fruits appeared. Diversification rates showed a slight decrease in the Eocene and Oligocene (33.9 Ma to 23.03 Ma) followed by an uptick in the Miocene (Fig. 4). This time period corresponds to the emergence of genera within /Nicotianoideae and /Physalideae as well as major lineages within /Solanum (Fig. 2). It is worth noting that these diversification dynamics are closely tied to the taxonomic sampling for our study, which is focused on major clades and genera.

**Figure 4.**
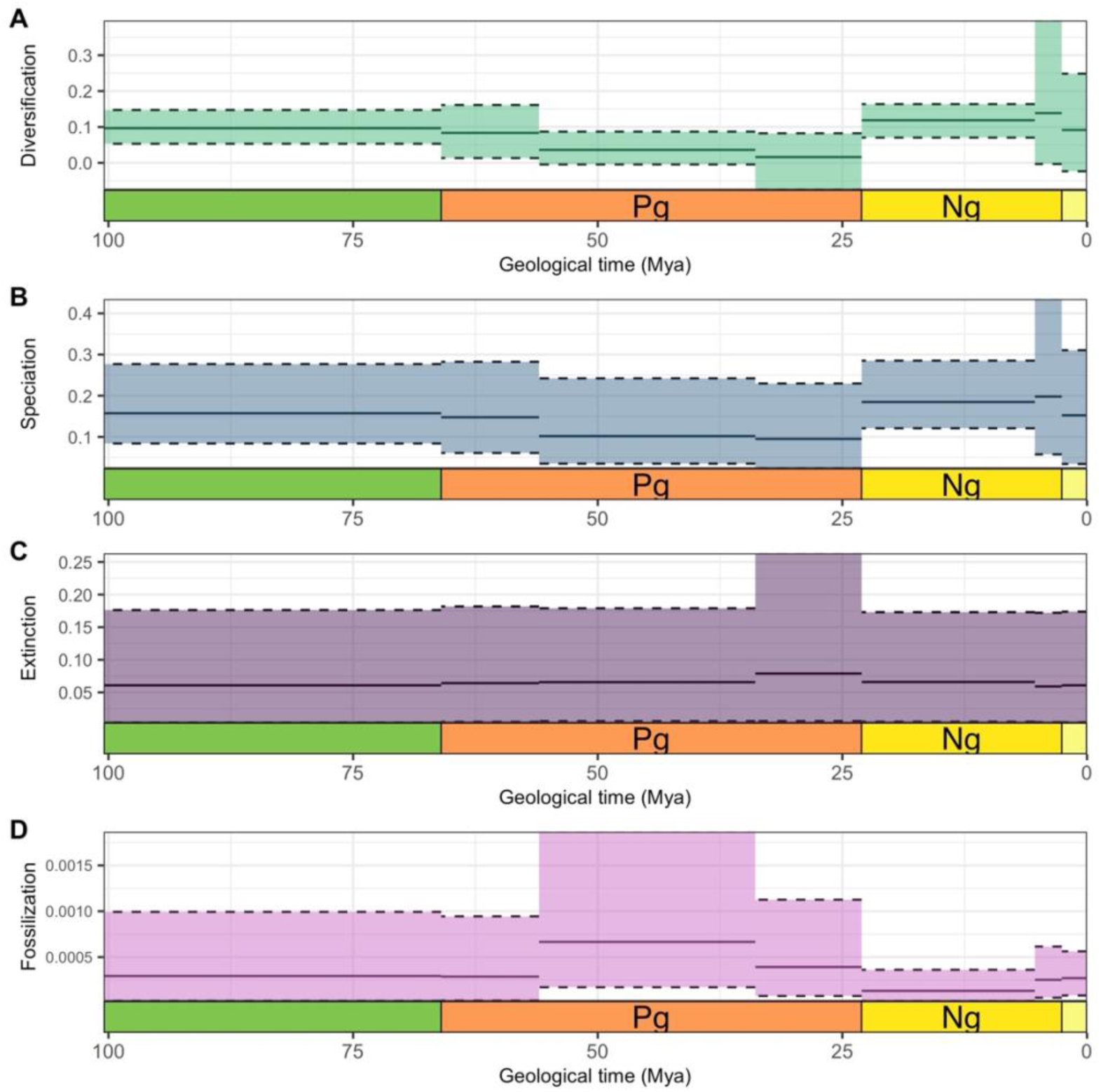
Inferred diversification, speciation, extinction, and fossilization rates through time for Solanaceae under the time-heterogeneous model. The intervals reflect the 95% HPD for each parameter.

## DISCUSSION

### Timeline of nightshade evolution

As one of the most economically important angiosperm families, understanding the timing of Solanaceae diversification of the family has been a long-standing goal. A time-calibrated phylogeny is essential not only for basic research (e.g., of biogeography, morphology, development) but a wide range of applied areas (e.g., plant breeding, genomics, natural products chemistry). Here we estimated that many diverse groups of Solanaceae once thought to be recent radiations, are instead much older, some with roots in the Cretaceous. For example, previous work estimated that chilipeppers and tomatoes diverged just 19 million years ago (Sarkinen *et al*., 2013), which would imply a diversification rate of roughly 0.4 new species per lineage per million years to produce the extant species richness (ca. 1850 spp.), much higher than the estimated rate of 0.1 across all angiosperms (Magallón and Castillo, 2009). Our analyses place the age for that common ancestor at 72MA (95% HPD: 67-77 Ma; Figs. 2 and S11), providing a much longer window for diversification. Similarly, the split between the two lineages comprising /Dodecachroma, namely, the berries (/Solanoideae) and the tobaccos and allies (/Nicotianoideae) has been suggested to be as recent as ca. 25Ma, in both node-dating analyses (Sarkinen *et al*., 2013) and analyses based on molecular substitution rates alone (Wang *et al*., 2024). Our total evidence dating estimates this common ancestor at 80Ma (95% HPD: 73.75-86.57, Figs. 2 and S11). Although most of our sampling was at the genus-level, they imply that previously estimated divergence time within genera (e.g. *Solanum*, Echeverría-Londoño *et al*., 2020, and *Sclerophylax*, Chiarini *et al*., 2022) are also likely much older.

These differences in the inferred dates are due to the sources of information and the analytical methods. First and foremost, the present study made use of a greatly expanded fossil record resulting from nearly a decade of Solanaceae-focused paleobotanical research (Wilf *et al*., 2017; Särkinen *et al*., 2018; Deanna *et al*., 2020, 2023, 2025a; Franco and Brea, 2023). Among these taxa, the fruit macrofossils provide a wealth of characters that can be used for phylogenetic inference (Deanna *et al*., 2023). Second, we applied a fully Bayesian approach in which models were developed for each type of character and integrated into the estimate of topology and branch lengths (Fig. S1). With this total evidence approach, we were able to leverage the morphological variation in the fossils to estimate their position in the phylogeny, as opposed to fixing their placement a priori. Finally, we benefit from recent studies at the angiosperm-level that have clarified the ages of families (Ramirez Barahona *et al*., 2020; Zuntini *et al*., 2024) and could be used to select more appropriate constraints for the origin of the Solanaceae. In particular, these studies suggested that the stem age for the family traces to the Cretacous, consistent with the older fossils recently discovered in Solanaceae and used here for total evidence dating.

Deeper estimates for the age of Solanaceae carry significant implications for the diversification of the family across time and space. With its center of diversity in northern South America (Gentry, 1982), the evolutionary history in the family has often been tied to the phases of rapid northern Andean uplift during the Miocene (e.g., Dillon *et al*., 2009; Chiarini *et al*., 2017; Tovar *et al*., 2021) and this history, in turn, has been used to calibrate the radiations of specialized herbivores (De-Silva *et al*., 2017; Chazot *et al*., 2018). Our model-based and total-evidence results provide strong support for the early diversification of the family, beginning in the mid to late Cretaceous and giving rise to all of the major lineages, from petunias, to tobaccos, to peppers and tomatillos. During this time, the uplift of the southern Andes had only just begun, and much of the northwestern edge of South America was below-sea level (Boschman, 2021). Moreover, South America remained in relatively close proximity to Africa and was still connected to Australia via Antarctica, facilitating the dispersal of plants and other taxa around the southern Hemisphere (Hill & Scriven, 1995; Sanmartin and Ronquist, 2004; Poropat, 2016; Ruhfel *et al*., 2016). This geological context may help to explain the wide geographic range among major Solanaceae lineages, with many lineages split between South America and Australasia (e.g., *Lycianthes*, Knapp, 2022; /Nicotianoideae, Clarkson *et al*., 2004). The coincidence of their splits with the late Cretacous break-up of Gondwana opens the possibility for a larger role for vicariance in Solanaceae biogeography than has been previously considered (Olmstead, 2013; Dupin *et al*., 2017).

### Incorporating fossils as tips in model-based divergence time estimation

In the same way that extant tips are samples of an evolutionary history, fossils are also samples of the history of a lineage, with the particularity that they provide unique information regarding the timing of events. Estimating fossils’ phylogenetic relationships within a model that treats fossils as samples is the most coherent and biologically realistic approach for timetree analysis (Heath *et al*., 2014). Still, even the most character-rich fossil taxa provide limited information for phylogenetic inference compared to extant taxa for which we have sequence data and a morphological description of the entire organism. Thus, we expect a much larger degree of uncertainty in the phylogenetic position of fossil taxa. Taking a Bayesian approach to co-infer divergence times and topologies is ideal to account for the effect of topological uncertainty in age estimates. Nevertheless, the combined effect of multiple fossils topological uncertainty poses a challenge when summarizing tree topology. Notably, when we compute a MCC tree including all the fossils at a time, we obtain negative branch lengths for a few nodes, those that subtend distantly related clades between which a couple fossils “jump” (Fig. S9). Furthermore, we observe large drops in the PP of clades that are highly supported when including only for the extant tips, e.g., compare the posterior probability along the backbone of the berry clade between the extant-only tree (Fig. 1) and the tree containing fossils (Fig. 2). In other contexts, tips found in contrasting positions are often referred to as ‘rogue taxa’ and a common approach that researchers take is to eliminate them from the analyses, at least in part because they might be interested in point estimates of topology. While point estimates are useful in the situation in which data is abundant, they might lead to biased results and interpretations of datasets in which we expect considerable uncertainty in our estimates. In this analysis, we are interested in leveraging the age information of all fossils (by including all of them in the analyses) while also comprehending the extent to which we can make inferences about fossil placement (by carefully examining our posterior sample of trees).

We argue that the best way to understand fossil placement is to study the posterior distribution of trees *one-fossil-at-a-time*. The combination of looking at ‘one-fossil’ trees (Figs. S10) and observing the posterior probability distribution of unique sampled sister groups (Fig. 3, Table S6) allowed us to identify the highest probability sister groups for each fossil without the confounding effect of other fossils. We noted that the maximum PP was not necessarily predictive of the concentration of the distribution around a set of possible sister groups. For example, while the fruit fossil *Lycianthoides calycina* has the best supported placement as a member of the *Lycianthes* stem lineage in its one-fossil tree (Fig. S10, PP=0.68), the analysis explored five other placements within /Capsiceae with relatively high PP, including positions as sister to *Capsicum* (Fig. 3, Table S6). By contrast, while *Hyoscyosperma daturoides* has lower PP for its sister group relationship with *Datura* in its one-fossil tree (PP=0.61, Fig. S10), the posterior probabilities for other possible sister groups are negligible (0.00007-0.019). With this detailed picture of each fossil’s behavior in the analysis, we elected to remove five of the original 14 fossils in each tree sampled in the posterior, and use the remainder for computing the best point estimate of the timetree (Fig. 3).

These results provide new insight into the evolutionary history of Solanaceae fossil taxa and their suites of characters. Parsimony-based analyses previously placed *Lycianthoides calycina* in /Capsiceae, which is the only Solanaceae lineage presenting the finger-like calyx appendages instead of calyx lobes, and our analysis strongly supports its placement in /Capsiceae (PP = 1.0, Table S6). As with the unique /Capsiceae calyx teeth, our phylogeny points to a single origin for elaiosomes, the food bodies involved in ant dispersal of seeds (Gorb and Gorb, 2003). The *Hyoscyosperma daturoides* fossil was originally placed in *Physalis* (Dorofeev, 1957) although its small size and sinuate-cerebelloid anticlinal walls of its exotestal cells on the entire seed surface make it more similar to *Hyoscyamus* (Deanna *et al*., 2025a). However, these morphological features have apparently arisen multiple times in the family (Fig. S8), while elaiosomes are only present in *Datura*. The fossil berry *Eophysaloides inflata* also presents a unique combination of features in terms of its calyx venation, lack of calyx invagination at the base and berry loosely included in the calyx, characters shared with *Calliphysalis* and *Brachistus* (now subsumed under *Witheringia*; Stone *et al*., 2024), although its name translates to ‘early’ ‘physaloid’ due to its overall similarity to other members of /Physalideae (Deanna *et al*., 2023). Our results support the conclusion that *Eophysaloides* belongs to /Physalideae although its position within the clade is less clear (Figs. 2, 3, S10, and Table S6). While the relationships of the seed fossils, e.g. *Solanum foveolatum*, *Capsicum pliocenicum*, to extant taxa has been more difficult to discern (Deanna *et al*., 2025a), and our analysis provides a fair support for the placement of some of the seeds, e.g.,*Thanatosperma minutum* (Figs. 2, 3, S10, and Table S6).

Furthermore, we emphasize that a total-evidence dating approach not only provides a framework for understanding the relationships of fossils to other lineages while accounting for the uncertainty in their placement, in fact, it is the expected uncertainty in the placement of some fossils which makes a Bayesian approach ideal to average over the topological uncertainty when inferring divergence times.

### Implementing the total evidence dating of plant clades with the fossilized birth-death model

Applications of the FBD model to total evidence dating are overwhelmingly concentrated in animal lineages, with only ca. 15% of studies to date focused on plants (Mulvey *et al*., 2025). The pattern may relate to greater challenge in obtaining fossil information, e.g. due to the lower tendency for plant tissues to be preserved compared to shells or bones and the separation of whole plants into their component organs (Locatelli, 2014). Furthermore, studies of plant fossils often focus on a single type of fossil that is best represented in the fossil record, e.g. pollen (Bansal *et al*., 2021; Woutersen *et al*., 2023), fronds (Grimm *et al*., 2015), or fruits (Zhang *et al*., 2022), but all tips need to be scored for the same set of characters in a total-evidence analysis. In order to apply the FBD model for total-evidence dating in Solanaceae, we led a concerted effort to revise fossil material globally and describe new fossil taxa for the most common type of material (seeds) as well as the most character-rich material (fruits) (Deanna *et al*., 2020, 2023, 2025a). Thus, for plant clades with a sparse fossil record scattered across different types of organs, total evidence dating may need to begin with targeted revisions of the rich and understudied paleobotanical collections worldwide.

The Bayesian approach also provides an explicit framework for incorporating prior information to generate biologically realistic estimates. Prior studies have carried out in-depth morphological analyses of each of the fossil taxa and in cases where our preliminary ‘no-time’ analyses confirmed the placement of fossils within extant genera (Fig. S2), we employed topological constraints to guide the broader analysis. We also recognize that morphological characters evolve at different rates, just like sites in DNA sequences, and applied an analogous approach using the gamma distribution to model that variation (Fig. S1). This experimental design choice may have allowed the extremely phylogenetically restricted characters (calyx teeth, elaisomes) to fall into the slow-rate category and make grouping species with those traits more probable. Indeed, our estimates indicate a difference of three orders of magnitude (0.07 vs 2.6) between the slowest and highest rate categories for the model of categorical trait evolution. We allowed for similar variation in rates of continuous trait evolution, as we expect, for example, that size may evolve faster than shape (Liu and Smith, 2023). While these analytical choices were tailored for our taxa and our characters, we suspect that the tools employed here (incorporation of a small set of topological constraints; comparison of analyses with and without time; one-by-one examination of fossil placement) may be the key to successfully scaling fully model-based divergence time estimation to larger datasets with greater numbers of both fossils and extant taxa.

## CONCLUSIONS

Evolutionary inferences rely on an understanding of the absolute timescale, which in turn is best understood from the fossil record (Forest, 2009). For over a decade, basic and applied research on Solanaceae has relied on dates for major splits that are roughly three-fold younger than the dates estimated here. These dates, in turn, have been used to estimate many aspects of the family’s biology, from rates of molecular evolution (e.g., Kim *et al*., 2014; Cao *et al*., 2021), to the timing of polyploid events (Clarkson *et al*., 2005; Sato *et al*., 2012), to the biogeographic history (Dupin *et al*., 2017) and patterns of diversification (Echeverría-Londoño *et al*., 2020). While some inferences may still hold even under deeper dates, we expect that many of the conclusions will need to be revisited. Moving forward, our timetree may serve as a benchmark to calibrate more densely sampled phylogenies, for which implementing a similar analysis would be impractical. For example, our results did not recover any fossils that fall inside of the large genus *Solanum* (ca. 1240 spp.), and even with scoring additional extant taxa, there may simply not be enough information in some of the seed fossils to achieve a more refined estimate of divergence times within the clade. In this sense, we highlight the importance of paleobotanical exploration as the main source of new information with potential to better our understanding of the timing of clades like *Solanum*. However, our study gives conservative estimate for the crown age of *Solanum* (48.43 Ma, 95% HPD: 37.90-59.38) that could be used as a secondary calibration in subsequent studies of the genus. Indeed, expanding the taxon sampling within individual clades will be crucial for tracing the diversification of Solanaceae over the past 100 million years and identifying the events that could explain the incredible dominance of nightshade berries around the globe.

## Supporting information

Supplementary Materials including all referenced supplementary figures and tables.

## Author Contributions

SDS and RD conceived the study. RD and SDS assembled the data matrices. IGR designed and implemented the statistical analyses. IGR, RD, and SDS wrote the original manuscript and edited it.

## Acknowledgements.

We thank Michael May and Ben Redelings for their help with various questions regarding the implementation of the analyses.

## REFERENCES

Álvarez-Carretero S, Goswami A, Yang Z, Dos Reis M. 2019. Bayesian estimation of species divergence times using correlated quantitative characters. Systematic Biology 68(6): 967–986.

Bansal M, Nagaraju SK, Mishra AK, Selvaraj J, Patnaik R, Prasad V. 2021. Fossil pollen from early Palaeogene sediments in western India provides phylogenetic insights into divergence history and pollen character evolution in the pantropical family Ebenaceae. Botanical Journal of the Linnean Society 197(2): 147–169.

Barba-Montoya J, Dos Reis M, Schneider H, Donoghue PCJ, Yang Z. 2018. Constraining uncertainty in the timescale of angiosperm evolution and the veracity of a Cretaceous Terrestrial Revolution. New Phytologist 218(2): 819–834.

Barido-Sottani J, Van Tiel NM, Hopkins MJ, Wright DF, Stadler T, Warnock RC. 2020. Ignoring fossil age uncertainty leads to inaccurate topology and divergence time estimates in time-calibrated tree inference. Frontiers in Ecology and Evolution 8: 183.

Blaimer BB, Santos BF, Cruaud A, et al. 2023. Key innovations and the diversification of Hymenoptera. Nature Communications 14: 1212.

Boschman LM. 2021. Andean mountain building since the Late Cretaceous: A paleoelevation reconstruction. Earth-Science Reviews 220: 103640.

Brightly WH, Crifò C, Gallaher TJ, Hermans R, Lavin S, Lowe AJ, Smythies CA, Stiles E, Wilson Deibel P, Strömberg CAE. 2024. Palms of the past: can morphometric phytolith analysis inform deep time evolution and palaeoecology of Arecaceae? Annals of Botany 134(2): 263–282.

Cantino PD, De Queiroz K, eds. 2020. PhyloCode: A phylogenetic code of biological nomenclature. Boca Raton: CRC Press.

Cao YL, Li YL, Fan YF, et al. 2021. Wolfberry genomes and the evolution of *Lycium* (Solanaceae). Communications Biology 4: 671.

Chazot N, De-Silva DL, Willmott KR, Freitas AVL, Lamas G, Mallet J, Giraldo CE, Uribe S, Elias M. 2018. Contrasting patterns of Andean diversification among three diverse clades of Neotropical clearwing butterflies. Ecology and Evolution 8(8): 3965–3982.

Coiro M, Allio R, Mazet N, Seyfullah LJ, Condamine FL. 2023. Reconciling fossils with phylogenies reveals the origin and macroevolutionary processes explaining the global cycad biodiversity. New Phytologist 240: 1616–1635.

Crisp MD, Hardy NB, Cook LG. 2014. Clock model makes a large difference to age estimates of long-stemmed clades with no internal calibration: a test using Australian grasstrees. BMC Evolutionary Biology 14: 263.

Chiarini F, Moreno N, Moré M, Barboza G. 2017. Chromosomal changes and recent diversification in the Andean genus *Jaborosa* (Solanaceae). Botanical Journal of the Linnean Society 183(1): 57–74.

Clarkson JJ, Knapp S, Garcia VF, Olmstead RG, Leitch AR, Chase MW. 2004. Phylogenetic relationships in *Nicotiana* (Solanaceae) inferred from multiple plastid DNA regions. Molecular Phylogenetics and Evolution 33(1): 75–90.

Clarkson JJ, Lim KY, Kovarik A, Chase MW, Knapp S, Leitch AR. 2005. Long-term genome diploidization in allopolyploid *Nicotiana* section Repandae (Solanaceae) New Phytologist 168(1): 241–252.

De-Silva LD, Mota LL, Chazot N, Mallarino R, Silva-Brandão KL, Piñerez LMG, et al. 2017. North Andean origin and diversification of the largest Ithomiine butterfly genus. Scientific Reports 7(1): 45966.

Deanna R, Barboza GE, Bosch L, Dodsworth S, Gagnon E, Giacomin LL, Knapp S, Orejuela A, Poczai P, Särkinen T, Smith SD, Olmstead RD. 2025. A new phylogeny and phylogenetic classification for Solanaceae. Taxon, submitted. *bioRxiv* https://www.biorxiv.org/content/10.1101/2025.07.10.663745v1

Deanna R, Larter MD, Barboza GE, Smith SD. 2019. Repeated evolution of a morphological novelty: a phylogenetic analysis of the inflated fruiting calyx in the Physalideae tribe (Solanaceae). American Journal of Botany 106: 270–279.

Deanna R, Wilf P, Gandolfo MA. 2020. New physaloid fruit-fossil species from early Eocene South America. American Journal of Botany 107(12): 1749–1762.

Deanna R, Martínez C, Manchester S, Wilf P, Campos A, Knapp S, Smith SD. 2023. Fossil berries reveal global radiation of the nightshade family by the early Cenozoic. New Phytologist 238(6): 2685– 2697.

Deanna R, Hvalj AV, Martinetto E, Knapp S, Sadowski E-M, Manchester S, Campos A, Fernandez V., Barboza GE, Sauquet H, Dean E, Särkinen T, Chiarini FE, Bernardello G, Smith SD. 2025. Seed fossil record of Solanaceae revisited. Taxon, submitted. *bioRxiv* https://www.biorxiv.org/content/10.1101/2025.07.03.662944v1

Dellinger AS, Lagomarsino L, Michelangeli F, Dullinger S, Smith SD. 2024. The sequential direct and indirect effects of mountain uplift, climatic niche, and floral trait evolution on diversification dynamics in an Andean plant clade. Systematic Biology 73(3): 594–612.

Dillon MO, Tu T, Xie L, Quipuscoa Silvestre V, Wen J. 2009. Biogeographic diversification in *Nolana*(Solanaceae), a ubiquitous member of the Atacama and Peruvian Deserts along the western coast of South America. Journal of Systematics and Evolution 47: 457–476.

Dorofeev PI. 1957a. O pliotsenovoy flore nagavskikh glin na Donu [On the Pliocene flora of the Nagava clays on the Don]. Doklady Akademii Nauk SSSR 117(1): 124–126.

Drummond AJ, Ho SYW, Phillips MJ, Rambaut A. 2006. Relaxed phylogenetics and dating with confidence. PLoS Biology 4(5): e88.

Dupin J, Matzke NJ, Särkinen T, Knapp S, Olmstead RG, Bohs L, Smith SD. 2017. Bayesian estimation of the global biogeographical history of the Solanaceae. Journal of Biogeography 44: 887– 899.

Echeverría-Londoño S, Särkinen T, Fenton IS, Purvis A, Knapp S. 2020. Dynamism and context-dependency in diversification of the megadiverse plant genus *Solanum* (Solanaceae). Journal of Systematics and Evolution 58: 767–782.

Edgar RC. 2004. MUSCLE: multiple sequence alignment with high accuracy and high throughput. Nucleic Acids Research 32: 1792–1797.

Foley NM, Mason VC, Harris AJ, Bredemeyer KR, Damas J, Lewin HA, Eizirik E, Gatesy J, Karlsson EK, Lindblad-Toh K, Zoonomia Consortium, Springer MS, Murphy WJ. 2023. A genomic timescale for placental mammal evolution. Science 380(6643): eabl8189.

Forest F. 2009. Calibrating the Tree of Life: fossils, molecules and evolutionary timescales. Annals of Botany 104(5): 789–794.

Franco MJ, Brea M. 2023. Redescription of *Solanumxylon paranensis*, Late Miocene of Paraná Formation (Entre Ríos, Argentina): A Reliable Fossil Wood of Solanaceae. Ameghiniana 60: 65–77.

Fritz SA, Purvis A. 2010. Selectivity in mammalian extinction risk and threat types: A new measure of phylogenetic signal strength in binary traits. Conservation Biology 24: 1042–1051.

Gagnon E, Hilgenhof R, Orejuela A, McDonnell A, Sablok G, Aubriot X, Giacomin LL, Gouvêa Y, Bragionis T, Stehmann JR, Bohs L, Dodsworth S, Martine C, Poczai P, Knapp S, Särkinen T. 2022. Phylogenomic discordance suggests polytomies along the backbone of the large genus *Solanum* (Solanaceae). American Journal of Botany 109(4): 580–601.

Gearty W. 2024. deeptime: an R package that facilitates highly customizable visualizations of data over geological time intervals. EarthArXiv. DOI: 10.31223/X5841N

Gentry AH. 1982. Neotropical floristic diversity: phytogeographical connections between Central and South America, Pleistocene climatic fluctuations, or an accident of the Andean orogeny? Annals of the Missouri Botanical Garden 69: 557–593.

Gorb E, Gorb S. 2003. Seed dispersal by ants in a deciduous forest ecosystem: mechanisms, strategies, adaptations. Dordrecht: Springer Science & Business Media.

Grimm GW, Kapli P, Bomfleur B, McLoughlin S, Renner SS. 2015. Using more than the oldest fossils: dating Osmundaceae with three Bayesian clock approaches. Systematic Biology 64(3): 396– 405.

Heath TA, Huelsenbeck JP, Stadler T. 2014. The fossilized birth–death process for coherent calibration of divergence-time estimates. Proceedings of the National Academy of Sciences USA 111: E2957–E2966.

Hill RS, Scriven LJ. 1995. The angiosperm-dominated woody vegetation of Antarctica: a review. Review of Palaeobotany and Palynology 86(3–4): 175–198.

Höhna S, Landis MJ, Heath TA, Boussau B, Lartillot N, Moore BR, Huelsenbeck JP, Ronquist F. 2016. RevBayes: Bayesian phylogenetic inference using graphical models and an interactive model-specification language. Systematic Biology 65(4): 726–736.

Huang W, Liu W, Wang X. 2024. The first macrofossil record of parasitic plant flowers from an Eocene Baltic amber. Heliyon 10(23): e40794.

Huang J, Xu W, Zhai J, Guo J, Zhang C, Zhao Y, Zhang L, Martine C, Ma H, Huang C-H. 2023. Nuclear phylogeny and insights into whole-genome duplications and reproductive development of Solanaceae plants. Plant Communications 4: 100595.

Huelsenbeck JP, Nielsen R, Bollback JP. 2003. Stochastic mapping of morphological characters. Systematic Biology 52: 131–158.

Hunziker AT. 2001. Genera Solanacearum: The genera of Solanaceae illustrated, arranged according to a new system. Ruggell: A.R.G. Gantner Verlag.

Kara E, Bardin J, De Franceschi D, Del Rio C. 2024. Fossil endocarps of Menispermaceae from the late Paleocene of Paris Basin, France. Journal of Systematics and Evolution 62(4): 809–828.

Kim S, Park M, Yeom SI, et al. 2014. Genome sequence of the hot pepper provides insights into the evolution of pungency in *Capsicum* species. Nature Genetics 46: 270–278.

Knapp S. 2022. A revision of *Lycianthes* (Solanaceae) in Australia, New Guinea, and the Pacific. PhytoKeys 209: 1–134.

Kumar S, Stecher G, Suleski M, Sanderford M, Sharma S, Tamura K. 2024. MEGA12: molecular evolutionary genetics analysis version 12 for adaptive and green computing. Molecular Biology and Evolution 41: 1–9.

Lee MSY, Soubrier J, Edgecombe GD. 2013. Rates of phenotypic and genomic evolution during the Cambrian Explosion. Current Biology 23(19): 1889–1895.

Lepage T, Bryant D, Philippe H, Lartillot N. 2007. A general comparison of relaxed molecular clock models. Molecular Biology and Evolution 24(12): 2669–2680.

Léveillé-Bourret É, Starr JR, Ford BA, Lemmon EM, Lemmon AR. 2018. Resolving rapid radiations within angiosperm families using anchored phylogenomics. Systematic Biology 67(1): 94–112. DOI: 10.1093/sysbio/syx050

Lewis PO. 2001. A likelihood approach to estimating phylogeny from discrete morphological character data. Systematic Biology 50: 913–925.

Li HT, Yi TS, Gao LM, Ma PF, Zhang T, Yang JB, Gitzendanner MA, et al. 2019. Origin of angiosperms and the puzzle of the Jurassic gap. Nature Plants 5(5): 461–470.

Liu K, Li E, Cui X, et al. 2024. Key innovations and niche variation promoted rapid diversification of the widespread *Juniperus* (Cupressaceae). Communications Biology 7: 1002.

Liu S, Smith SD. 2023. Replicated radiations in the South American marsh pitcher plants (*Heliamphora*) lead to convergent carnivorous trap morphologies. American Journal of Botany 110(10): e16230.

Locatelli ER. 2014. The exceptional preservation of plant fossils: a review of taphonomic pathways and biases in the fossil record. The Paleontological Society Papers 20: 237–258.

Magallón SA. 2004. Dating lineages: molecular and paleontological approaches to the temporal framework of clades. International Journal of Plant Sciences 165: S7–S21.

Magallón S, Castillo A. 2009. Angiosperm diversification through time. American Journal of Botany 96(1):349–365.

Magallón S, Gómez-Acevedo S, Sánchez-Reyes LL, Hernández-Hernández T. 2015. A metacalibrated time-tree documents the early rise of flowering plant phylogenetic diversity. New Phytologist 207(2): 437–453.

May MR, Contreras DL, Sundue MA, Nagalingum NS, Looy CV, Rothfels CJ. 2021. Inferring the total-evidence timescale of marattialean fern evolution in the face of model sensitivity. Systematic Biology 70(6): 1232–1255.

Messeder JVS, Carlo TA, Zhang G, Tovar JD, Arana C, Huang J, Huang CH, Ma H. 2024. A highly resolved nuclear phylogeny uncovers strong phylogenetic conservatism and correlated evolution of fruit color and size in *Solanum* L. New Phytologist 243: 765–780.

Minh BQ, Schmidt HA, Chernomor O, Schrempf D, Woodhams MD, von Haeseler A, Lanfear R. 2020. IQ-TREE 2: New models and efficient methods for phylogenetic inference in the genomic era. Molecular Biology and Evolution 37: 1530–1534.

Mongiardino Koch N, Garwood RJ, Parry LA. 2021. Fossils improve phylogenetic analyses of morphological characters. Proceedings of the Royal Society B 288: 20210044.

Mulvey LP, Nikolic MC, Allen BJ, Heath TA, Warnock RCM. 2025. From fossils to phylogenies: exploring the integration of paleontological data into Bayesian phylogenetic inference. Paleobiology 51(1): 214–236.

Ng J, Smith SD. 2016. Widespread flower color convergence in Solanaceae via alternate biochemical pathways. New Phytologist 209: 407–417. 10.1111/nph.13576

Nielsen R. 2001. Mutations as missing data: Inferences on the ages and distributions of nonsynonymous and synonymous mutations. Genetics 159: 401–411. 10.1093/genetics/159.401

Olmstead RG. 2013. Phylogeny and biogeography in Solanaceae, Verbenaceae and Bignoniaceae: a comparison of continental and intercontinental diversification patterns. Botanical Journal of the Linnean Society 171(1): 80–102.

Olmstead RG, Palmer JD. 1992. A chloroplast DNA phylogeny of the Solanaceae: subfamilial relationships and character evolution. Annals of the Missouri Botanical Garden 79: 346–360.

Olmstead RG, Bohs L, Abdel Migid H, Santiago-Valentin E, Garcia VF, Collier SM. 2008. A molecular phylogeny of the Solanaceae. Taxon 57: 1159–1181.

Orme D, Freckleton R, Thomas G, Petzoldt T, Fritz S, Isaac N, Pearse W. 2023. caper: Comparative Analyses of Phylogenetics and Evolution in R. R package version 1.0.

Parins-Fukuchi CT. 2018. Bayesian placement of fossils on phylogenies using quantitative morphometric data. Evolution 72(9): 1801–1814.

Pagel M. 1994. Detecting correlated evolution on phylogenies: a general method for the comparative analysis of discrete characters. Proceedings of the Royal Society of London B 255: 37–45.

Poropat S, Mannion P, Upchurch P, et al. 2016. New Australian sauropods shed light on Cretaceous dinosaur palaeobiogeography. Scientific Reports 6: 34467.

Powell AF, Zhang J, Hauser D, Vilela JA, Hu A, Gates DJ, Mueller LA, Li FW, Strickler SR, Smith S. 2022. Genome sequence for the blue-flowered Andean shrub *Iochroma cyaneum* reveals extensive discordance across the berry clade of Solanaceae. The Plant Genome 15: e20223.

Pyron RA. 2011. Divergence time estimation using fossils as terminal taxa and the origins of lissamphibia. Systematic Biology 60: 466–481.

R Core Team. 2024. R: A Language and Environment for Statistical Computing. Vienna, Austria: R Foundation for Statistical Computing. https://www.R-project.org/.

Ramírez-Barahona S, Sauquet H, Magallón S. 2020. The delayed and geographically heterogeneous diversification of flowering plant families. Nature Ecology & Evolution 4(9): 1232–1238.

Ramírez S, Gravendeel B, Singer R, et al. 2007. Dating the origin of the Orchidaceae from a fossil orchid with its pollinator. Nature 448: 1042–1045.

Ren K. 2021. rlist: A Toolbox for Non-Tabular Data Manipulation. R package version 0.4.6.2. https://CRAN.R-project.org/package=rlist.

Renner SS, Grimm GW, Kapli P, Denk T. 2016. Species relationships and divergence times in beeches: new insights from the inclusion of 53 young and old fossils in a birth–death clock model. Philosophical Transactions of the Royal Society B 371: 20150135.

Revell LJ. 2024. phytools 2.0: an updated R ecosystem for phylogenetic comparative methods (and other things). PeerJ12: e16505.

Ronquist F, Klopfstein S, Vilhelmsen L, Schulmeister S, Murray DL, Rasnitsyn AP. 2012a. A total-evidence approach to dating with fossils, applied to the early radiation of the Hymenoptera. Systematic Biology 61: 973–999.

Rothfels CJ, Larsson A, Kuo LY, Korall P, Chiou WL, Pryer KM. 2012. Overcoming deep roots, fast rates, and short internodes to resolve the ancient rapid radiation of Eupolypod II ferns. Systematic Biology 61(3): 490.

Ruhfel BR, Bove CP, Philbrick CT, Davis CC. 2016. Dispersal largely explains the Gondwanan distribution of the ancient tropical clusioid plant clade. American Journal of Botany 103: 1117–1128.

Sanmartín I, Ronquist F. 2004. Southern hemisphere biogeography inferred by event-based models: plant versus animal patterns. Systematic Biology 53: 216–243.

Sanderson MJ, Doyle JA. 2001. Sources of error and confidence intervals in estimating the age of angiosperms from rbcL and 18S rDNA data. American Journal of Botany 88(8): 1499–1516.

Särkinen T, Bohs L, Olmstead RG, Knapp S. 2013. A phylogenetic framework for evolutionary study of the nightshades (Solanaceae): a dated 1000-tip tree. BMC Evolutionary Biology 13: 214.

Särkinen T, Kottner S, Stuppy W, Ahmed F, Knapp S. 2018. A new commelinid monocot seed fossil from the early Eocene previously identified as Solanaceae. American Journal of Botany 105(1): 95– 107.

Sato S, Tabata S, Hirakawa H, et al. 2012. The tomato genome sequence provides insights into fleshy fruit evolution. Nature 485 (11119): 635–641.

Schliep KP. 2011. phangorn: phylogenetic analysis in R. Bioinformatics 27(4): 592–593.

Silvestro D, Bacon CD, Ding W, Zhang Q, Donoghue PC, Antonelli A, Xing Y. 2021. Fossil data support a pre-Cretaceous origin of flowering plants. Nature Ecology & Evolution 5(4): 449–457.

Smith SA, Walker JF. 2018. PyPHLAWD: A python tool for phylogenetic dataset construction. Methods in Ecology and Evolution 10: 104–108.

Stadler T. 2010. Sampling-through-time in birth–death trees. Journal of Theoretical Biology 267: 396–404.

Stadler T, Gavryushkina A, Warnock RCM, Drummond AJ, Heath TA. 2018. The fossilized birth-death model for the analysis of stratigraphic range data under different speciation modes. Journal of Theoretical Biology 447: 41–55.

Stone JL, Flores J, Bohs L. 2024. Phylogenetic Relationships of *Brachistus* and *Witheringia* (Solanaceae). Systematic Botany 49(2): 496–506.

Szafer W. 1946. Flora plioceńska z Kroscienka nad Dunajcem [The Pliocene Flora of Kroscienko over Dunajec] Krakow: Polska Akademia umiejtnosci.

Tepe EJ, Anderson GJ, Spooner DM, Bohs L. 2016. Relationships among wild relatives of the tomato, potato, and pepino. Taxon 65(2): 262–276.

Tovar JD, André T, Wahlert GA, Bohs L, Giacomin LL. 2021. Phylogenetics and historical biogeography of *Solanum* section *Brevantherum* (Solanaceae). Molecular Phylogenetics and Evolution 162: 107195.

Tribble CM, Freyman WA, Landis MJ, Lim JY, Barido-Sottani J, Kopperud BT, et al. 2022. RevGadgets: An R package for visualizing Bayesian phylogenetic analyses from RevBayes. Methods in Ecology and Evolution 13(2): 314–323.

Troyer EMK, Evans KM, Goatley CHR, Friedman M, Carnevale G, Nicholas B, Kolmann M, Bemis KE, Arcila D. 2024. Evolutionary innovation accelerates morphological diversification in pufferfishes and their relatives. Evolution 78(11): 1869–1882.

Wilf P, Carvalho MR, Gandolfo MA, Cúneo NR. 2017. Eocene lantern fruits from Gondwanan Patagonia and the early origins of Solanaceae. Science 355(6320): 71–75.

Won H, Renner SS. 2006. Dating dispersal and radiation in the gymnosperm *Gnetum* (Gnetales)— clock calibration when outgroup relationships are uncertain. Systematic Biology 55(4): 610–622.

Wang H, Moore MJ, Soltis PS, Bell CD, Brockington SF, Alexandre R, Davis CC, Latvis M, Manchester SR, Soltis DE. 2009. Rosid radiation and the rapid rise of angiosperm-dominated forests. Proceedings of the National Academy of Sciences USA 106(10): 3853–3858.

Wang J, Zhang Q, Tung J, Zhang X, Liu D, Deng Y, Tian Z, Chen H, Wang T, Yin W, Li B, Lai Z, Dinesh-Kumar SP, Baker B, Li F. 2024. High-quality assembled and annotated genomes of *Nicotiana tabacum* and *Nicotiana benthamiana* reveal chromosome evolution and changes in defense arsenals. Molecular Plant 17(3): 423–437.

Warnock RC, Wright AM. 2020. Understanding the tripartite approach to Bayesian divergence time estimation. Cambridge University Press.

Whitfield JB, Lockhart PJ. 2007. Deciphering ancient rapid radiations. Trends in Ecology & Evolution 22(5): 258–265.

Wickham H. 2016. ggplot2: Elegant Graphics for Data Analysis. Springer-Verlag New York.

Wickham H, François R, Henry L, Müller K, Vaughan D. 2023. dplyr: A Grammar of Data Manipulation. R package version 1.1.4. https://CRAN.R-project.org/package=dplyr.

Wickham H, Vaughan D, Girlich M. 2024. tidyr: Tidy Messy Data. R package version 1.3.1. https://CRAN.R-project.org/package=tidyr.

Woutersen A, Jardine PE, Silvestro D, Bogotá-Angel RG, Zhang HX, Meijer N, Bouchal J, Barbolini N, Dupont-Nivet G, Koutsodendris A, Antonelli A, Hoorn C. 2023. The evolutionary history of the Central Asian steppe-desert taxon *Nitraria* (Nitrariaceae) as revealed by integration of fossil pollen morphology and molecular data. Botanical Journal of the Linnean Society 202(2): 195–214.

Wu Y, Kodrul T, Zheng Y, Maslova N, Ni ZJ, Wu XK, Jin JH. 2025. A naturally folded leaf fossil of *Bauhinia* sl from the middle Paleocene of South China and its phytogeographical and palaeoecological implications. Papers in Palaeontology 11(2): e70013.

Zhang Q, Ree RH, Salamin N, Xing Y, Silvestro D. 2022. Fossil-informed models reveal a boreotropical origin and divergent evolutionary trajectories in the walnut family (Juglandaceae). Systematic Biology 71(1): 242–258.

Zhang R, Drummond AJ, Mendes FK. 2024. Fast Bayesian inference of phylogenies from multiple continuous characters. Systematic Biology 73(1): 102–124.

Zuntini AR, Carruthers T, Maurin O, et al. 2024. Phylogenomics and the rise of the angiosperms. Nature 629: 843–850.

