## Supplementary Materials including all referenced supplementary figures and tables. for "Late Cretaceous origins for major nightshade lineages from total evidence timetree analysis"

**Figure S1**. Graphical representation of the total-evidence dating model used for Bayesian time tree estimation. Each of the three data types is assigned a different model of evolution (see methods). The clock models are linked to the discrete morphological characters and the sequence data. The tree model includes rates of speciation, extinction, and fossilization.

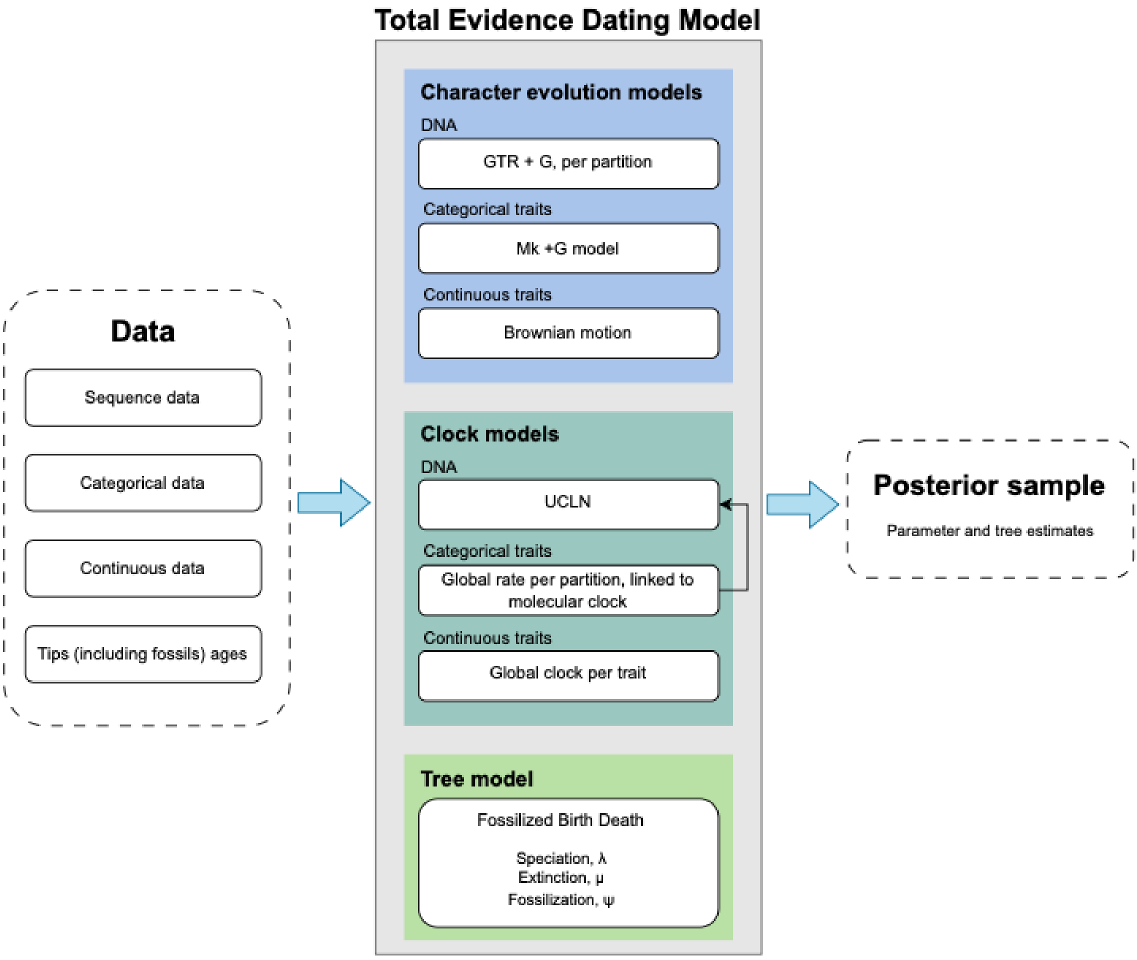

**Figure S2. 'No-time MCC tree'.** Preliminary total evidence tree including the three types of data (sequence data, categorical traits, and continuous traits) but using a uniform tree prior, effectively not modeling time. Fossil taxa are in red and numbers on each branch correspond to the posterior probability of each clade.

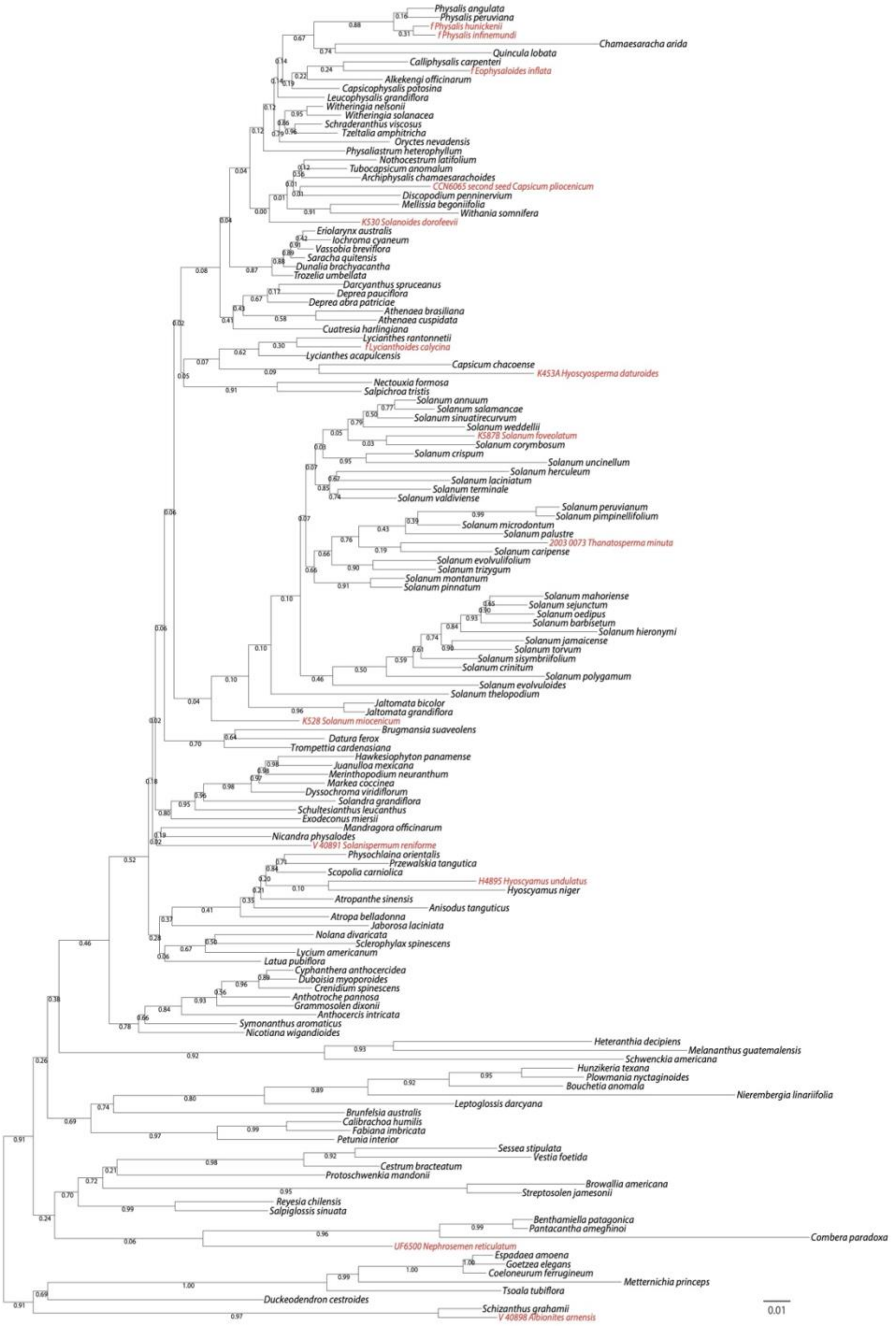

**Figure S3.** Topological convergence in MCMC chains for the time-homogeneous and time-heterogeneous models. For two independent chains of each model, we show the tree topology traces and the tree space heatmaps.

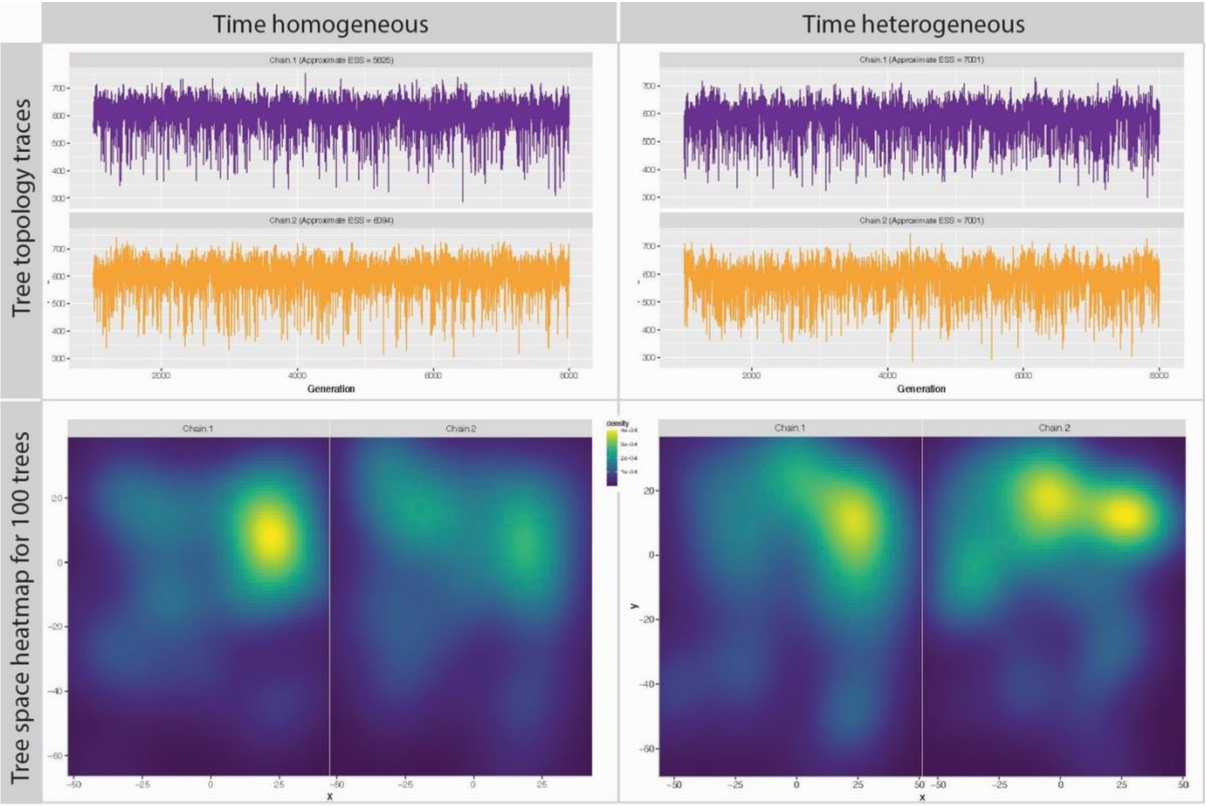

**Figure S4.** Maximum clade credibility trees from the time-homogeneous and time-heterogeneous models. Trees are pruned to show only extant taxa. The bar at each node denotes the 95% HPD for the date of the split, and the color of the bar indicates the posterior probability for the node.

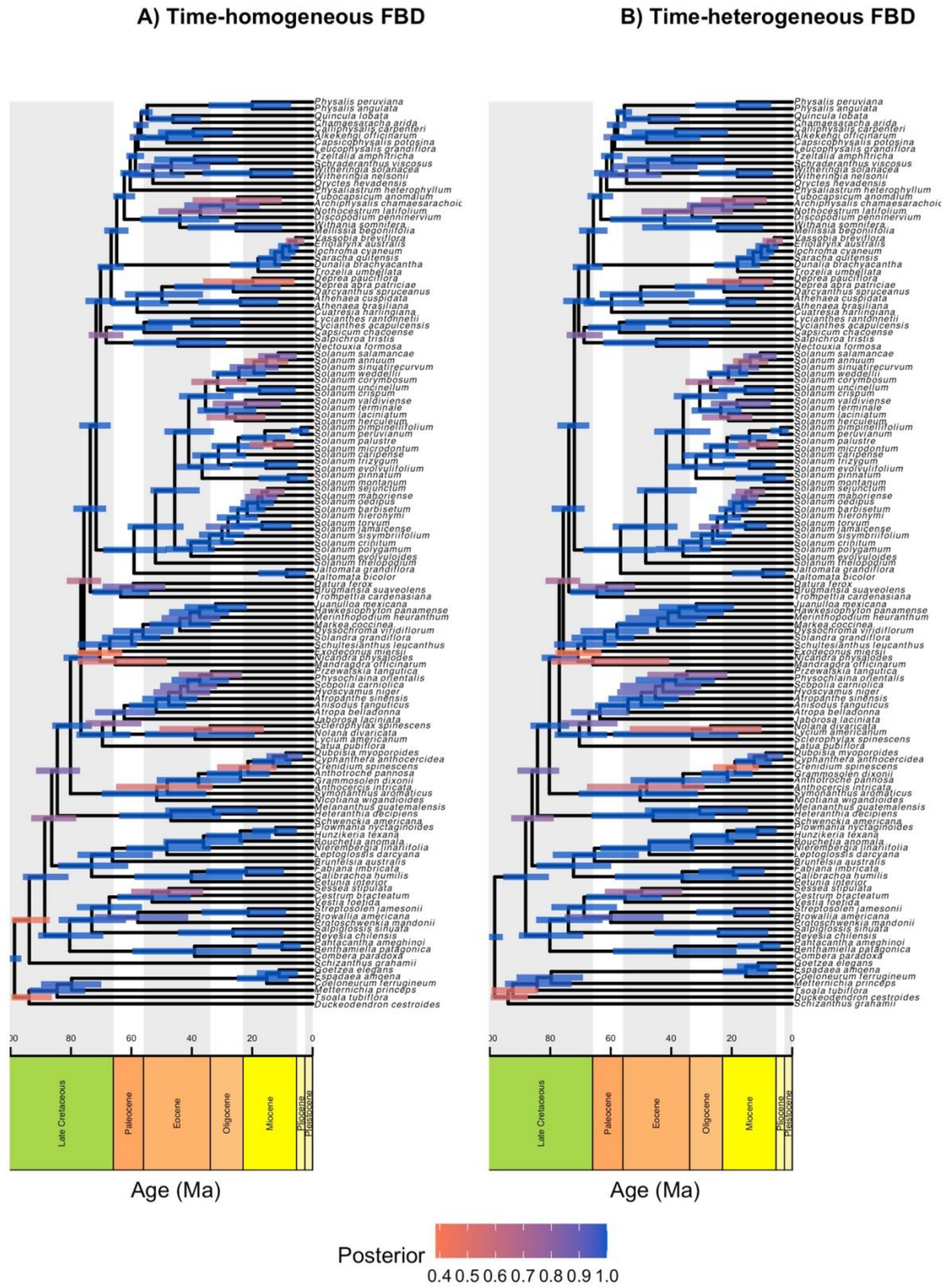

**Figure S5**. Maximum extant-only clade credibility trees from the time-homogeneous (left) and time-heterogeneous (right) FBD models. Blue lines connect the same taxa in the two different trees, thus highlighting topological differences between them.

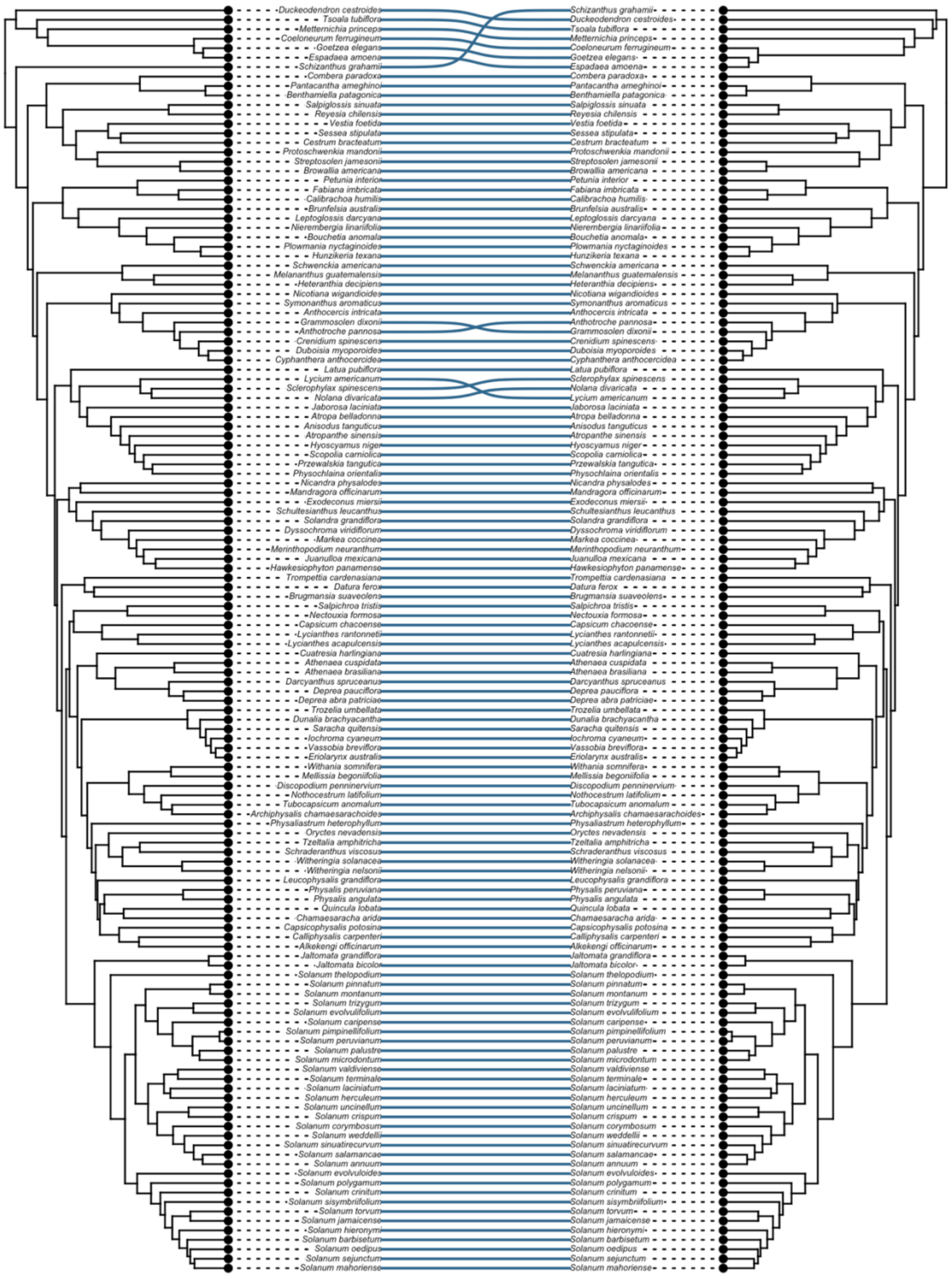

**Figure S6**. Differences in age estimates for clades of Solanaceae under the time homogeneous and the time heterogeneous FBD models. The dot inside each violin plot represents the median value, and the width of the curve indicates the density of points across the range of values. Clade names follow Figure 1.

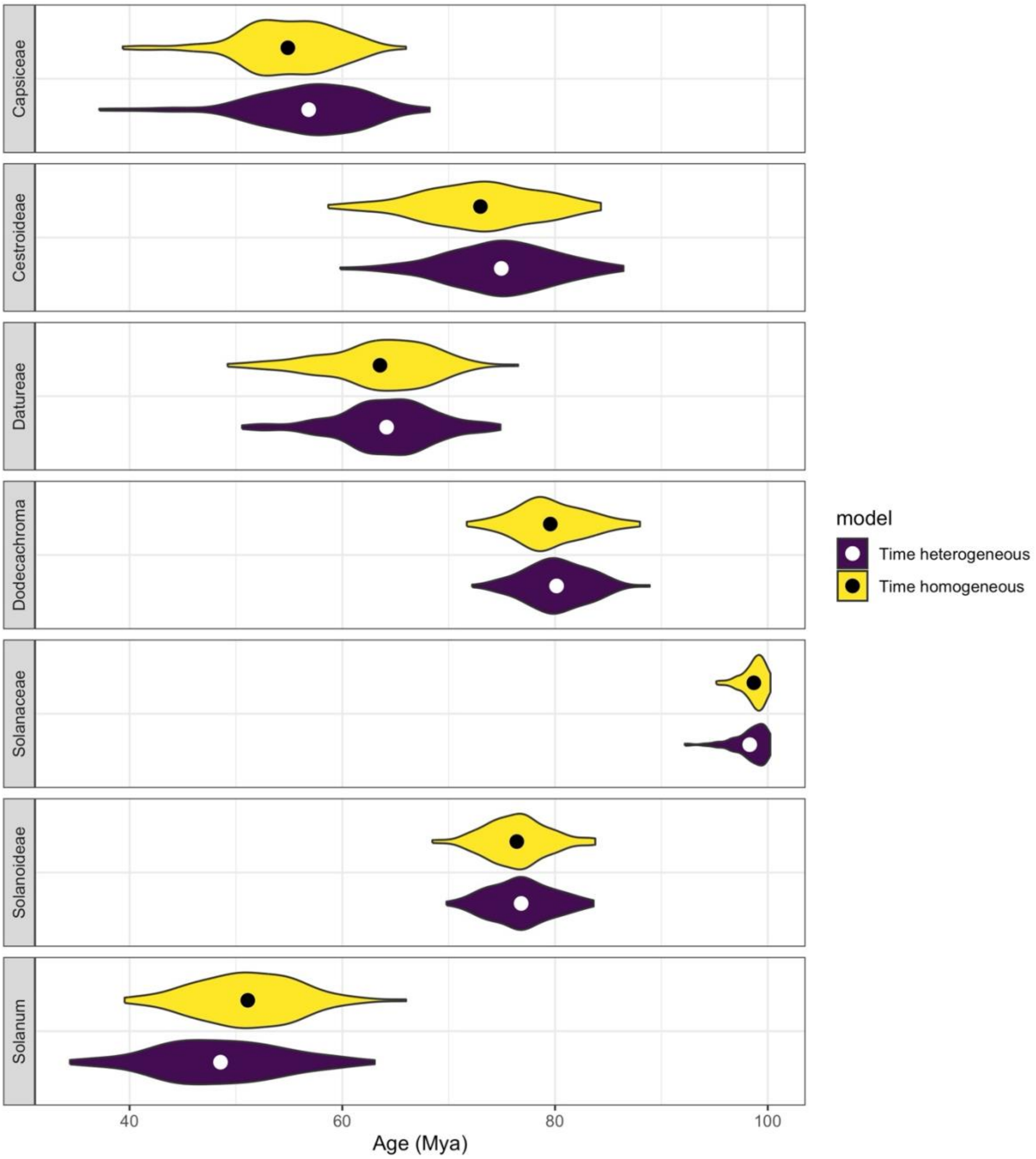

**Figure S7.** Maximum likelihood reconstruction of eight continuous morphological traits across the extant Solanaceae. Taxa without data were removed from the MCC tree.

**A.** Seed length

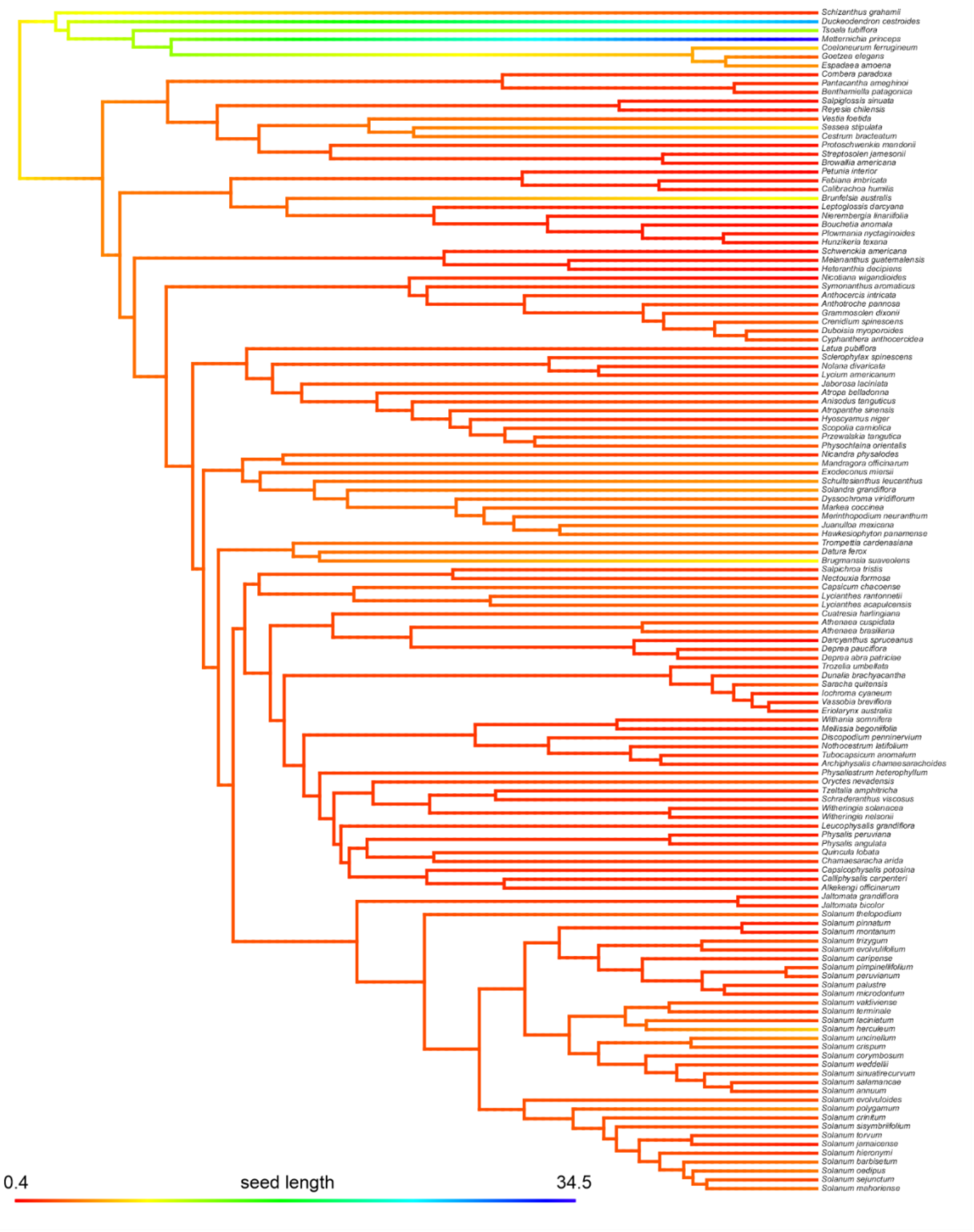

**B.** Seed length: width ratio

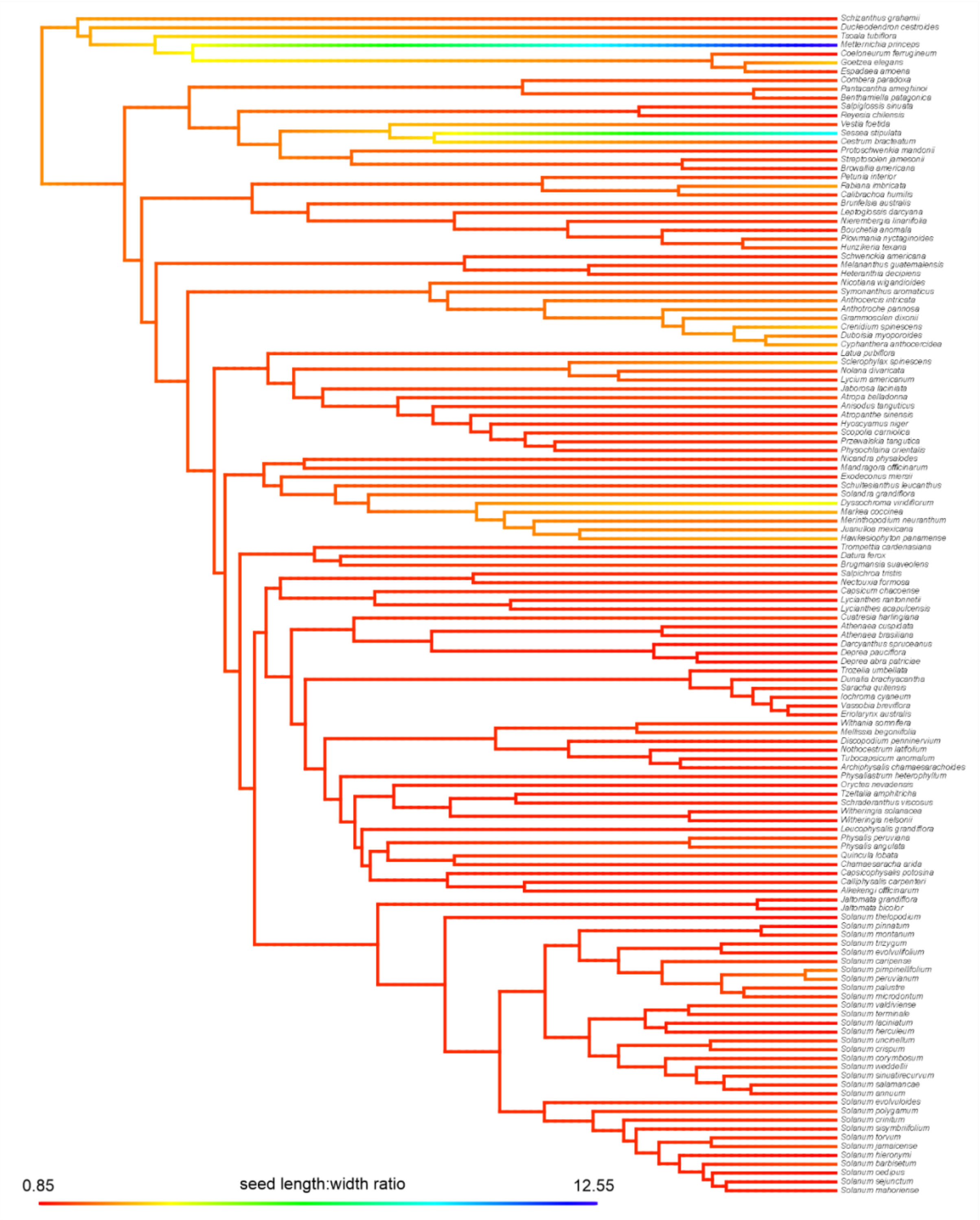

**C.** Fruiting calyx length

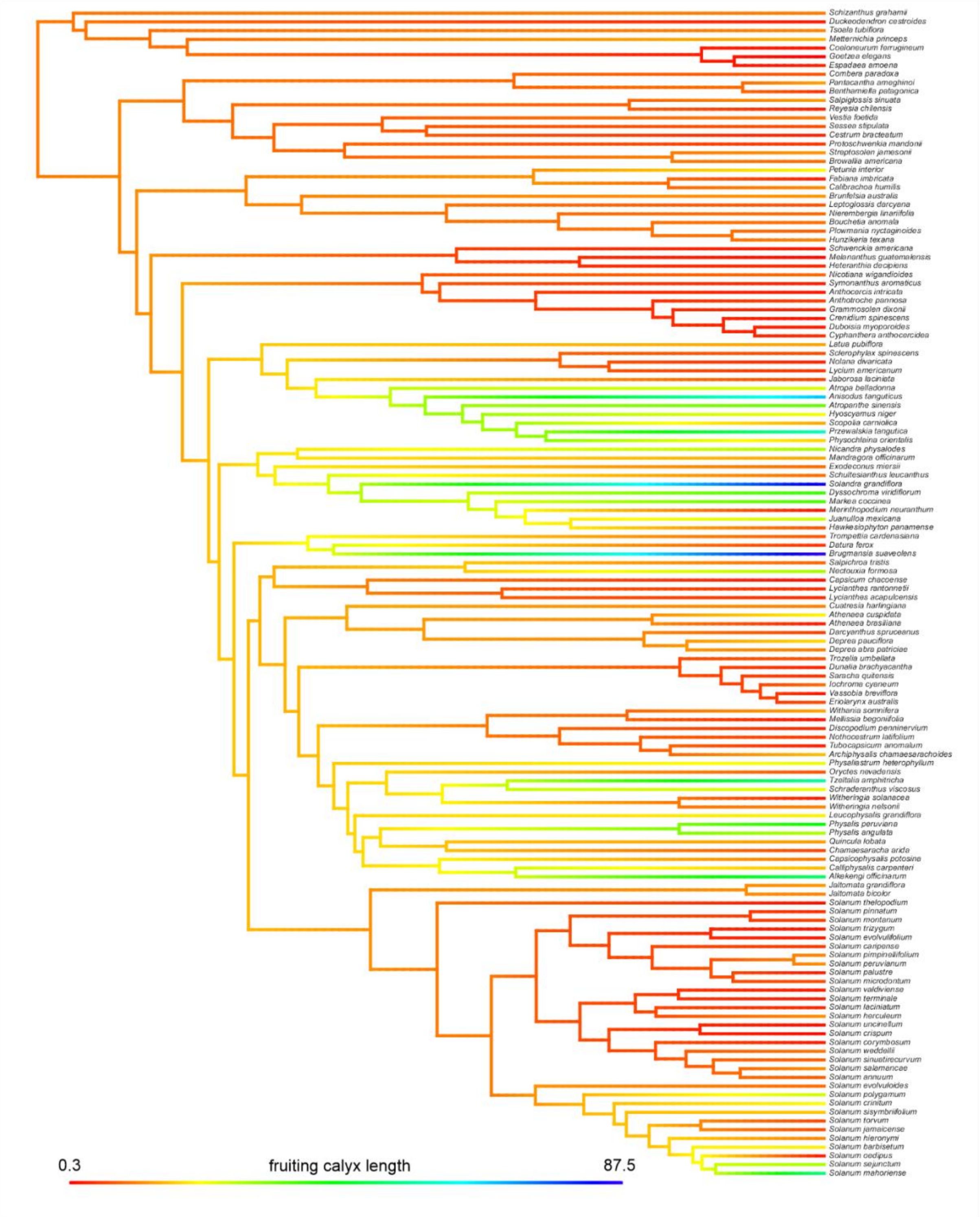

**D.** Fruiting calyx length: width ratio

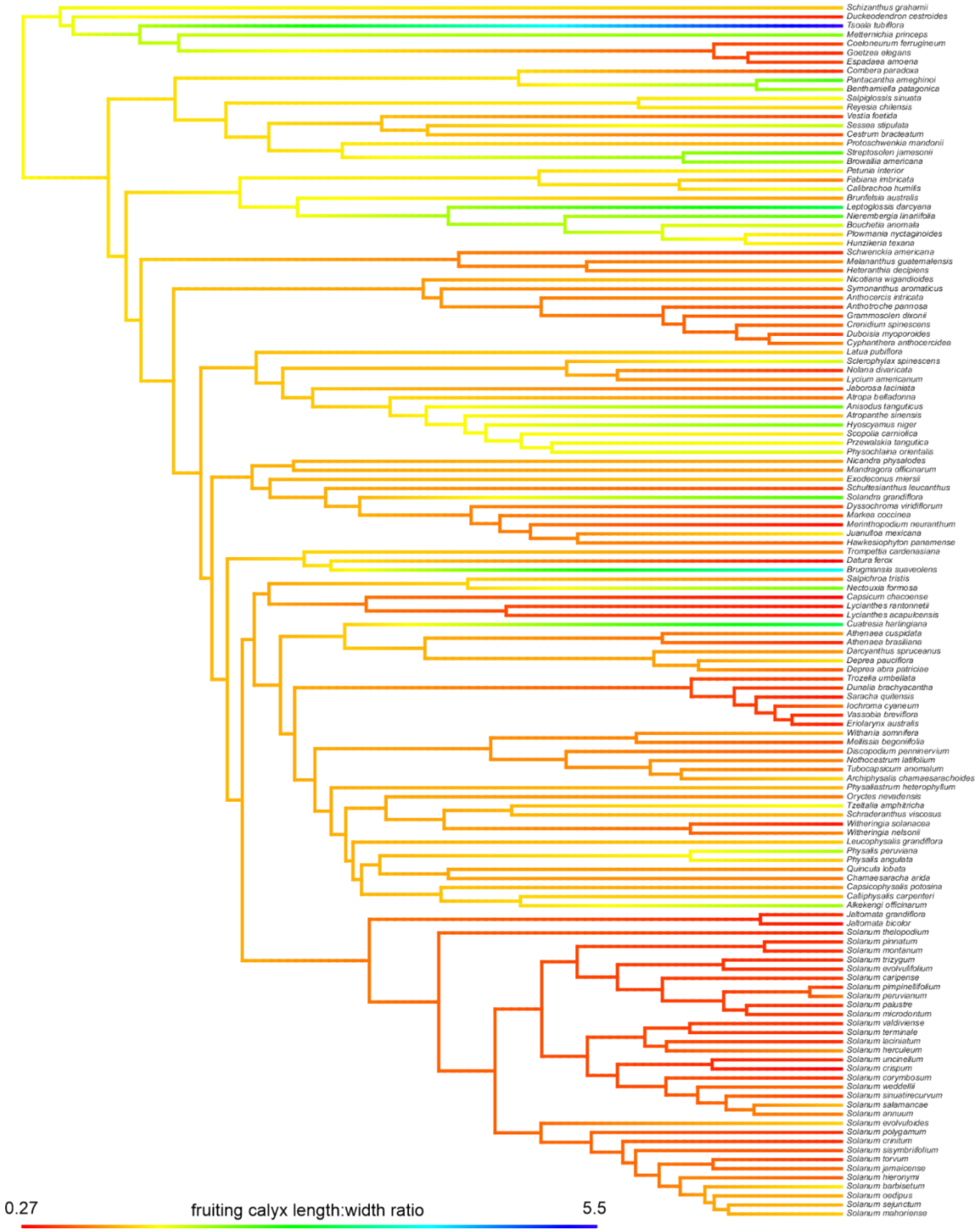

**E.** Length of fruiting calyx lobes

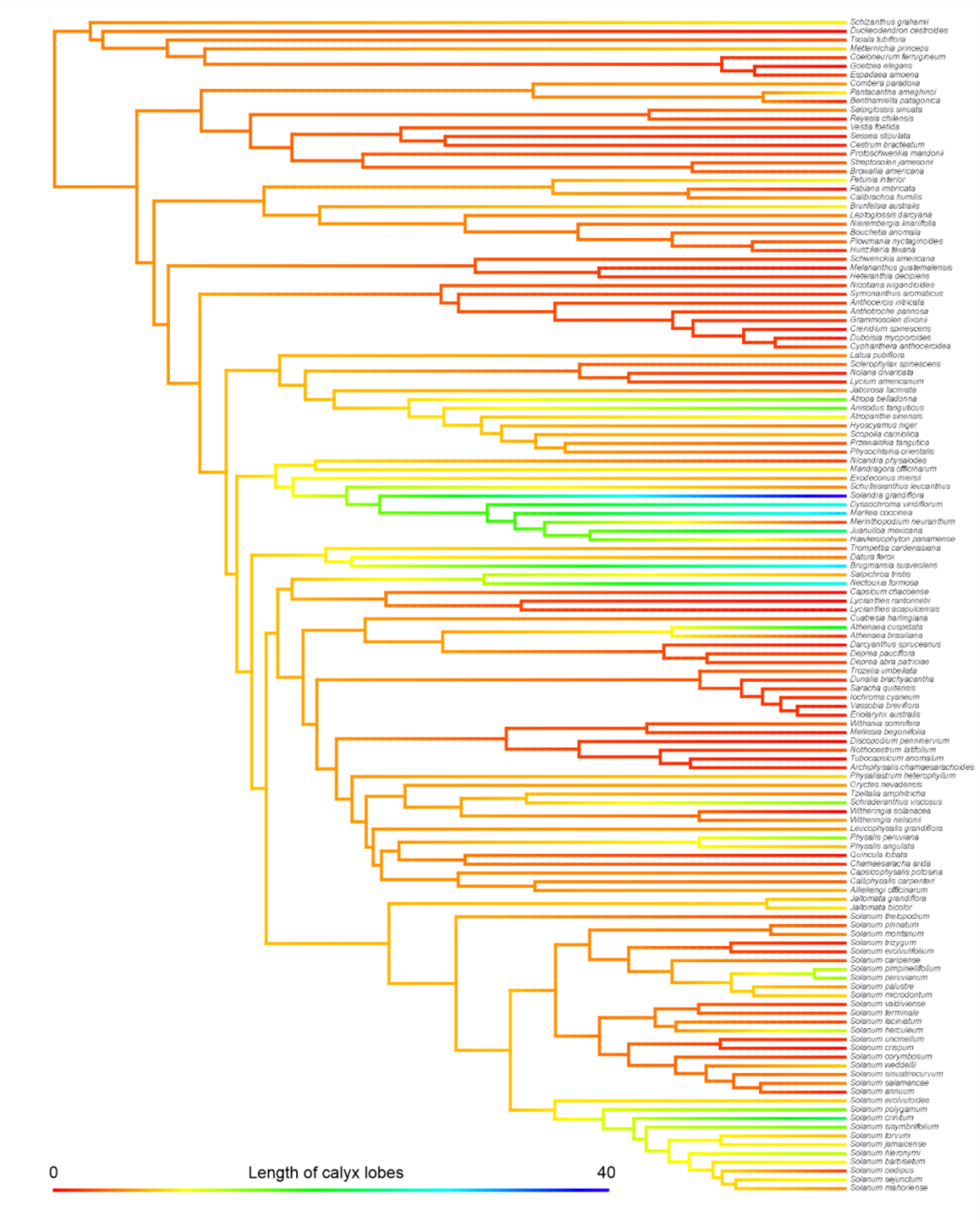

**F.** Fruit pedicel length

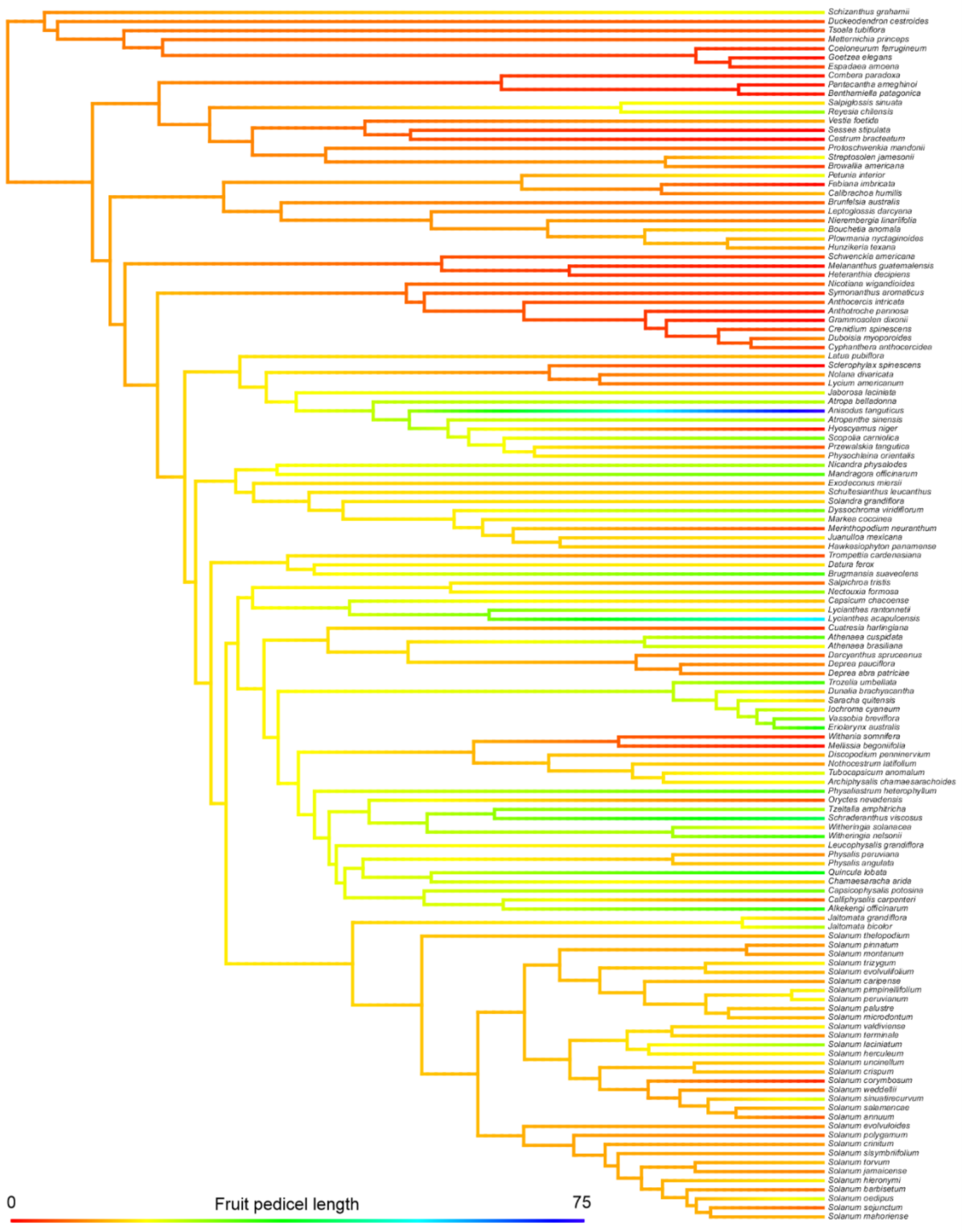

**G.** Calyx lobes length: total calyx length ratio

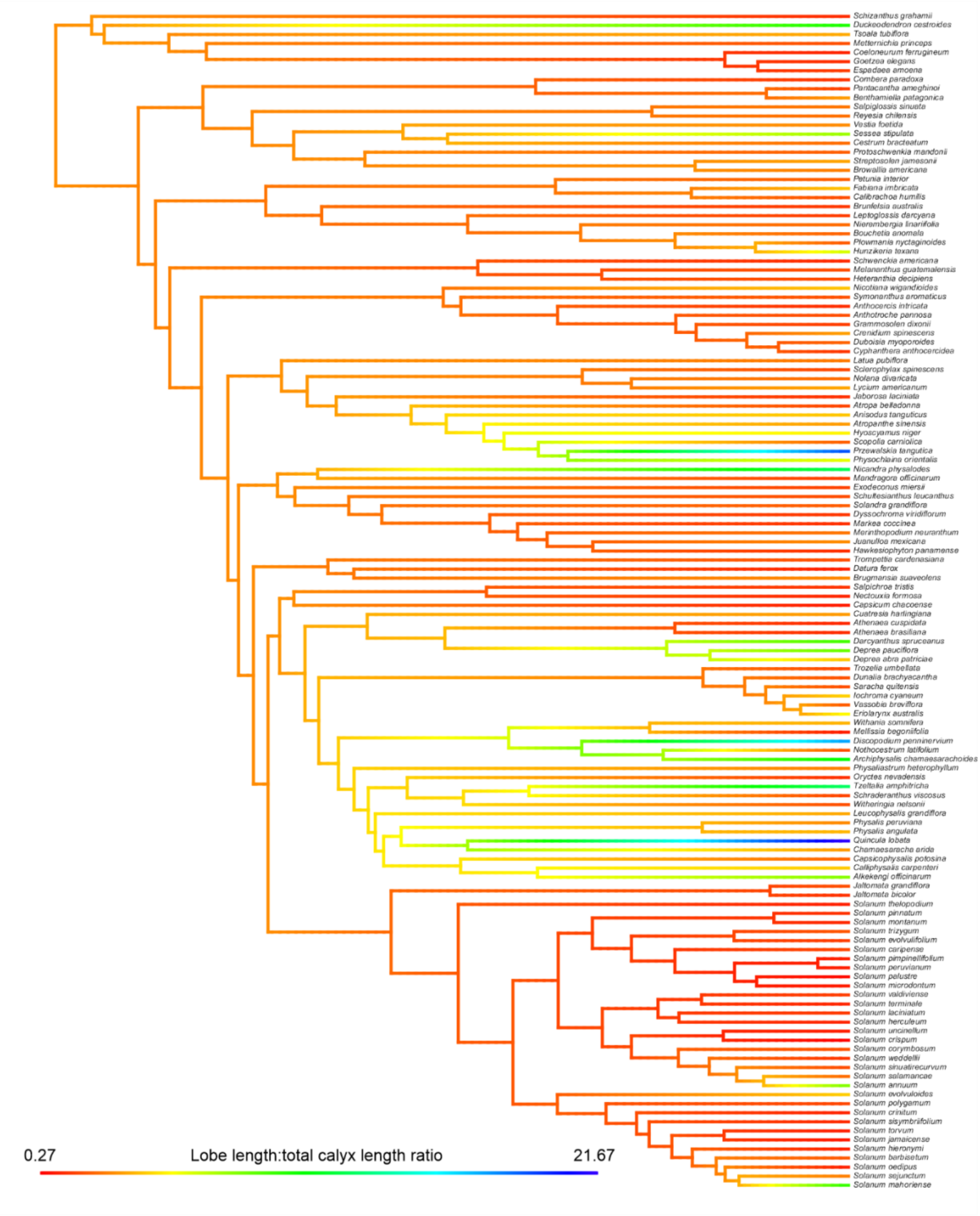

**H.** Fruit length: width ratio

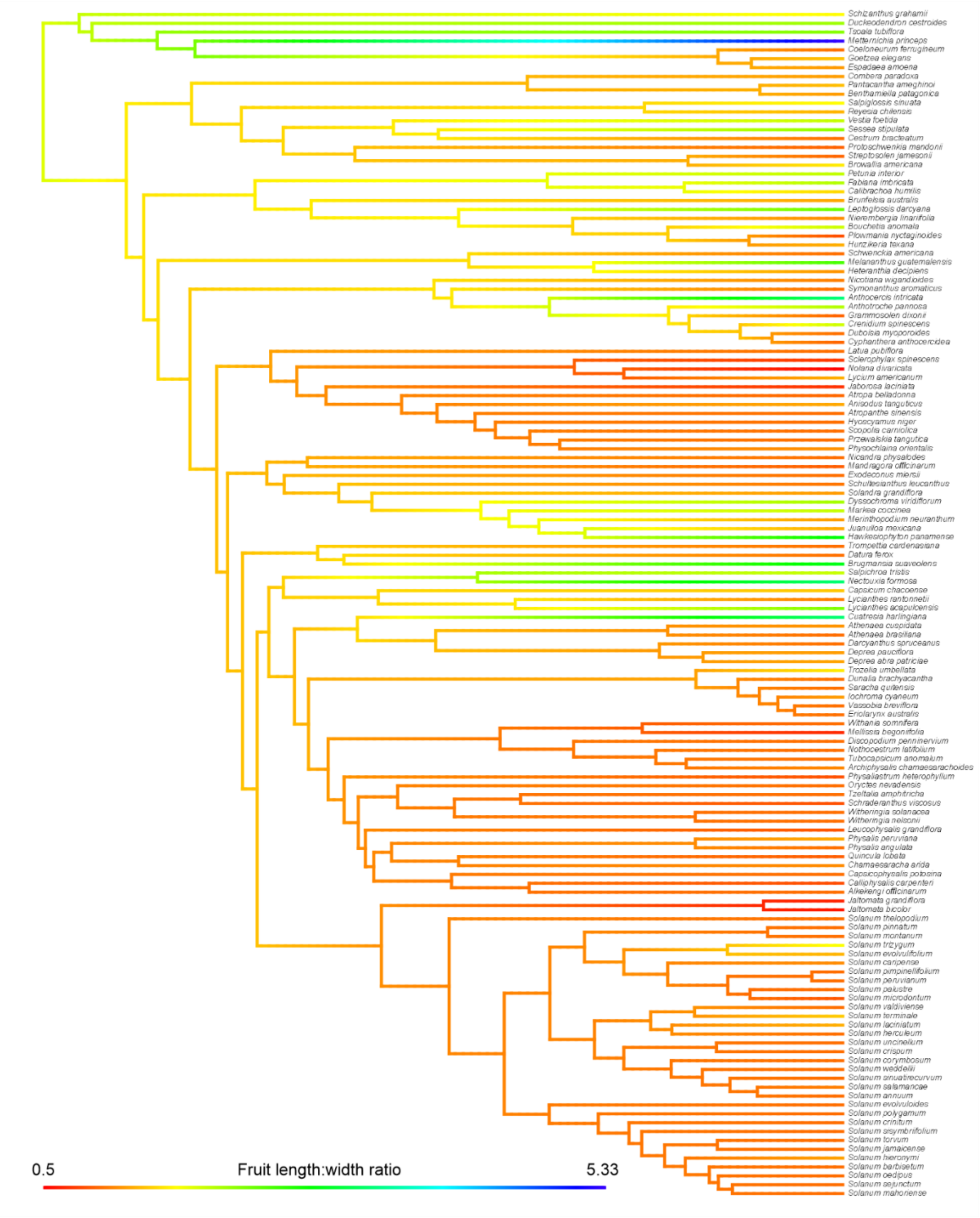

**Figure S8.** Ancestral state reconstruction of 17 categorical morphological traits across the extant Solanaceae using Bayesian stochastic mapping under the best-fitting model of trait evolution (ARD was the best model for embryo shape, fruiting calyx base invagination, fruiting calyx angled, calyx lobe sinus, fruiting calyx widest veins, fruit type, and fruiting calyx inflated; ER was the best model for seed compression, hilum position, seed hilar-chalazal cavity, exotestal cell walls, seed wings, seed elaiosomes, fruiting calyx secondary veins position and forking, calyx venation pattern, and fruiting calyx teeth). Pies at nodes indicate frequencies of node states across 1,000 simulations of character evolution, and the colors of the tip labels represent tip states.

**A.** Seed compression

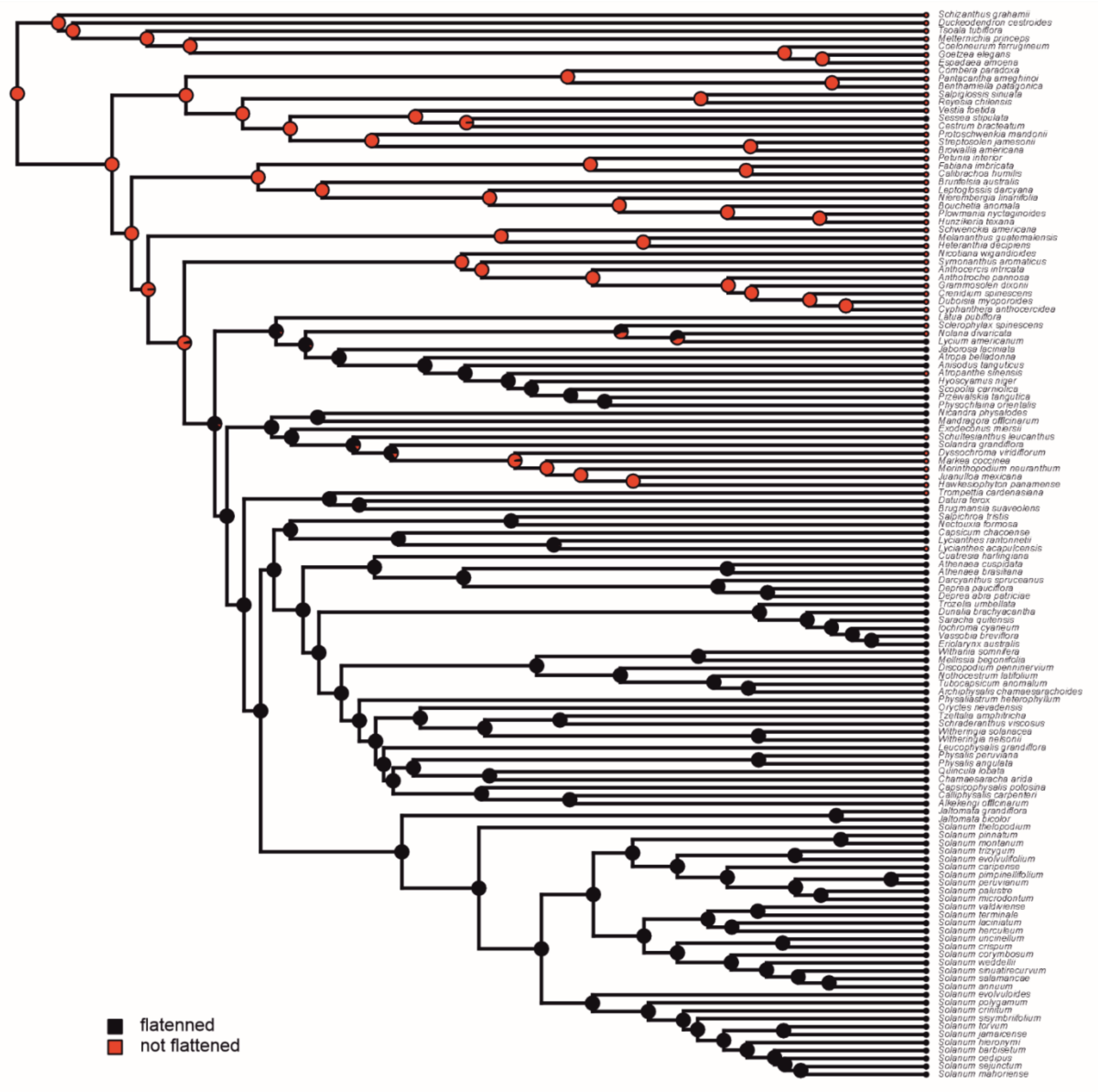

**B.** Hilum position

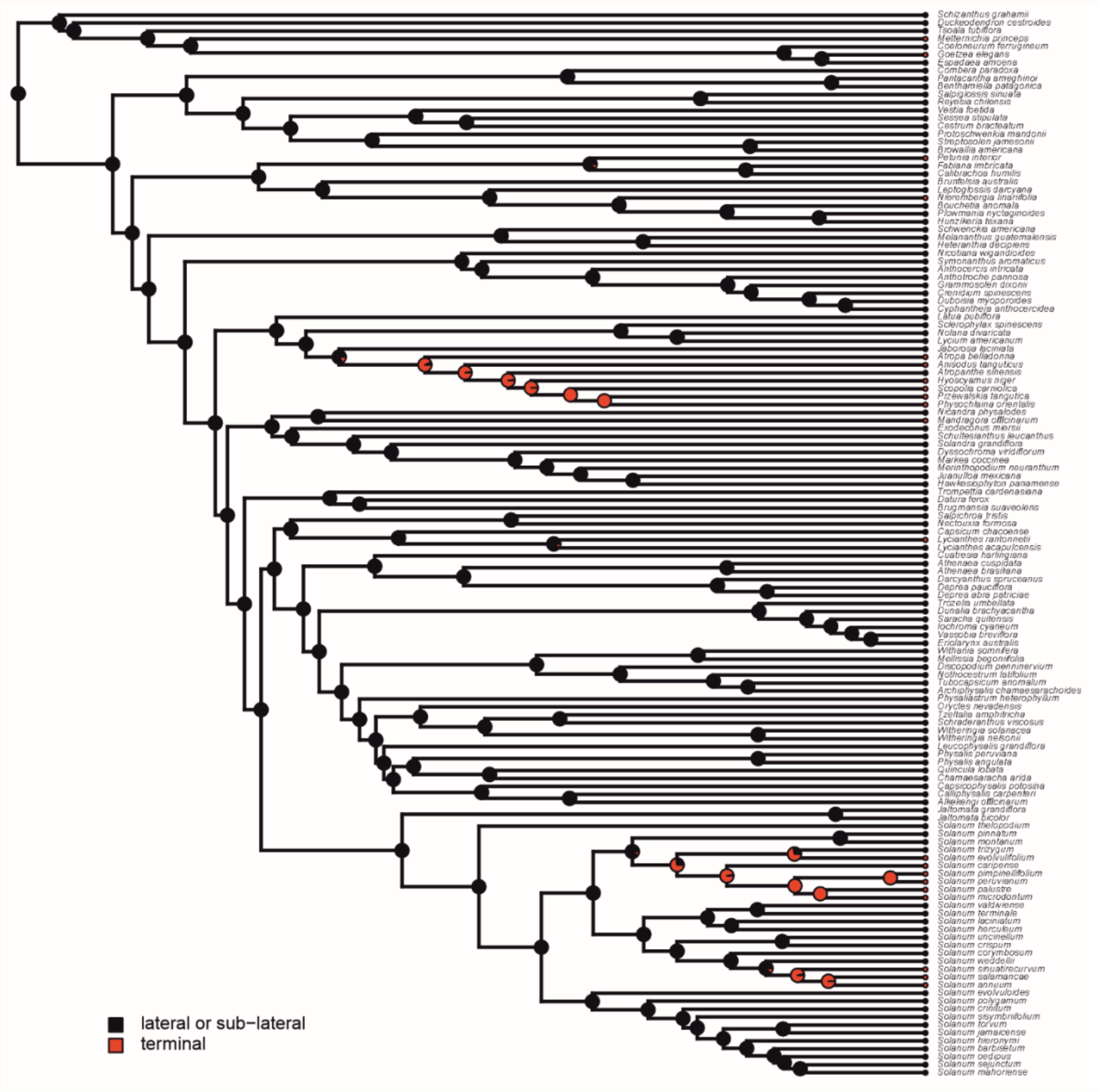

**C.** Embryo shape

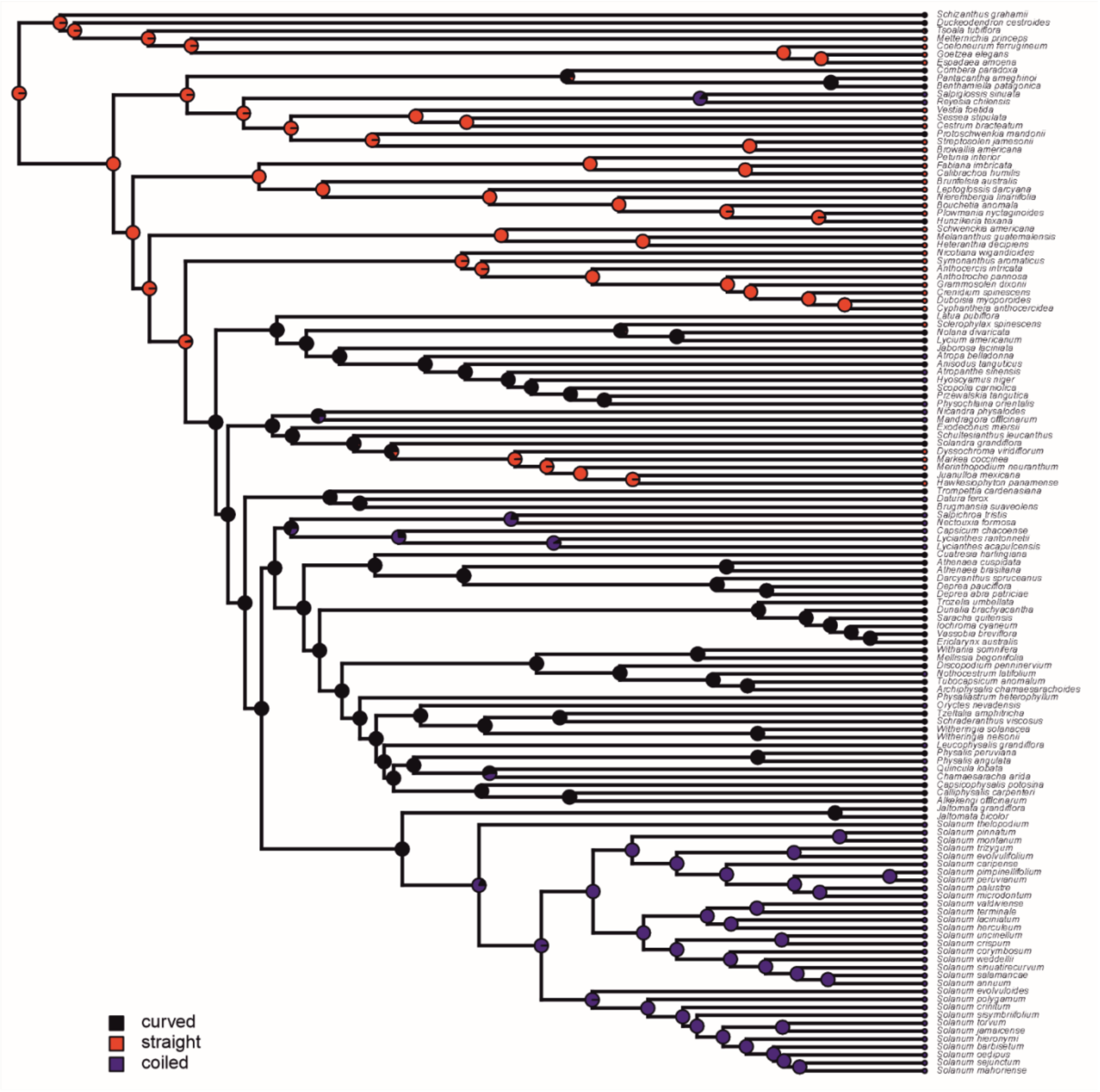

**D.** Seed hilar-chalazal cavity

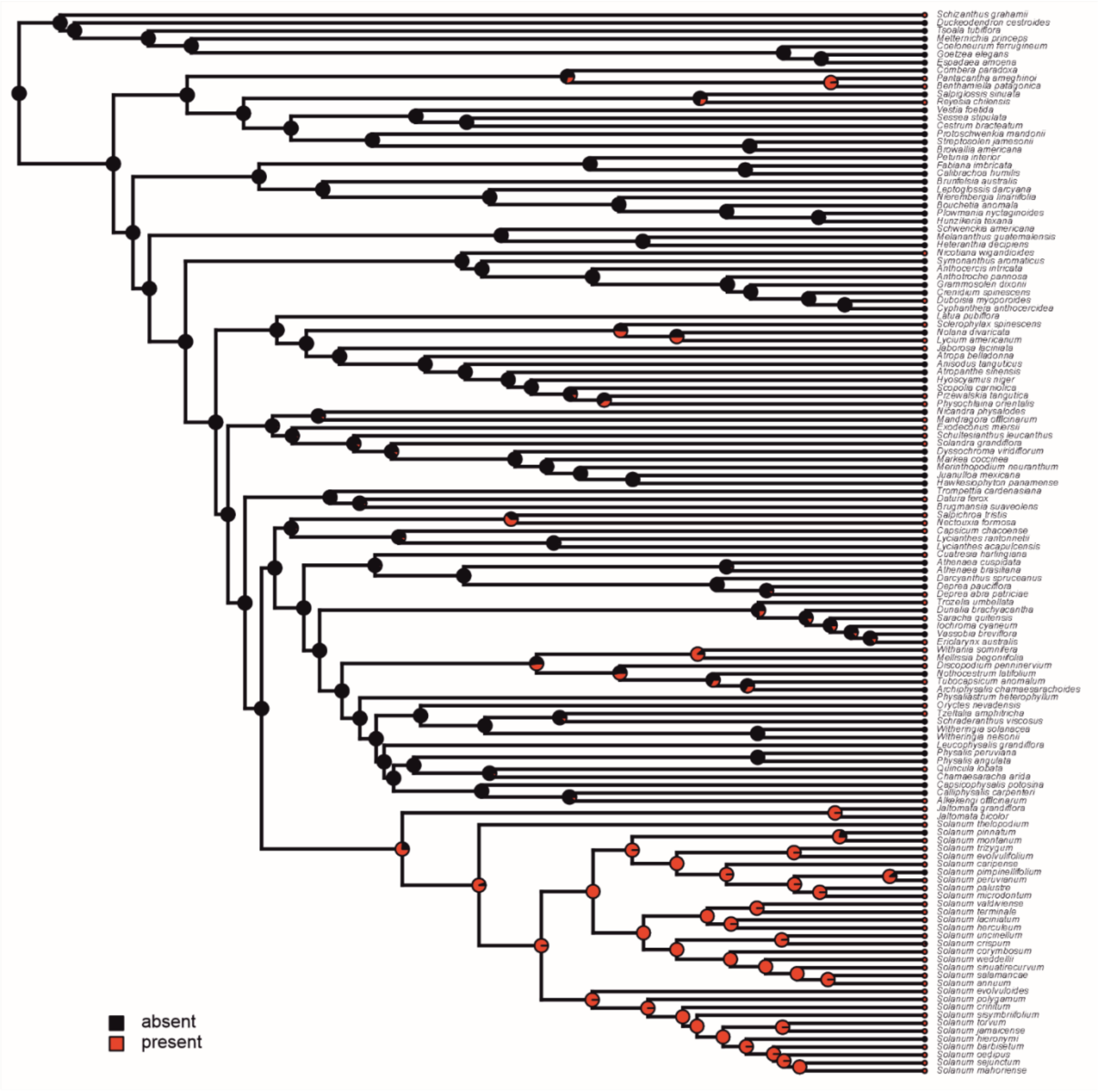

**E.** Seed exotestal cell walls

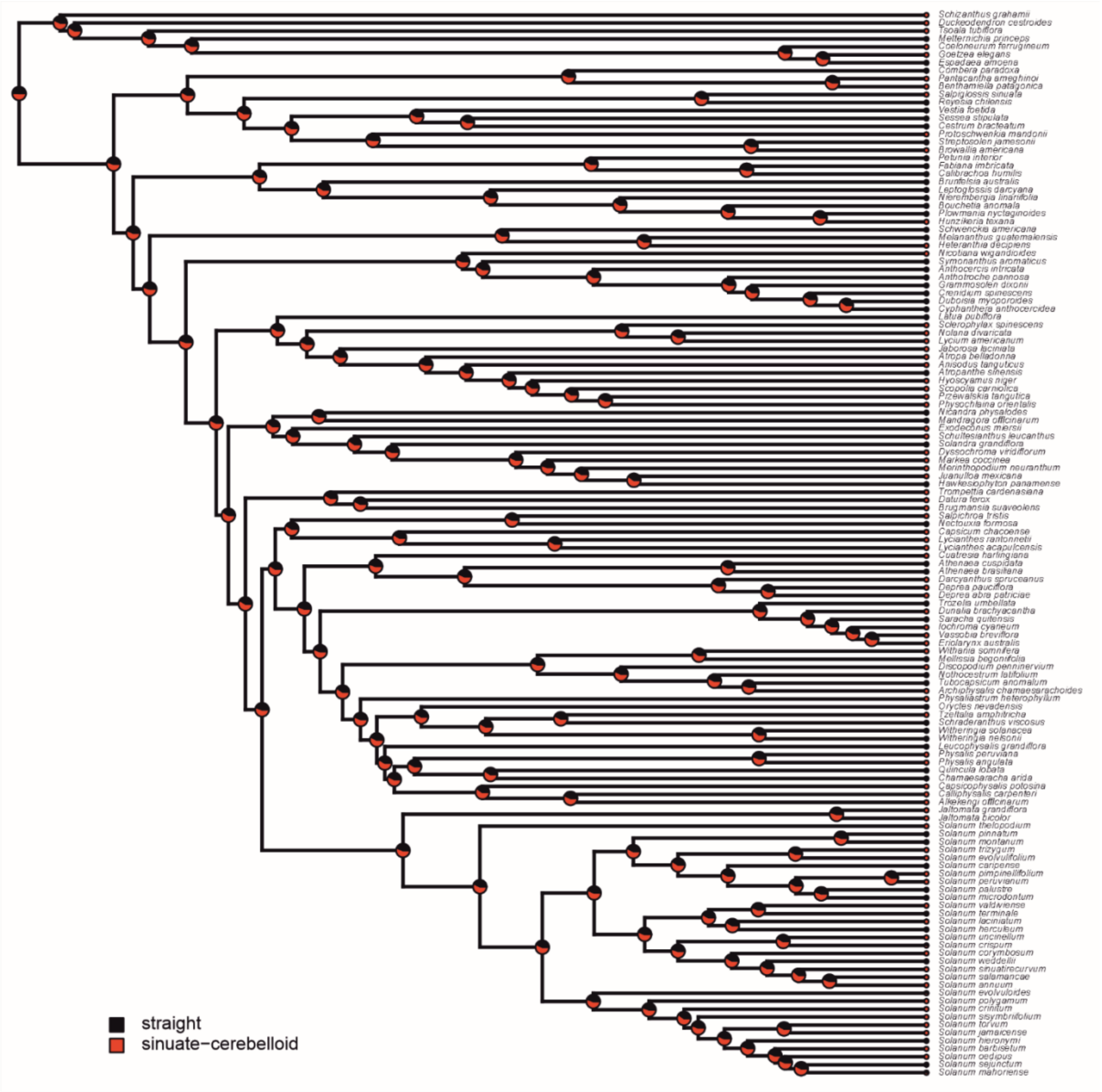

**F.** Seed wings

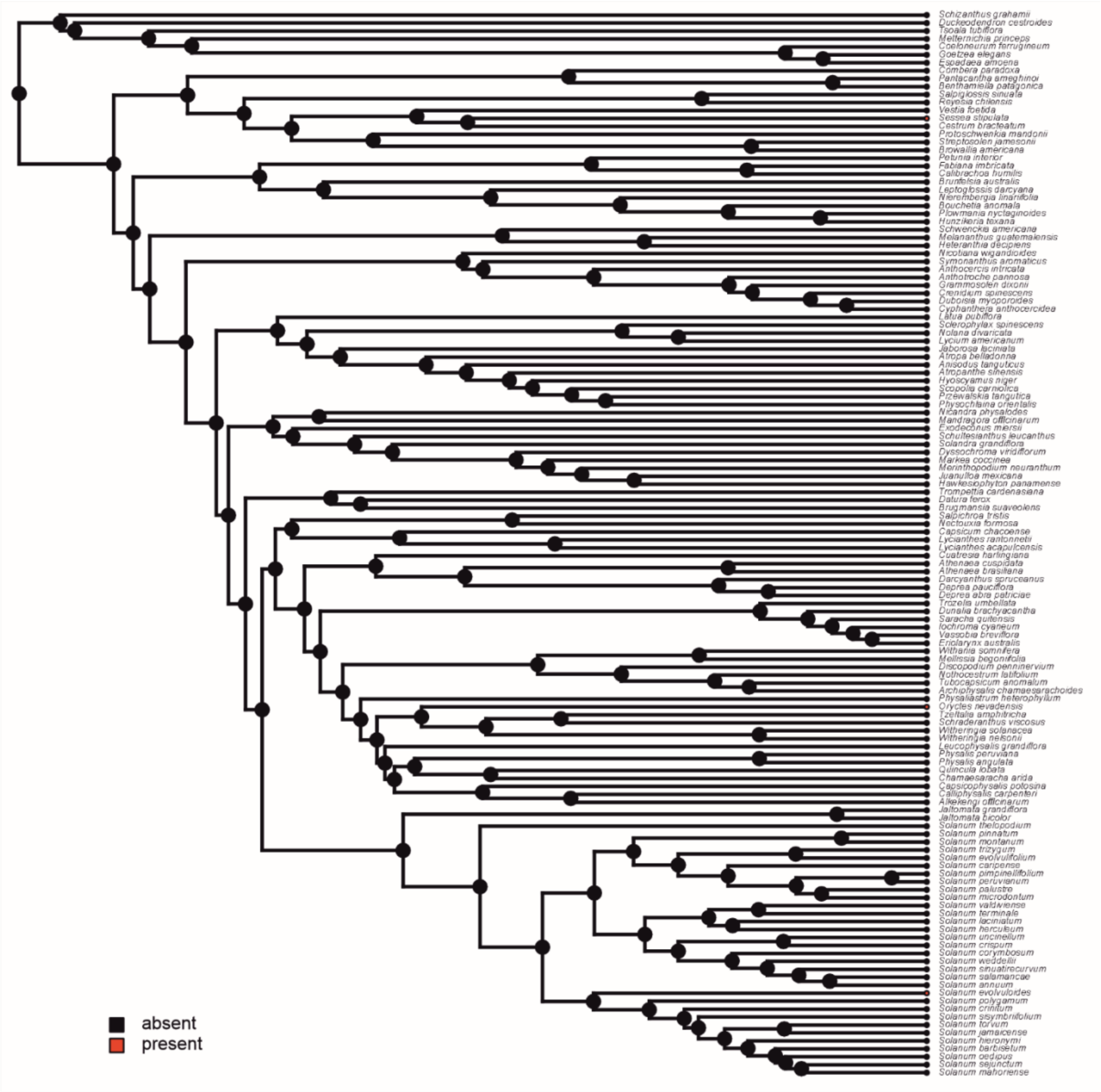
Schizanthus grahamii
Duckeodendron cestroides
Tsoala tubiflora
Metternichia princeps
Coeloneurum ferrugineum
Goetzea elegans
Espadaea amoena
Combera paradoxa
Pantacantha ameghinoi
Benthamiella patagonica
Salpiglossis sinuata
Reyesia chilensis
Vestia foetida
Sessea stipulata
Cestrum bracteatum
Protoschwenkia mandonii
Streptosolen jamesonii
Browallia americana
Petunia interior
Fabiana imbricata
Calibrachoa humilis
Brunfelsia australis
Leptoglossis darcyana
Nierembergia linariifolia
Bouchetia anomala
Plowmania nyctaginoides
Hunzikeria texana
Schwenckia americana
Melananthus guatemalensis
Heteranthia decipiens
Nicotiana wigandioides
Symonanthus aromaticus
Anthocercis intricata
Anthotroche pannosa
Grammosolen dixonii
Crenidium spinescens
Duboisia myoporoides
Cyphanthera anthocercidea
Latua pubiflora
Sclerophylax spinescens
Nolana divaricata
Lycium americanum
Jaborosa laciniata
Atropa belladonna
Anisodus tanguticus
Atropanthe sinensis
Hyoscyamus niger
Scopolia carniolica
Przewalskia tangutica
Physochlaina orientalis
Nicandra physalodes
Mandragora officinarum
Exodeconus miersii
Schultesianthus leucanthus
Solandra grandiflora
Dyssochroma viridiflorum
Markea coccinea
Merinthopodium neuranthum
Juanulloa mexicana
Hawkesiophyton panamense
Trompettia cardenasiana
Datura ferox
Brugmansia suaveolens
Salpichroa tristis
Nectouxia formosa
Capsicum chacoense
Lycianthes rantonnetii
Lycianthes acapulcensis
Cuatresia harlingiana
Athenaea cuspidata
Athenaea brasiliana
Darcyanthus spruceanus
Deprea pauciflora
Deprea abra patriciae
Trozelia umbellata
Dunalia brachyacantha
Saracha quitensis
Iochroma cyaneum
Vassobia breviflora
Eriolarynx australis
Withania somnifera
Mellissia begoniifolia
Discopodium penninervium
Nothocestrum latifolium
Tubocapsicum anomalum
Archiphysalis chamaesarachoides
Physaliastrum heterophyllum
Oryctes nevadensis
Tzeltalia amphitricha
Schraderanthus viscosus
Witheringia solanacea
Witheringia nelsonii
Leucophysalis grandiflora
Physalis peruviana
Physalis angulata
Quincula lobata
Chamaesaracha arida
Capsicophysalis potosina
Calliphysalis carpenteri
Alkekengi officinarum
Jaltomata grandiflora
Jaltomata bicolor
Solanum thelopodium
Solanum pinnatum
Solanum montanum
Solanum trizygum
Solanum evolvulifolium
Solanum caripense
Solanum pimpinellifolium
Solanum peruvianum
Solanum palustre
Solanum microdontum
Solanum valdiviense
Solanum terminale
Solanum laciniatum
Solanum herculeum
Solanum uncinellum
Solanum crispum
Solanum corymbosum
Solanum weddellii
Solanum sinuatirecurvum
Solanum salamancae
Solanum annuum
Solanum evolvuloides
Solanum polygamum
Solanum crinitum
Solanum sisymbriifolium
Solanum torvum
Solanum jamaicense
Solanum hieronymi
Solanum barbisetum
Solanum oedipus
Solanum sejunctum
Solanum mahoriense
absent
present

**G.** Seed elaiosomes

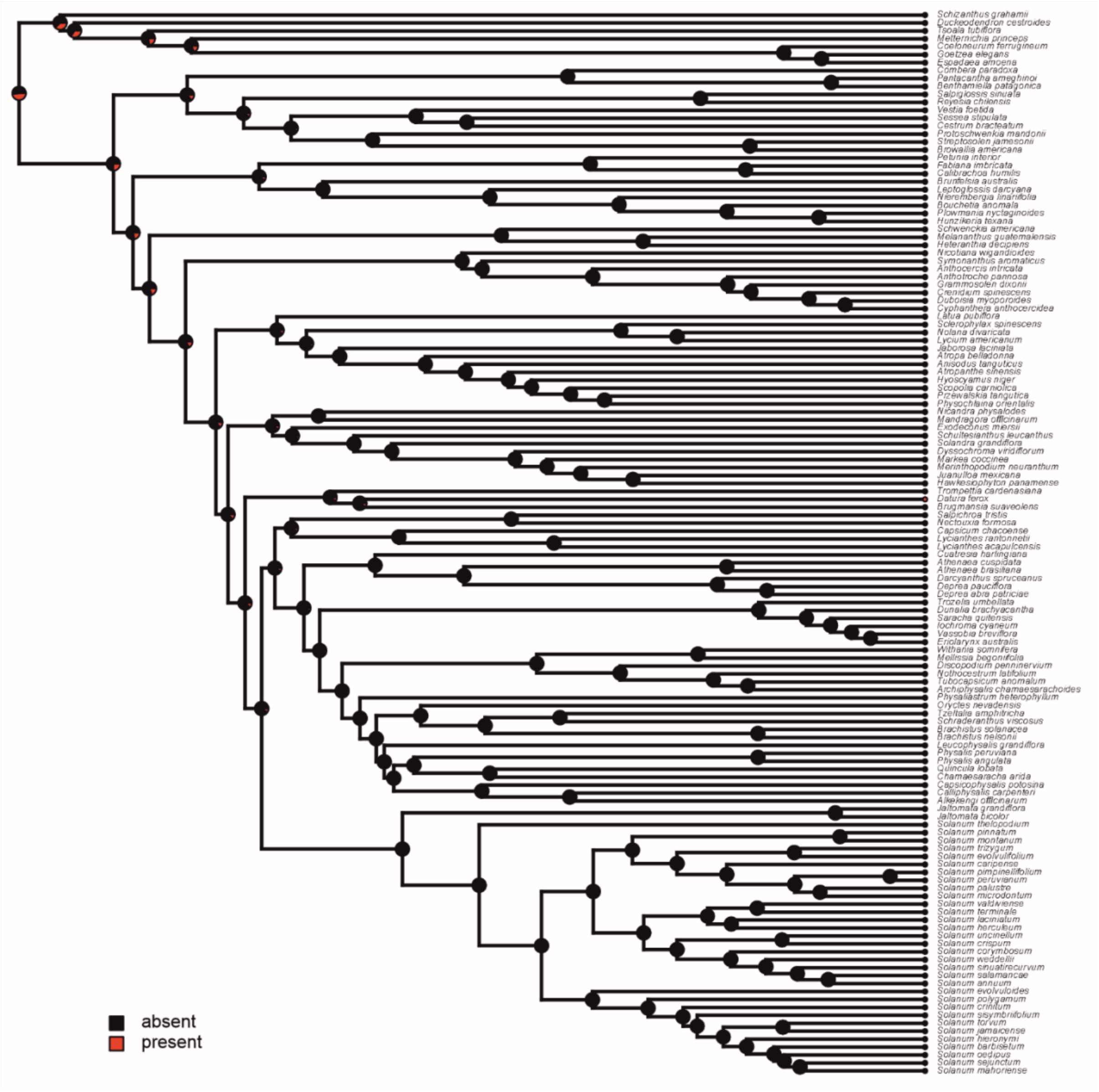

**H.** Fruiting calyx base invagination

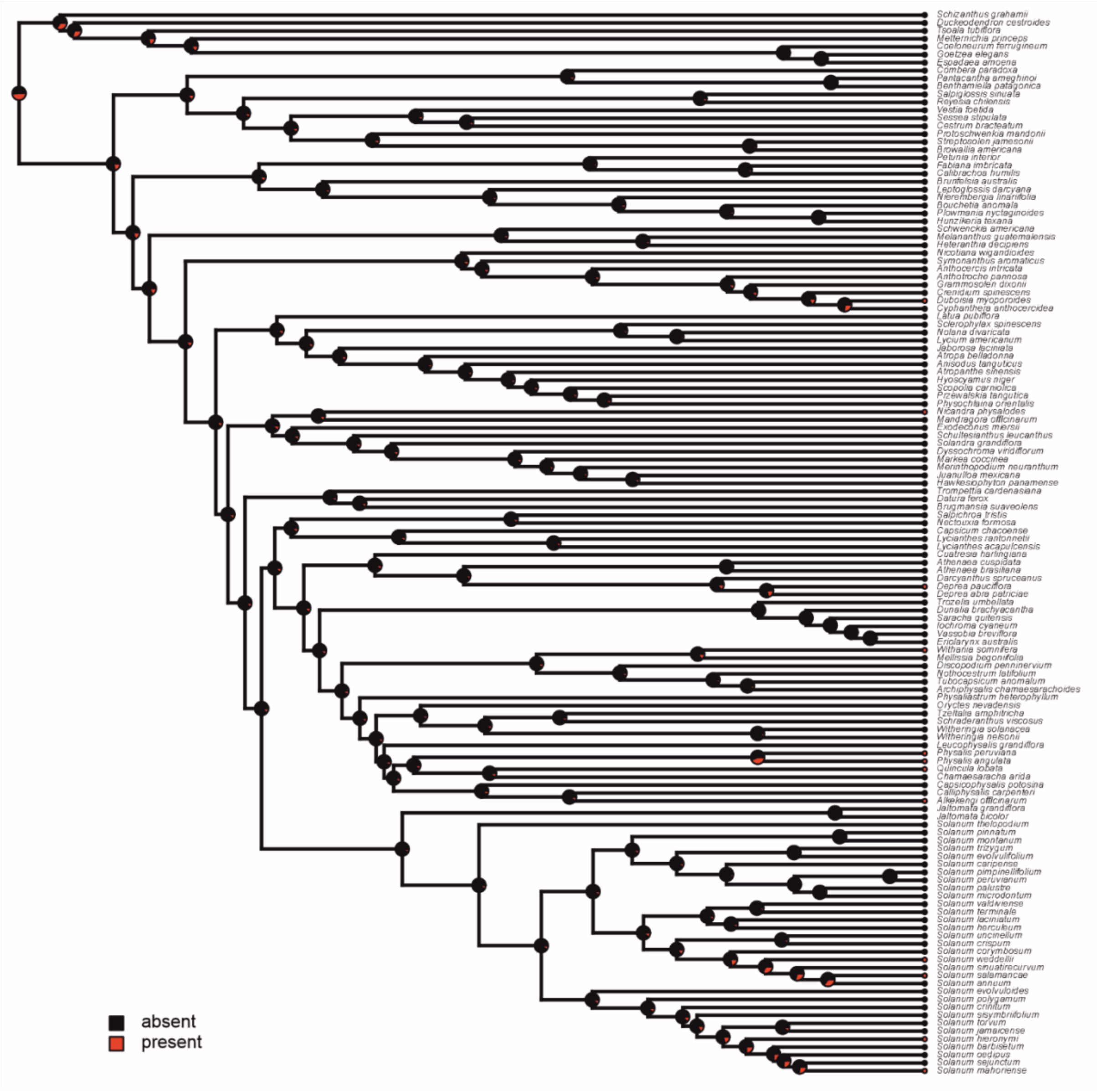

**I.** Fruiting calyx angled

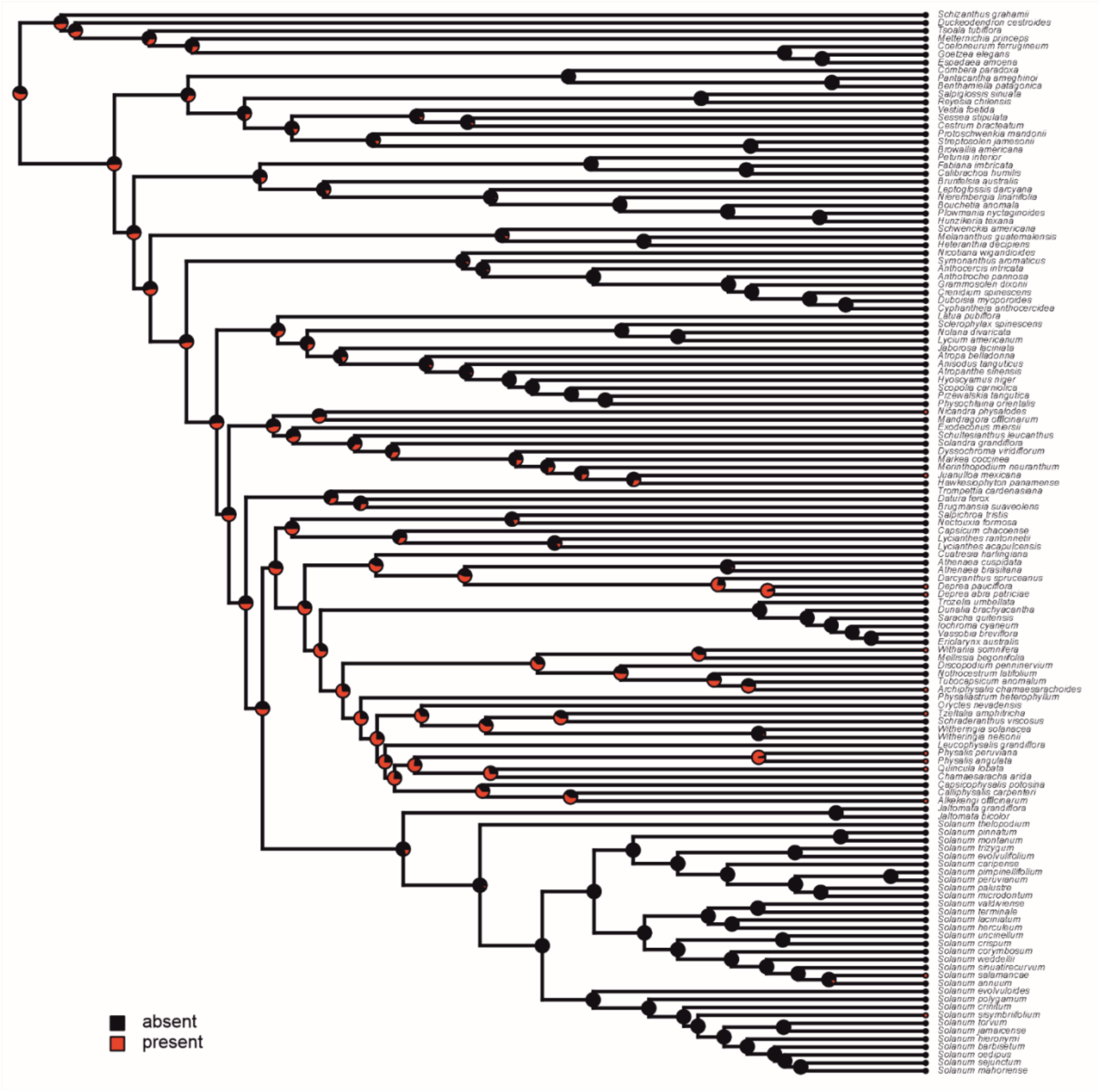

**I.** Fruiting calyx lobe sinus

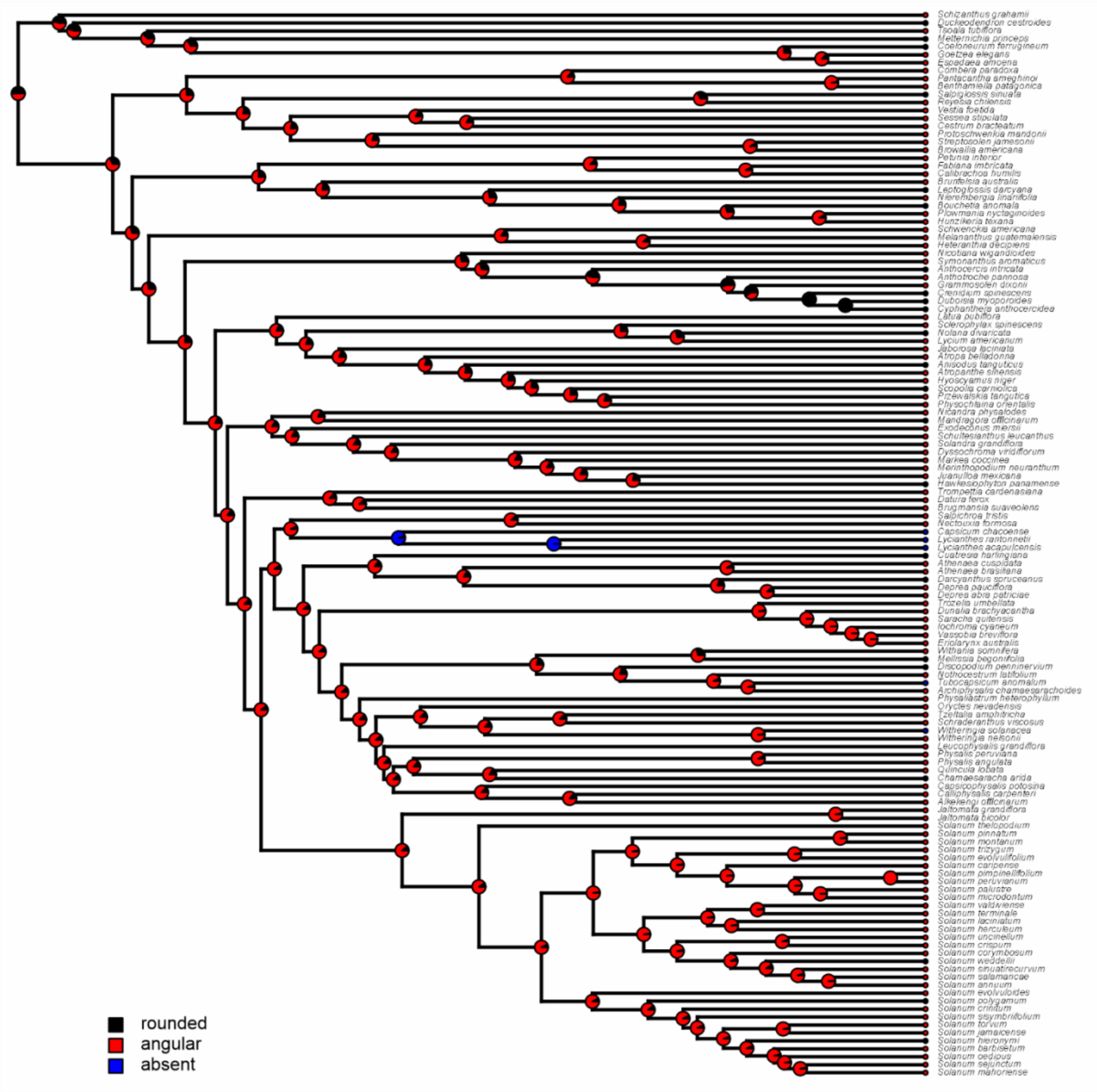

**J.** Fruiting calyx widest veins distinct and terminating in lobe tips

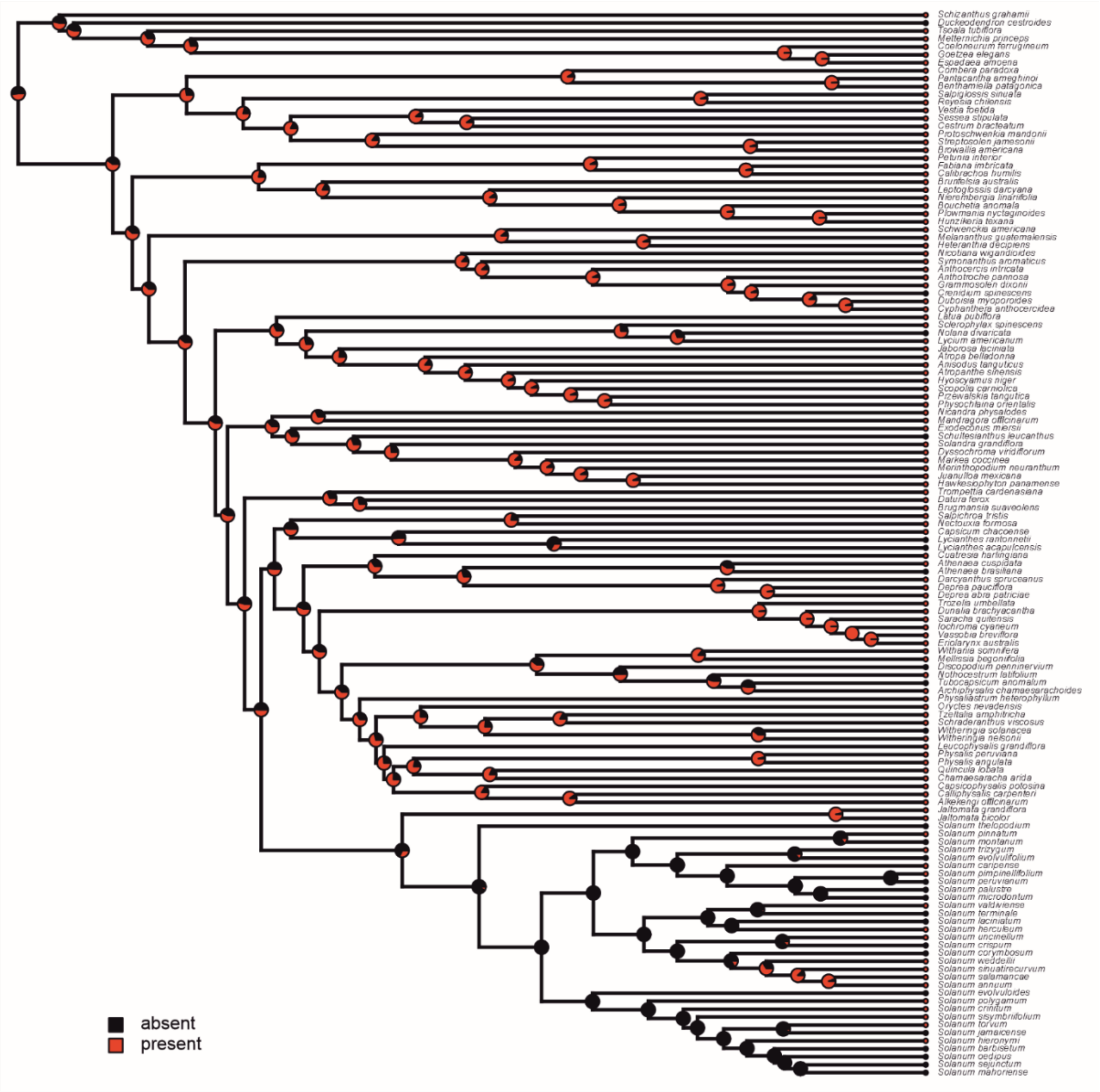

**K.** Fruiting calyx secondary veins distinct from other vein orders and emerge from the base

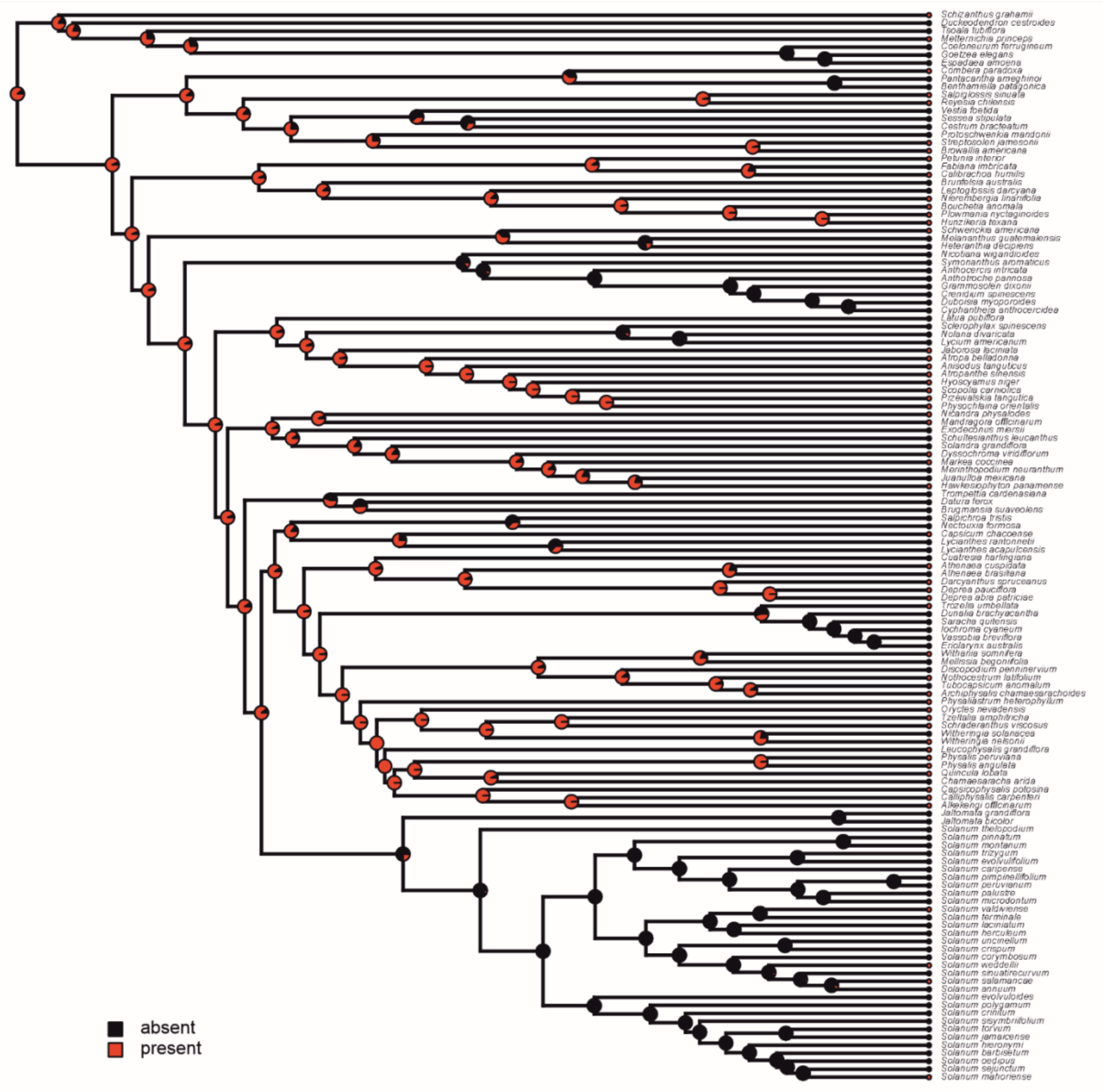

**L.** Fruiting calyx secondary veins fork before the lobe sinus

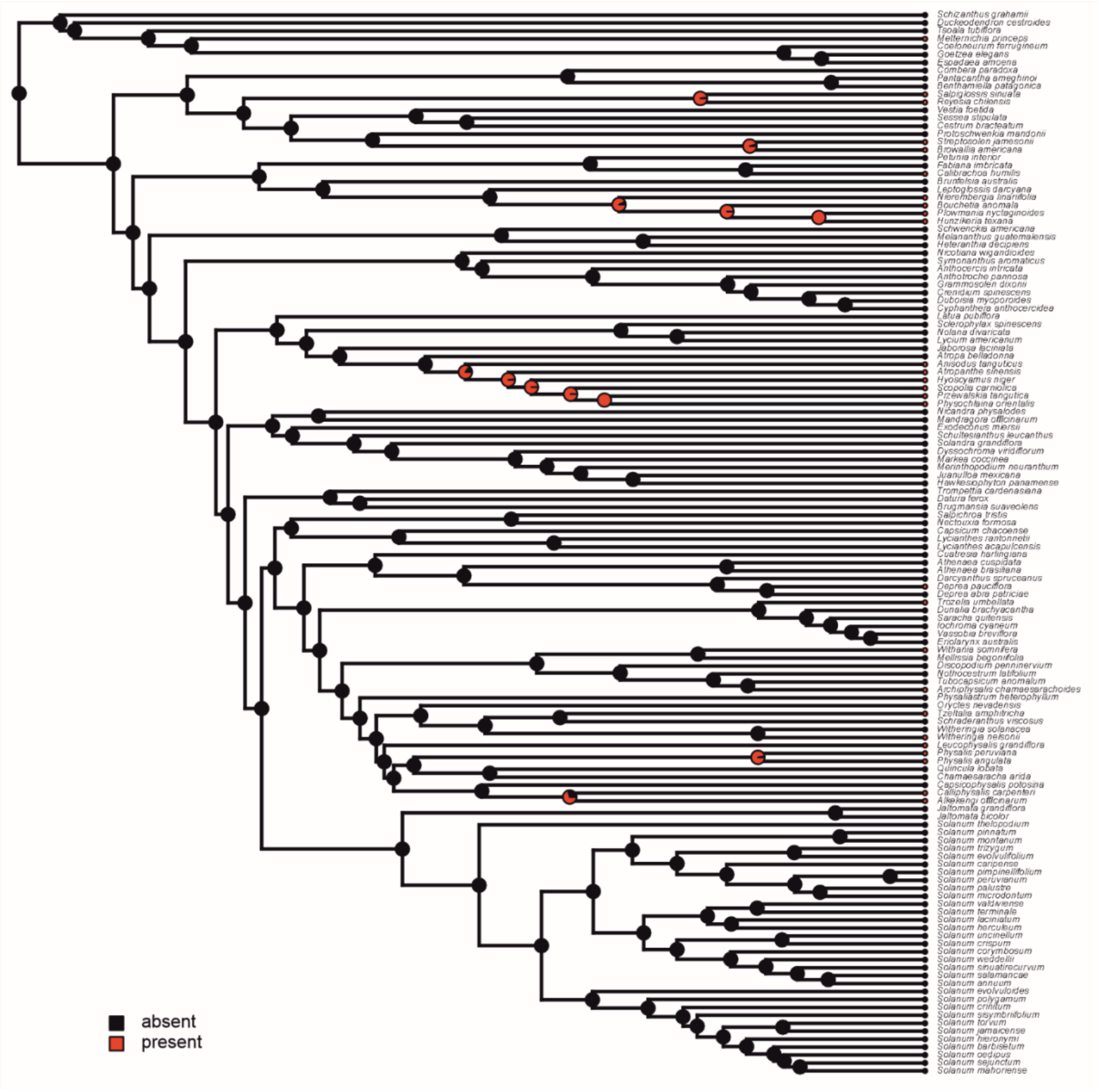

**M.** Fruit type

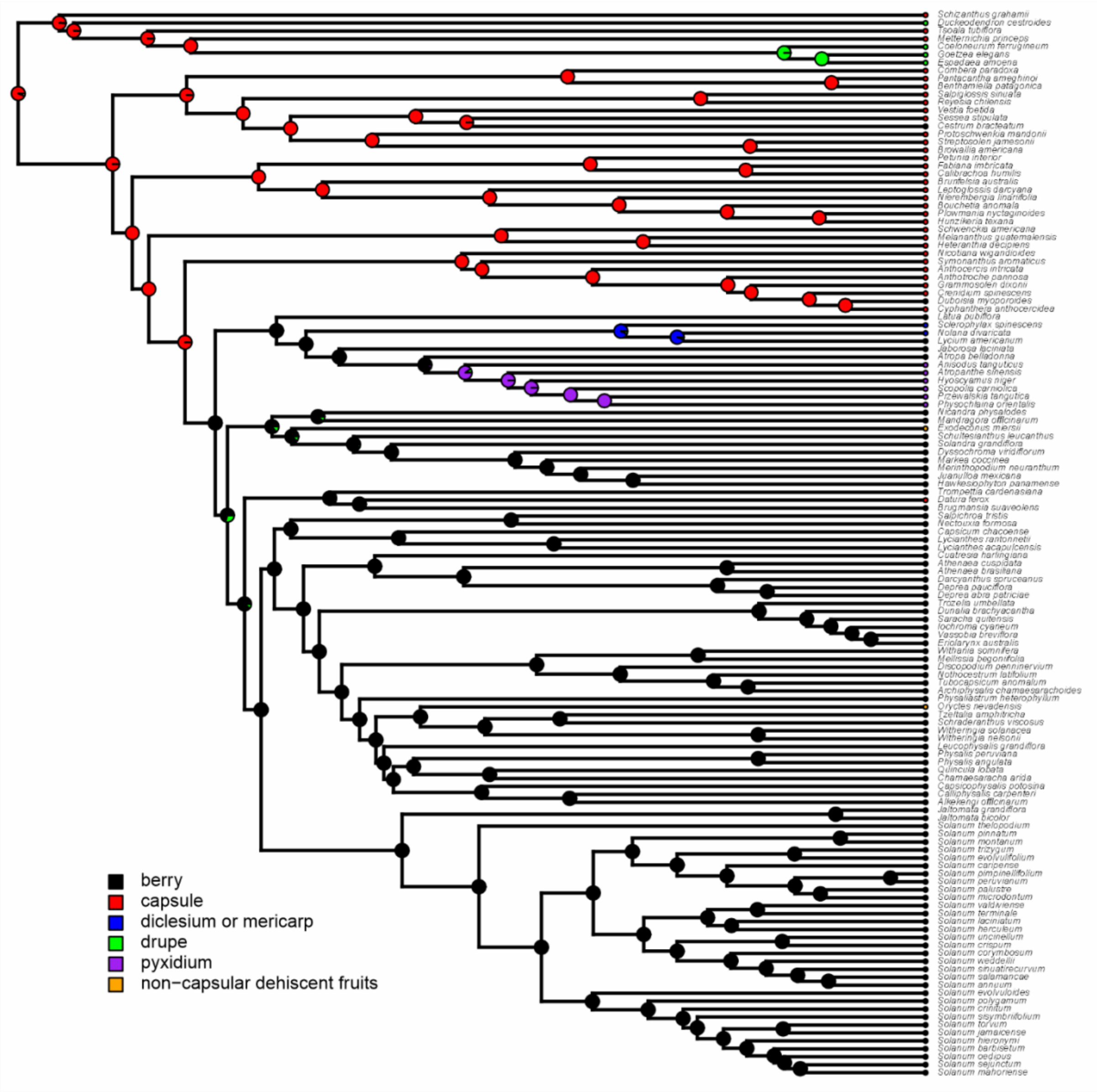
Schizanthus grahamii
Duckeodendron cestroides
Tsoala tubiflora
Metternichia princeps
Coeloneurum ferrugineum
Goetzea elegans
Espadaea amoena
Combera paradoxa
Pantacantha ameghinoi
Benthamiella patagonica
Salpiglossis sinuata
Reyesia chilensis
Vestia foetida
Sessea stipulata
Cestrum bracteatum
Protoschwenkia mandonii
Streptosolen jamesonii
Browallia americana
Petunia interior
Fabiana imbricata
Calibrachoa humilis
Brunfelsia australis
Leptoglossis darcyana
Nierembergia linariifolia
Bouchetia anomala
Plowmania nyctaginoides
Hunzikeria texana
Schwenckia americana
Melananthus guatemalensis
Heteranthia decipiens
Nicotiana wigandioides
Symonanthus aromaticus
Anthocercis intricata
Anthotroche pannosa
Grammosolen dixonii
Crenidium spinescens
Duboisia myoporoides
Cyphanthera anthocercidea
Latua pubiflora
Sclerophylax spinescens
Nolana divaricata
Lycium americanum
Jaborosa laciniata
Atropa belladonna
Anisodus tanguticus
Atropanthe sinensis
Hyoscyamus niger
Scopolia carniolica
Przewalskia tangutica
Physochlaina orientalis
Nicandra physalodes
Mandragora officinarum
Exodeconus miersii
Schultesianthus leucanthus
Solandra grandiflora
Dyssochroma viridiflorum
Markea coccinea
Merinthopodium neuranthum
Juanulloa mexicana
Hawkesiophyton panamense
Trompettia cardenasiana
Datura ferox
Brugmansia suaveolens
Salpichroa tristis
Nectouxia formosa
Capsicum chacoense
Lycianthes rantonnetii
Lycianthes acapulcensis
Cuatresia harlingiana
Athenaea cuspidata
Athenaea brasiliana
Darcyanthus spruceanus
Deprea pauciflora
Deprea abra patriciae
Trozelia umbellata
Dunalia brachyacantha
Saracha quitensis
Iochroma cyaneum
Vassobia breviflora
Eriolarynx australis
Withania somnifera
Mellissia begoniifolia
Discopodium penninervium
Nothocestrum latifolium
Tubocapsicum anomalum
Archiphysalis chamaesarachoides
Physaliastrum heterophyllum
Oryctes nevadensis
Tzeltalia amphitricha
Schraderanthus viscosus
Witheringia solanacea
Witheringia nelsonii
Leucophysalis grandiflora
Physalis peruviana
Physalis angulata
Quincula lobata
Chamaesaracha arida
Capsicophysalis potosina
Calliphysalis carpenteri
Alkekengi officinarum
Jaltomata grandiflora
Jaltomata bicolor
Solanum thelopodium
Solanum pinnatum
Solanum montanum
Solanum trizygum
Solanum evolvulifolium
Solanum caripense
Solanum pimpinellifolium
Solanum peruvianum
Solanum palustre
Solanum microdontum
Solanum valdiviense
Solanum terminale
Solanum laciniatum
Solanum herculeum
Solanum uncinellum
Solanum crispum
Solanum corymbosum
Solanum weddellii
Solanum sinuatirecurvum
Solanum salamancae
Solanum annuum
Solanum evolvuloides
Solanum polygamum
Solanum crinitum
Solanum sisymbriifolium
Solanum torvum
Solanum jamaicense
Solanum hieronymi
Solanum barbisetum
Solanum oedipus
Solanum sejunctum
Solanum mahoriense
berry
capsule
diclesium or mericarp
drupe
pyxidium
non−capsular dehiscent fruits

**N.** Fruiting calyx inflated

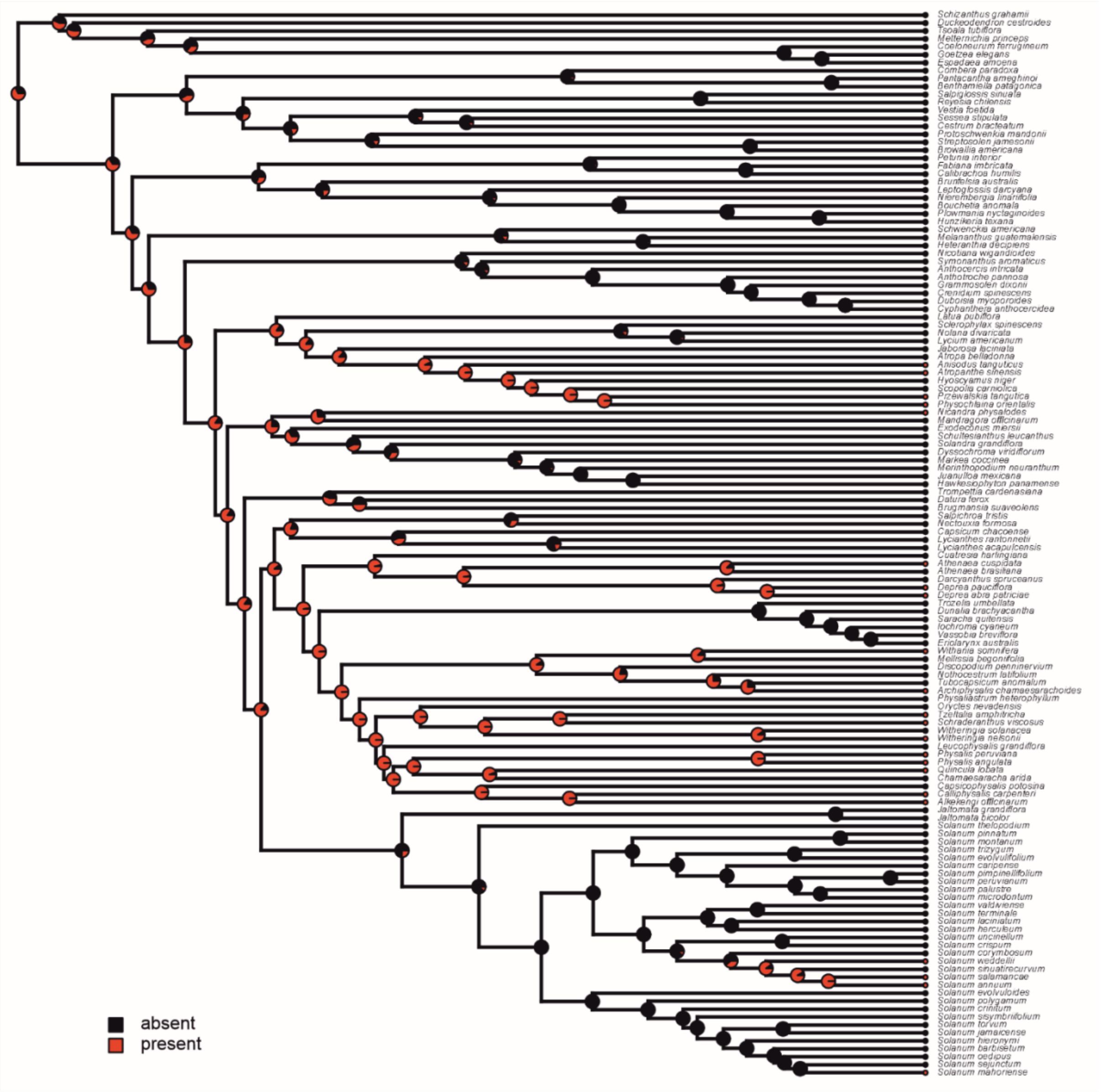

**O.** Fruiting calyx venation pattern

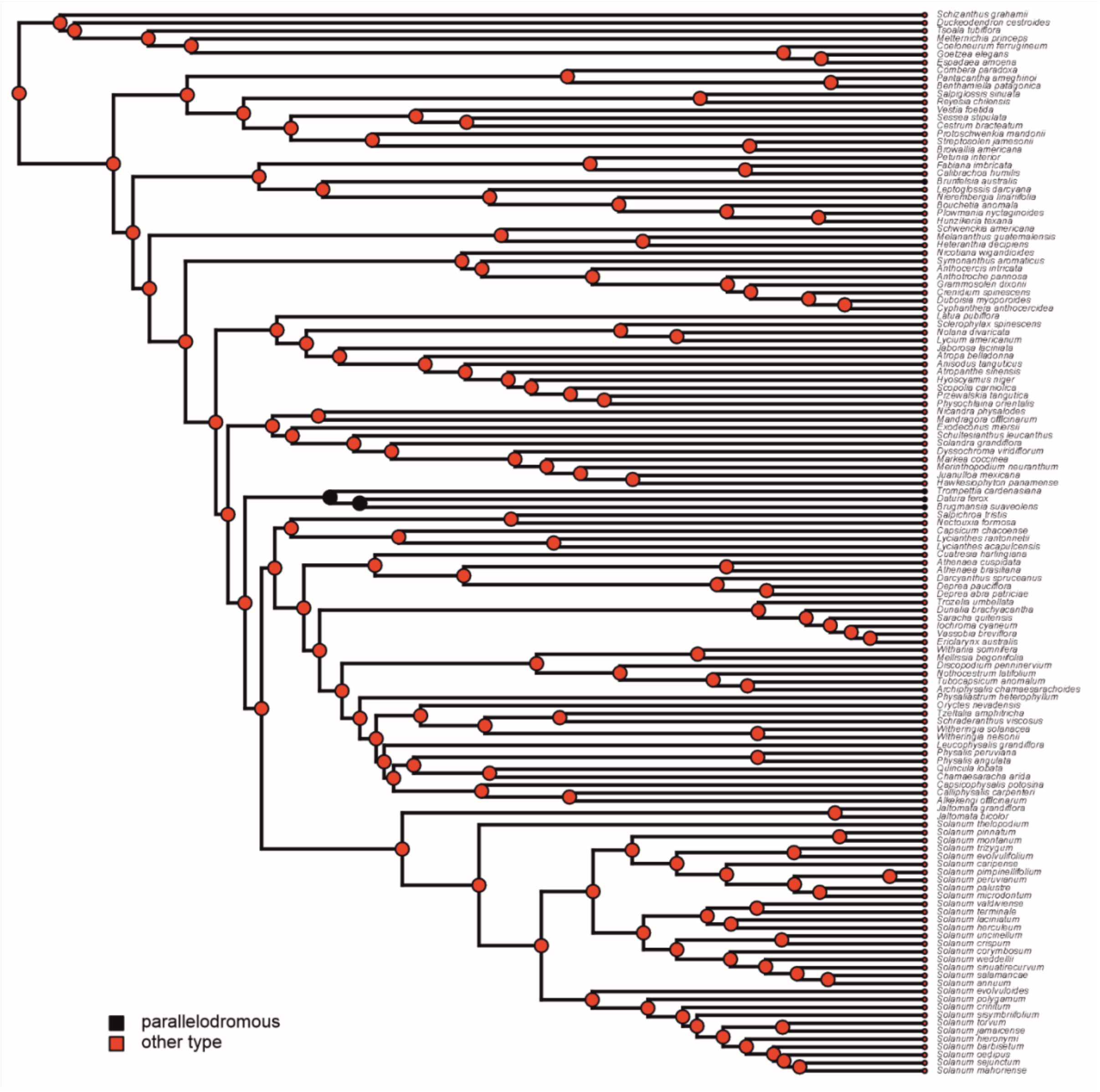

**P.** Fruiting calyx teeth

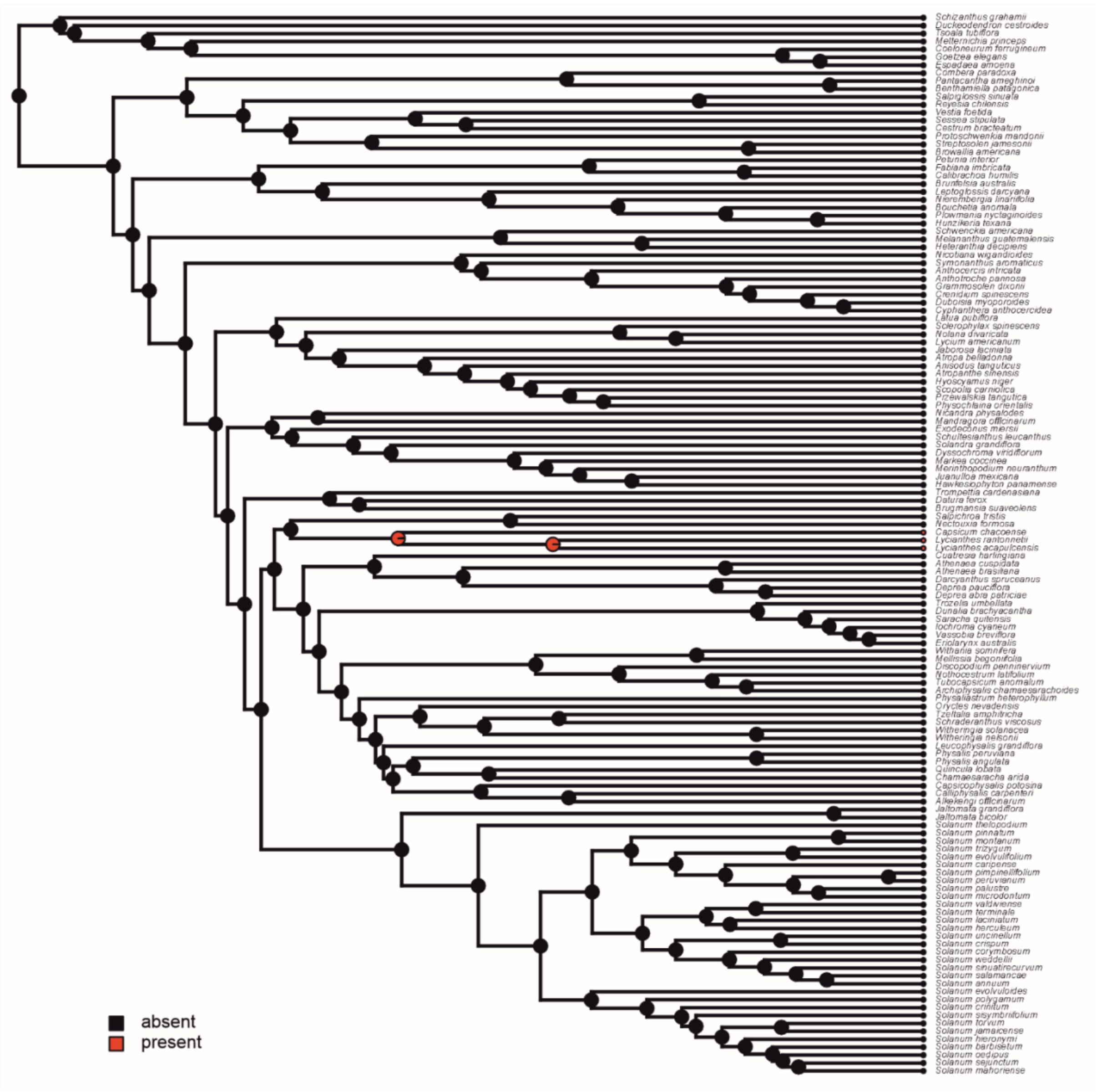

**Figure S9.** MCC tree of Solanaceae inferred from the time-heterogeneous TED model. Unlike Fig. 2 in the main manuscript, this summary tree includes all the fossils, including those with uncertain positions. These conflicting positions, particularly of *Solanum foveolatum* and *Solanum miocenicus* result in irreconcilable nodes in this summary tree.

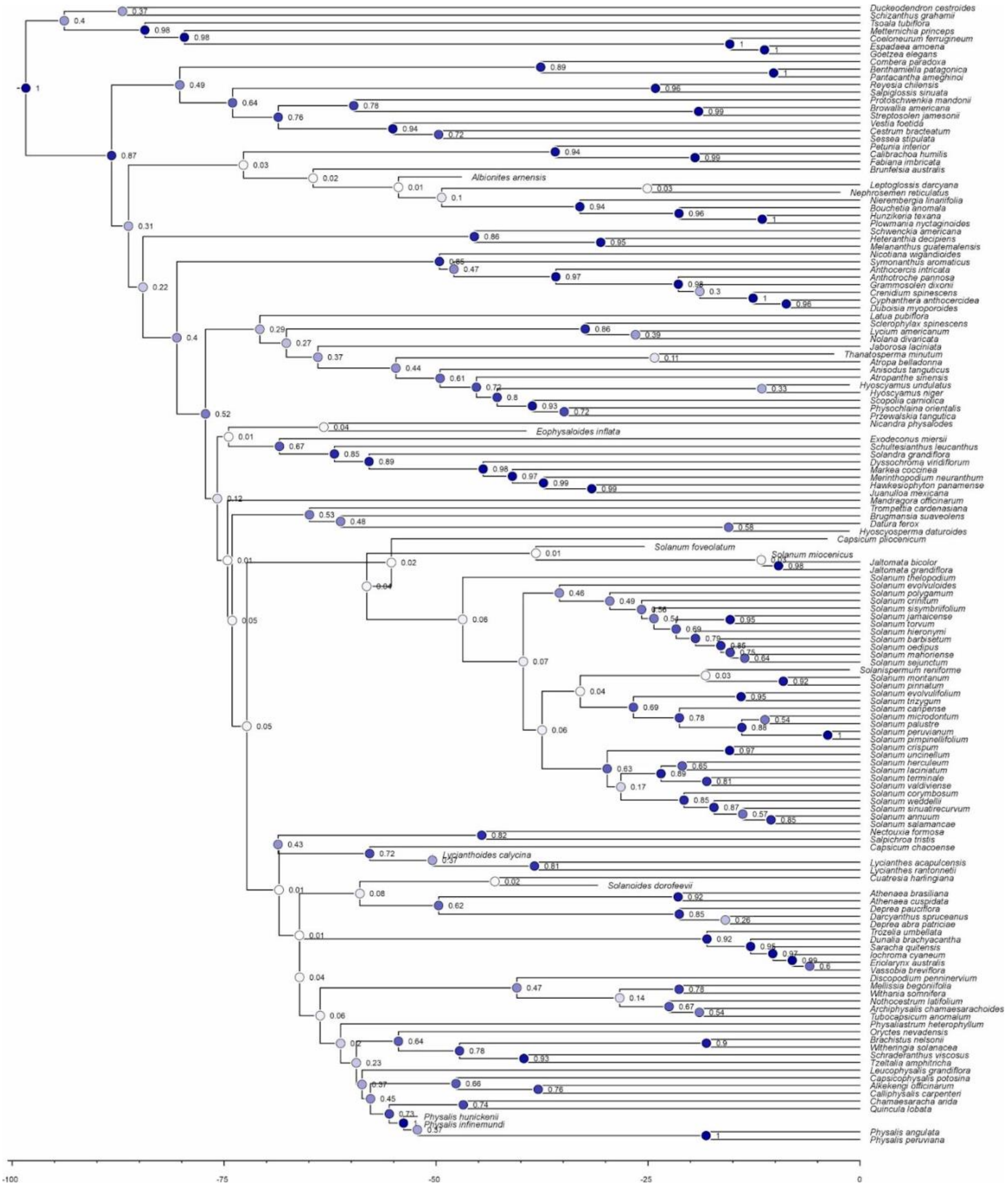

**Figure S10. —`One-fossil MCCs`.** Maximum credibility trees computed by removing all but one fossil at a time from the posterior distribution of topologies of the time-heterogeneous TED analysis.

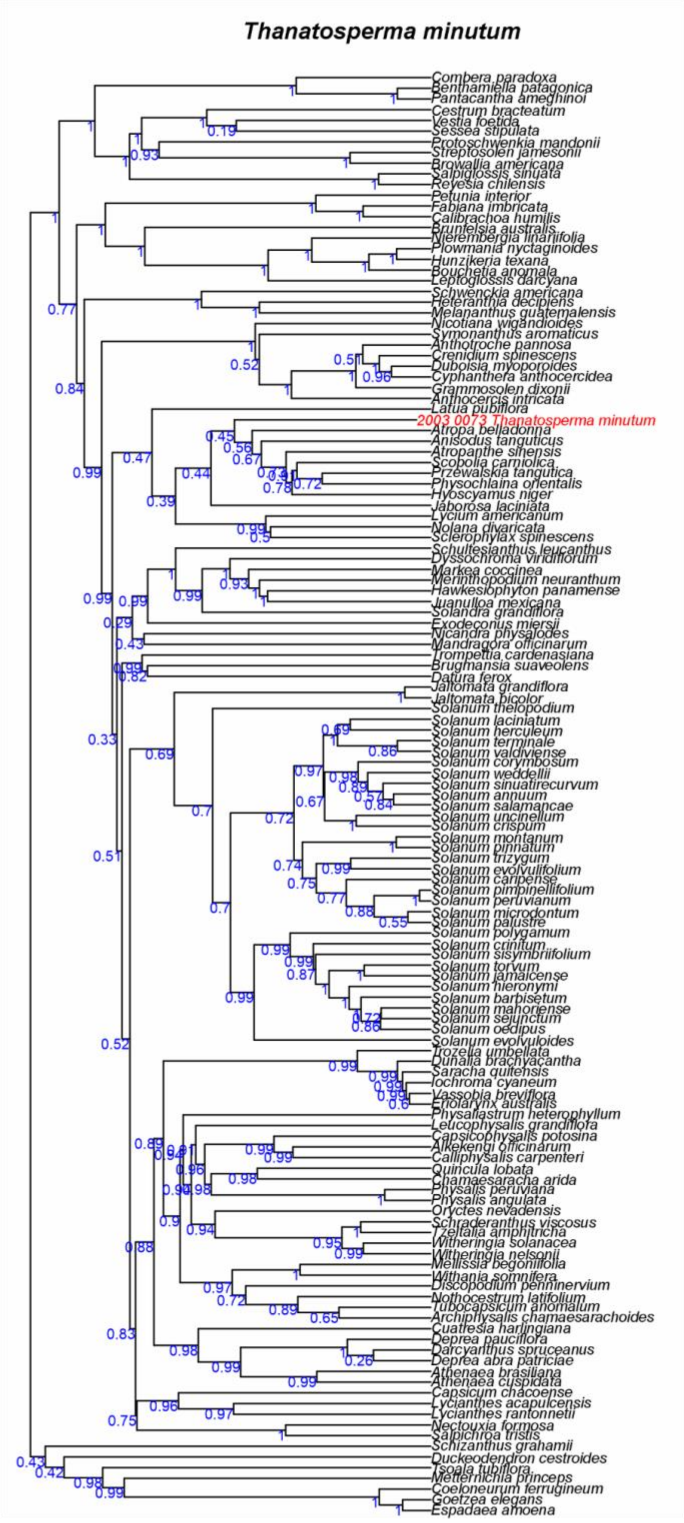

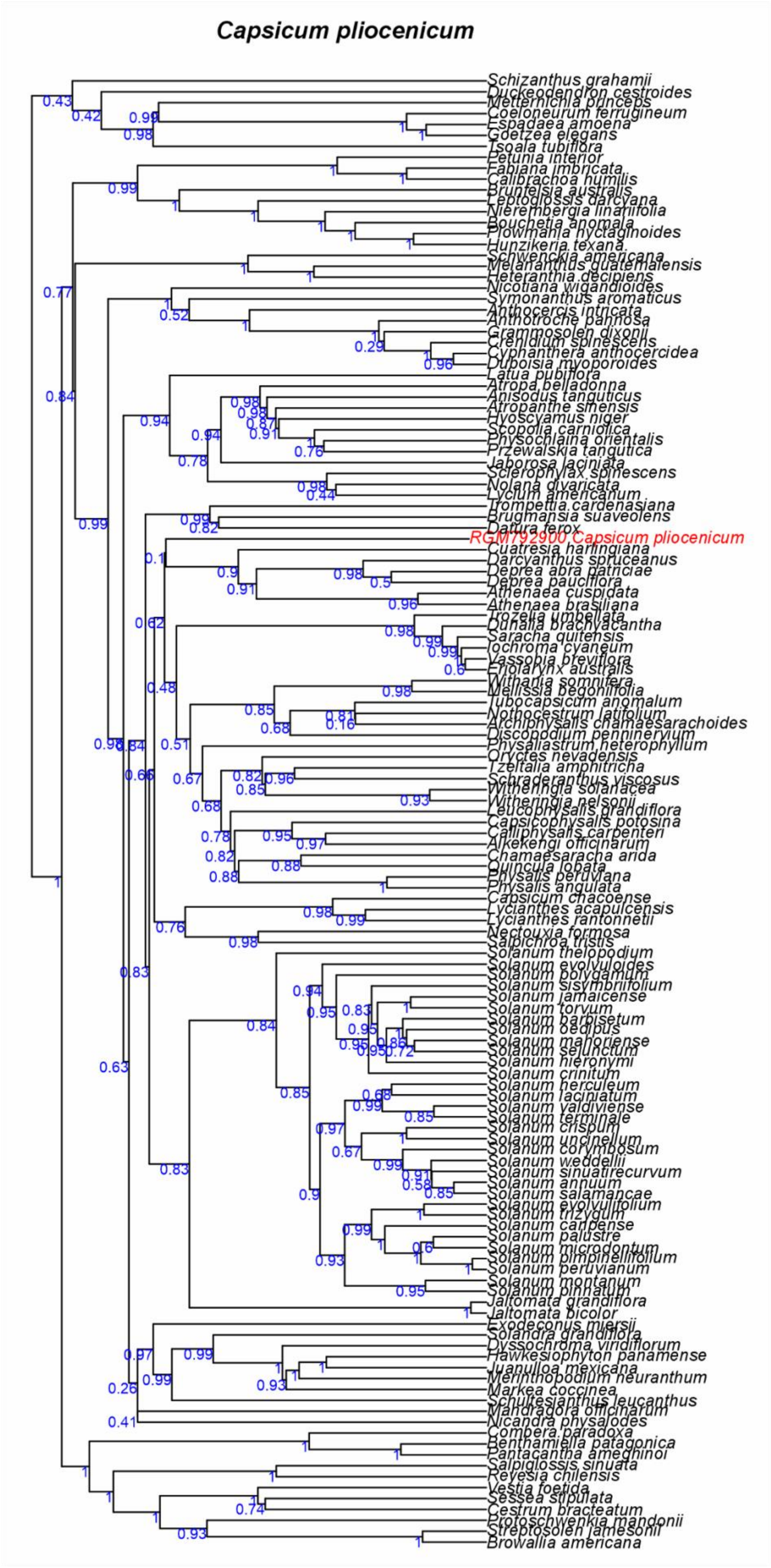
Capsicum pliocenicum
Schizanthus grahamii
Duckeodendron cestroides
Metternichia princeps
Coeloneurum ferrugineum
Espadaea amoena
Goetzea elegans
Tsoala tubiflora
Petunia interior
Fabiana imbricata
Calibrachoa humilis
Brunfelsia australis
Leptoglossis darcyana
Nierembergia linariifolia
Bouchetia anomala
Plowmania nyctaginoides
Hunzikeria texana
Schwenckia americana
Melananthus guatemalensis
Heteranthia decipiens
Nicotiana wigandioides
Symonanthus aromaticus
Anthocercis intricata
Anthotroche pannosa
Grammosolen dixonii
Crenidium spinescens
Cyphanthera anthocercidea
Duboisia myoporoides
Latua pubiflora
Atropa belladonna
Anisodus tanguticus
Atropanthe sinensis
Hyoscyamus niger
Scopolia carniolica
Physochlaina orientalis
Przewalskia tangutica
Jaborosa laciniata
Sclerophylax spinescens
Nolana divaricata
Lycium americanum
Trompettia cardenasiana
Brugmansia suaveolens
Datura ferox
RGM792900 Capsicum pliocenicum
Cuatresia harlingiana
Darcyanthus spruceanus
Deprea abra patriciae
Deprea pauciflora
Athenaea cuspidata
Athenaea brasiliana
Trozelia umbellata
Dunalia brachyacantha
Saracha quitensis
Iochroma cyaneum
Vassobia breviflora
Eriolarynx australis
Withania somnifera
Mellissia begoniifolia
Tubocapsicum anomalum
Nothocestrum latifolium
Archiphysalis chamaesarachoides
Discopodium penninervium
Physaliastrum heterophyllum
Oryctes nevadensis
Tzeltalia amphitricha
Schraderanthus viscosus
Witheringia solanacea
Witheringia nelsonii
Leucophysalis grandiflora
Capsicophysalis potosina
Calliphysalis carpenteri
Alkekengi officinarum
Chamaesaracha arida
Quincula lobata
Physalis peruviana
Physalis angulata
Capsicum chacoense
Lycianthes acapulcensis
Lycianthes rantonnetii
Nectouxia formosa
Salpichroa tristis
Solanum thelopodium
Solanum evolvuloides
Solanum polygamum
Solanum sisymbriifolium
Solanum jamaicense
Solanum torvum
Solanum barbisetum
Solanum oedipus
Solanum mahoriense
Solanum sejunctum
Solanum hieronymi
Solanum crinitum
Solanum herculeum
Solanum laciniatum
Solanum valdiviense
Solanum terminale
Solanum crispum
Solanum uncinellum
Solanum corymbosum
Solanum weddellii
Solanum sinuatirecurvum
Solanum annuum
Solanum salamancae
Solanum evolvulifolium
Solanum trizygum
Solanum caripense
Solanum palustre
Solanum microdontum
Solanum pimpinellifolium
Solanum peruvianum
Solanum montanum
Solanum pinnatum
Jaltomata grandiflora
Jaltomata bicolor
Exodeconus miersii
Solandra grandiflora
Dyssochroma viridiflorum
Hawkesiophyton panamense
Juanulloa mexicana
Merinthopodium neuranthum
Markea coccinea
Schultesianthus leucanthus
Mandragora officinarum
Nicandra physalodes
Combera paradoxa
Benthamiella patagonica
Pantacantha ameghinoi
Salpiglossis sinuata
Reyesia chilensis
Vestia foetida
Sessea stipulata
Cestrum bracteatum
Protoschwenkia mandonii
Streptosolen jamesonii
Browallia americana

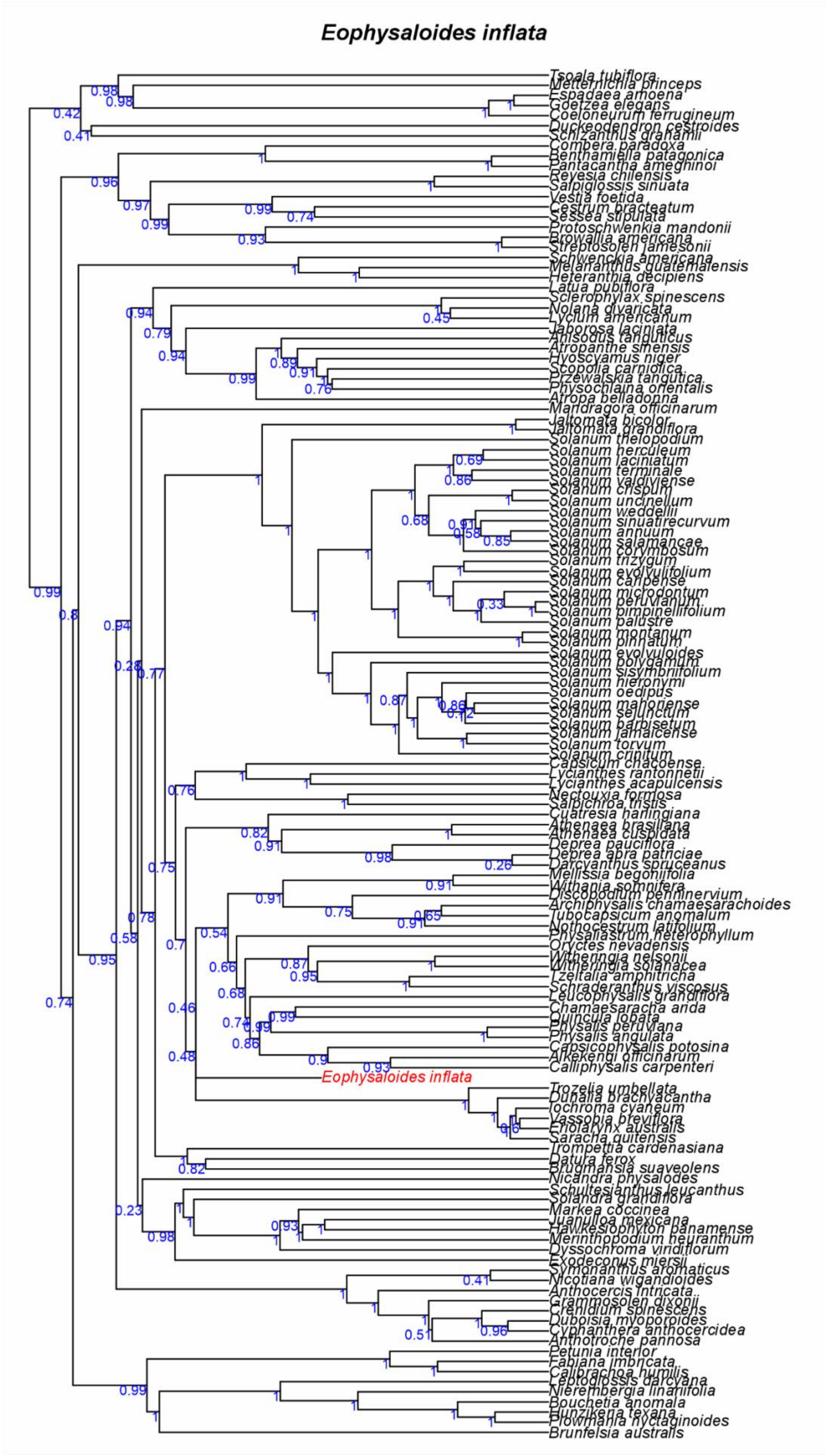
Eophysaloides inflata
Tsoala tubiflora
Metternichia princeps
Espadaea amoena
Goetzea elegans
Coeloneurum ferrugineum
Duckeodendron cestroides
Schizanthus grahamii
Combera paradoxa
Benthamiella patagonica
Pantacantha ameghinoi
Reyesia chilensis
Salpiglossis sinuata
Vestia foetida
Cestrum bracteatum
Sessea stipulata
Protoschwenkia mandonii
Browallia americana
Streptosolen jamesonii
Schwenckia americana
Melananthus guatemalensis
Heteranthia decipiens
Latua pubiflora
Sclerophylax spinescens
Nolana divaricata
Lycium americanum
Jaborosa laciniata
Anisodus tanguticus
Atropanthe sinensis
Hyoscyamus niger
Scopolia carniolica
Przewalskia tangutica
Physochlaina orientalis
Atropa belladonna
Mandragora officinarum
Jaltomata bicolor
Jaltomata grandiflora
Solanum thelopodium
Solanum herculeum
Solanum laciniatum
Solanum terminale
Solanum valdiviense
Solanum crispum
Solanum uncinellum
Solanum weddellii
Solanum sinuatirecurvum
Solanum annuum
Solanum salamancae
Solanum corymbosum
Solanum trizygum
Solanum evolvulifolium
Solanum caripense
Solanum microdontum
Solanum peruvianum
Solanum pimpinellifolium
Solanum palustre
Solanum montanum
Solanum pinnatum
Solanum evolvuloides
Solanum polygamum
Solanum sisymbriifolium
Solanum hieronymi
Solanum oedipus
Solanum mahoriense
Solanum sejunctum
Solanum barbisetum
Solanum jamaicense
Solanum torvum
Solanum crinitum
Capsicum chacoense
Lycianthes rantonnetii
Lycianthes acapulcensis
Nectouxia formosa
Salpichroa tristis
Cuatresia harlingiana
Athenaea brasiliana
Athenaea cuspidata
Deprea pauciflora
Deprea abra patriciae
Darcyanthus spruceanus
Mellissia begoniifolia
Withania somnifera
Discopodium penninervium
Archiphysalis chamaesarachoides
Tubocapsicum anomalum
Nothocestrum latifolium
Physaliastrum heterophyllum
Oryctes nevadensis
Witheringia nelsonii
Witheringia solanacea
Tzeltalia amphitricha
Schraderanthus viscosus
Leucophysalis grandiflora
Chamaesaracha arida
Quincula lobata
Physalis peruviana
Physalis angulata
Capsicophysalis potosina
Alkekengi officinarum
Calliphysalis carpenteri
Eophysaloides inflata
Trozelia umbellata
Dunalia brachyacantha
Iochroma cyaneum
Vassobia breviflora
Eriolarynx australis
Saracha quitensis
Trompettia cardenasiana
Datura ferox
Brugmansia suaveolens
Nicandra physalodes
Schultesianthus leucanthus
Solandra grandiflora
Markea coccinea
Juanulloa mexicana
Hawkesiophyton panamense
Merinthopodium neuranthum
Dyssochroma viridiflorum
Exodeconus miersii
Symonanthus aromaticus
Nicotiana wigandioides
Anthocercis intricata
Grammosolen dixonii
Crenidium spinescens
Duboisia myoporoides
Cyphanthera anthocercidea
Anthotroche pannosa
Petunia interior
Fabiana imbricata
Calibrachoa humilis
Leptoglossis darcyana
Nierembergia linariifolia
Bouchetia anomala
Hunzikeria texana
Plowmania nyctaginoides
Brunfelsia australis

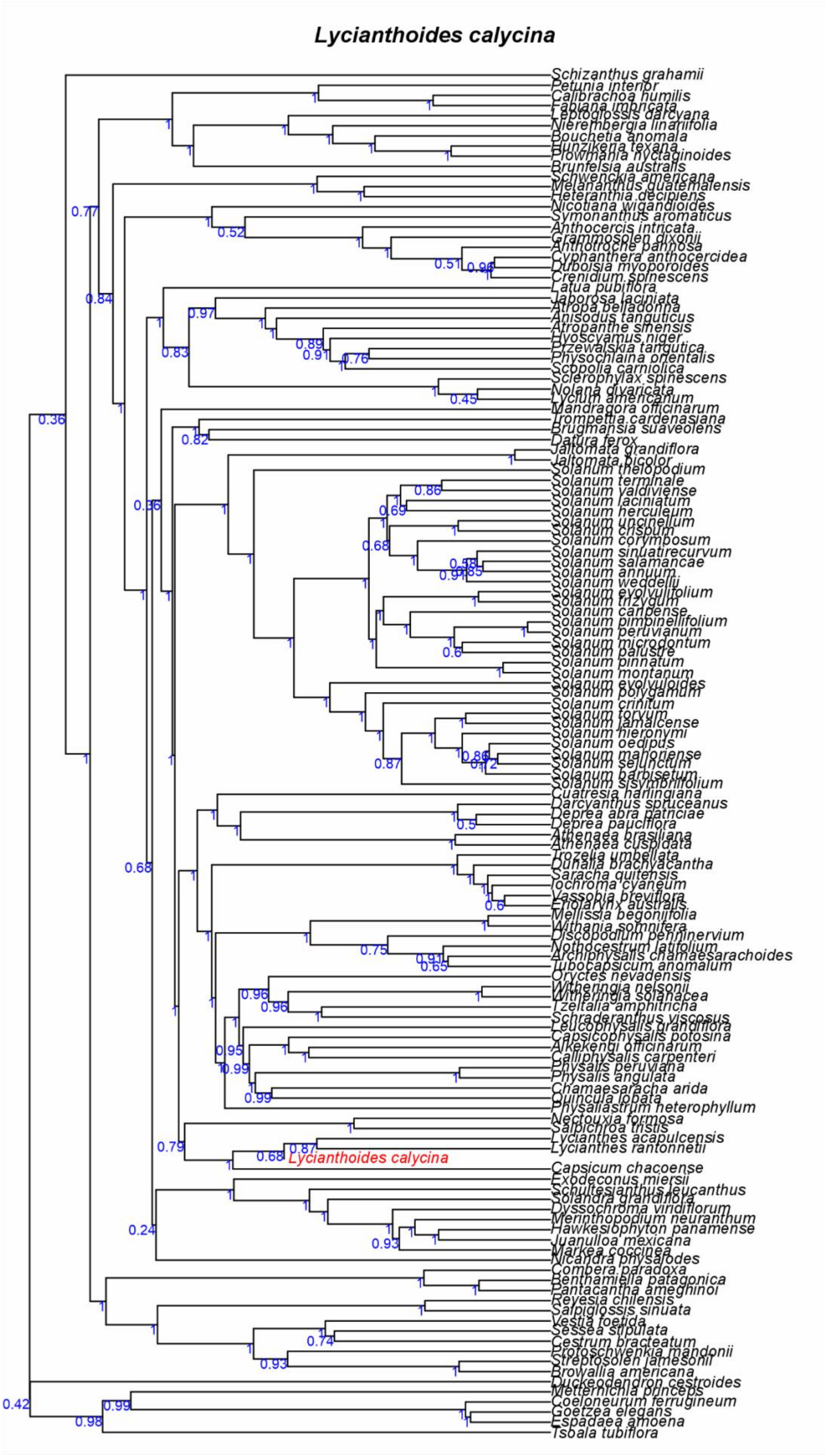
Lycianthoides calycina
Schizanthus grahamii
Petunia interior
Calibrachoa humilis
Fabiana imbricata
Leptoglossis darcyana
Nierembergia linariifolia
Bouchetia anomala
Hunzikeria texana
Plowmania nyctaginoides
Brunfelsia australis
Schwenckia americana
Melananthus guatemalensis
Heteranthia decipiens
Nicotiana wigandioides
Symonanthus aromaticus
Anthocercis intricata
Grammosolen dixonii
Anthotroche pannosa
Cyphanthera anthocercidea
Duboisia myoporoides
Crenidium spinescens
Latua pubiflora
Jaborosa laciniata
Atropa belladonna
Anisodus tanguticus
Atropanthe sinensis
Hyoscyamus niger
Przewalskia tangutica
Physochlaina orientalis
Scopolia carniolica
Sclerophylax spinescens
Nolana divaricata
Lycium americanum
Mandragora officinarum
Trompettia cardenasiana
Brugmansia suaveolens
Datura ferox
Jaltomata grandiflora
Jaltomata bicolor
Solanum theiopodium
Solanum terminale
Solanum valdiviense
Solanum laciniatum
Solanum herculeum
Solanum uncinellum
Solanum crispum
Solanum corymbosum
Solanum sinuatirecurvum
Solanum salamancae
Solanum annuum
Solanum weddellii
Solanum evolvulifolium
Solanum trizygum
Solanum caripense
Solanum pimpinellifolium
Solanum peruvianum
Solanum microdontum
Solanum palustre
Solanum pinnatum
Solanum montanum
Solanum evolvuloides
Solanum polygamum
Solanum crinitum
Solanum torvum
Solanum jamaicense
Solanum hieronymi
Solanum oedipus
Solanum mahoriense
Solanum sejunctum
Solanum barbisetum
Solanum sisymbriifolium
Cuatresia harlingiana
Darcyanthus spruceanus
Deprea abra patriciae
Deprea pauciflora
Athenaea brasiliana
Athenaea cuspidata
Trozelia umbellata
Dunalia brachyacantha
Saracha quitensis
Iochroma cyaneum
Vassobia breviflora
Eriolarynx australis
Mellissia begoniifolia
Withania somnifera
Discopodium penninervium
Nothocestrum latifolium
Archiphysalis chamaesarachoides
Tubocapsicum anomalum
Oryctes nevadensis
Witheringia nelsonii
Witheringia solanacea
Tzeltalia amphitricha
Schraderanthus viscosus
Leucophysalis grandiflora
Capsicophysalis potosina
Alkekengi officinarum
Calliphysalis carpenteri
Physalis peruviana
Physalis angulata
Chamaesaracha arida
Quincula lobata
Physaliastrum heterophyllum
Nectouxia formosa
Salpichroa tristis
Lycianthes acapulcensis
Lycianthes rantonnetii
Lycianthoides calycina
Capsicum chacoense
Exodeconus miersii
Schultesianthus leucanthus
Solandra grandiflora
Dyssochroma viridiflorum
Merinthopodium neuranthum
Hawkesiophyton panamense
Juanulloa mexicana
Markea coccinea
Nicandra physalodes
Combera paradoxa
Benthamiella patagonica
Pantacantha ameghinoi
Reyesia chilensis
Salpiglossis sinuata
Vestia foetida
Sessea stipulata
Cestrum bracteatum
Protoschwenkia mandonii
Streptosolen jamesonii
Browallia americana
Duckeodendron cestroides
Metternichia princeps
Coeloneurum ferrugineum
Goetzea elegans
Espadaea amoena
Tsoala tubiflora

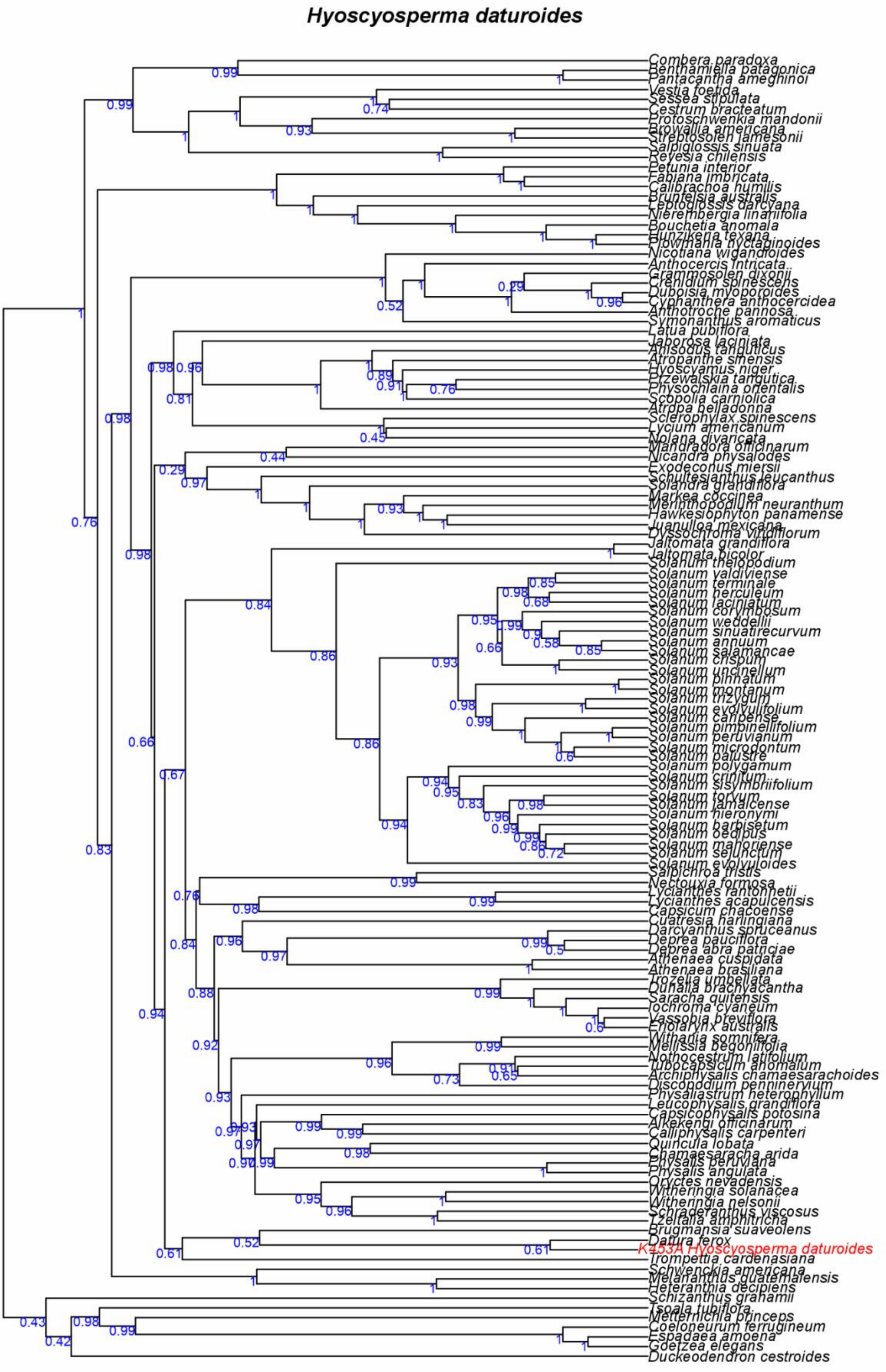
Hyoscyosperma daturoides
Combera paradoxa
Benthamiella patagonica
Pantacantha ameghinoi
Vestia foetida
Sessea stipulata
Cestrum bracteatum
Protoschwenkia mandonii
Browallia americana
Streptosolen jamesonii
Salpiglossis sinuata
Reyesia chilensis
Petunia interior
Fabiana imbricata
Calibrachoa humilis
Brunfelsia australis
Leptoglossis darcyana
Nierembergia linariifolia
Bouchetia anomala
Hunzikeria texana
Plowmania nyctaginoides
Nicotiana wigandioides
Anthocercis intricata
Grammosolen dixonii
Crenidium spinescens
Duboisia myoporoides
Cyphanthera anthocercidea
Anthotroche pannosa
Symonanthus aromaticus
Latua pubiflora
Jaborosa laciniata
Anisodus tanguticus
Atropanthe sinensis
Hyoscyamus niger
Przewalskia tangutica
Physochlaina orientalis
Scopolia carniolica
Atropa belladonna
Sclerophylax spinescens
Lycium americanum
Nolana divaricata
Mandragora officinarum
Nicandra physalodes
Exodeconus miersii
Schultesianthus leucanthus
Solandra grandiflora
Markea coccinea
Merinthopodium neuranthum
Hawkesiophyton panamense
Juanulloa mexicana
Dyssochroma viridiflorum
Jaltomata grandiflora
Jaltomata bicolor
Solanum thelopodium
Solanum valdiviense
Solanum terminale
Solanum herculeum
Solanum laciniatum
Solanum corymbosum
Solanum weddellii
Solanum sinuatirecurvum
Solanum annuum
Solanum salamancae
Solanum crispum
Solanum uncinellum
Solanum pinnatum
Solanum montanum
Solanum trizygum
Solanum evolvulifolium
Solanum caripense
Solanum pimpinellifolium
Solanum peruvianum
Solanum microdontum
Solanum palustre
Solanum polygamum
Solanum crinitum
Solanum sisymbriifolium
Solanum torvum
Solanum jamaicense
Solanum hieronymi
Solanum barbisetum
Solanum oedipus
Solanum mahoriense
Solanum sejunctum
Solanum evolvuloides
Salpichroa tristis
Nectouxia formosa
Lycianthes rantonnetii
Lycianthes acapulcensis
Capsicum chacoense
Cuatresia harlingiana
Darcyanthus spruceanus
Deprea pauciflora
Deprea abra patriciae
Athenaea cuspidata
Athenaea brasiliana
Trozelia umbellata
Dunalia brachyacantha
Saracha quitensis
Iochroma cyaneum
Vassobia breviflora
Eriolarynx australis
Withania somnifera
Mellissia begoniifolia
Nothocestrum latifolium
Tubocapsicum anomalum
Archiphysalis chamaesarachoides
Discopodium penninervium
Physaliastrum heterophyllum
Leucophysalis grandiflora
Capsicophysalis potosina
Alkekengi officinarum
Calliphysalis carpenteri
Quincula lobata
Chamaesaracha arida
Physalis peruviana
Physalis angulata
Oryctes nevadensis
Witheringia solanacea
Witheringia nelsonii
Schraderanthus viscosus
Tzeltalia amphitricha
Brugmansia suaveolens
Datura ferox
K453A Hyoscyosperma daturoides
Trompettia cardenasiana
Schwenckia americana
Melananthus guatemalensis
Heteranthia decipiens
Schizanthus grahamii
Tsoala tubiflora
Metternichia princeps
Coeloneurum ferrugineum
Espadaea amoena
Goetzea elegans
Duckeodendron cestroides

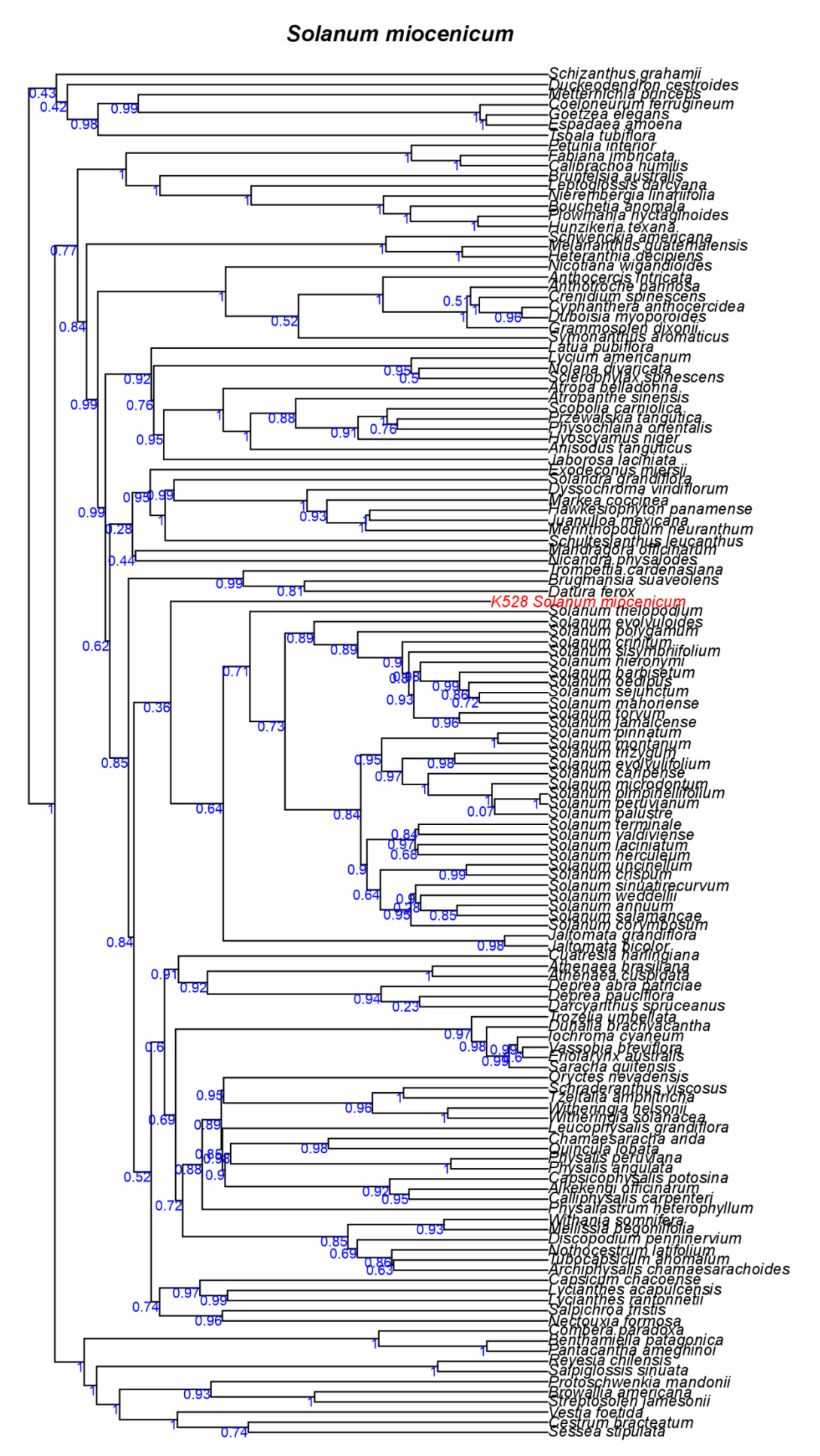

Solanum miocenicum
K528 Solanum miocenicum

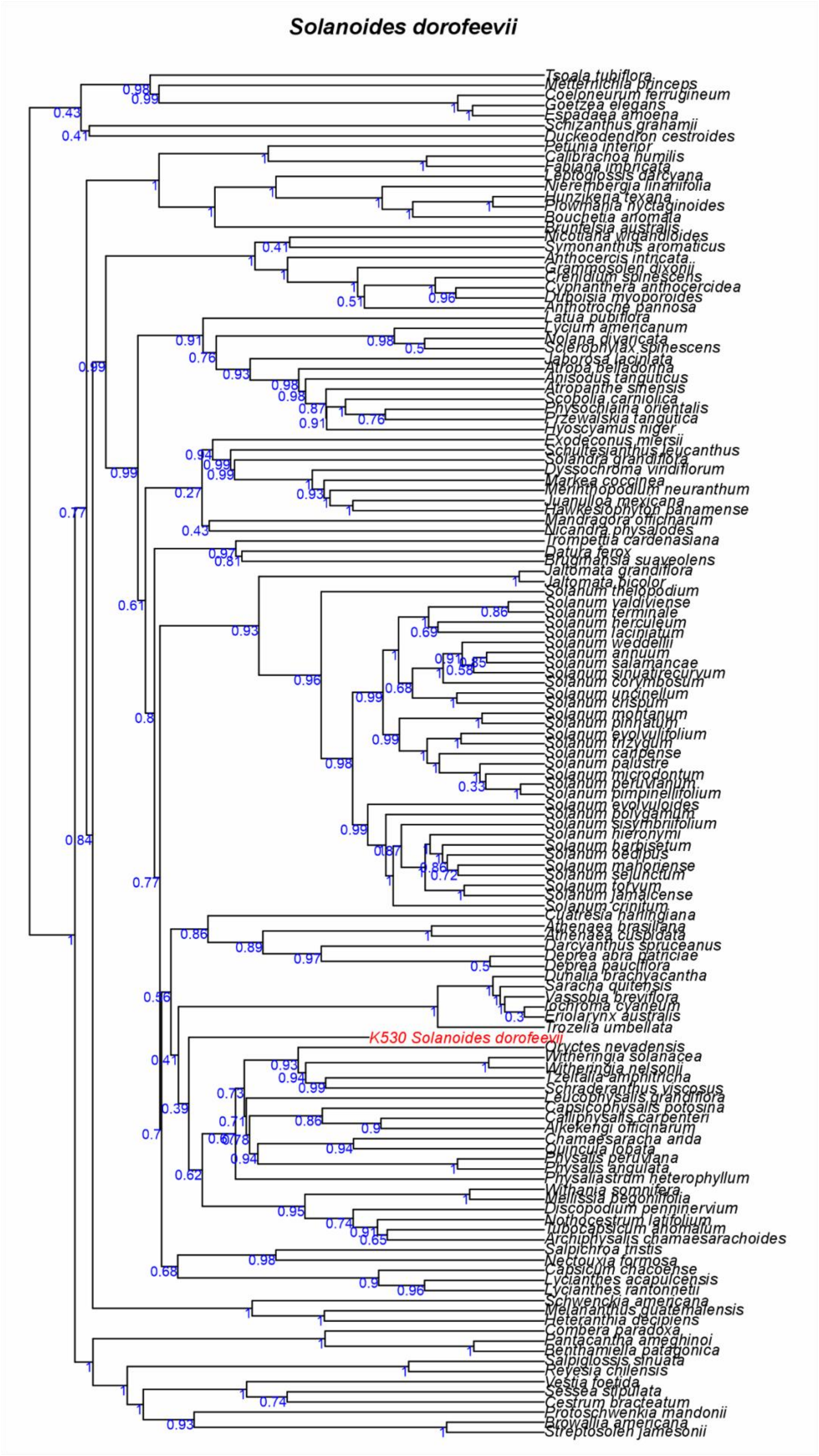
Solanoides dorofeevii
Tsoala tubiflora
Metternichia princeps
Coeloneurum ferrugineum
Goetzea elegans
Espadaea amoena
Schizanthus grahamii
Duckeodendron cestroides
Petunia interior
Calibrachoa humilis
Fabiana imbricata
Leptoglossis darcyana
Nierembergia linariifolia
Hunzikeria texana
Plowmania nyctaginoides
Bouchetia anomala
Brunfelsia australis
Nicotiana wigandioides
Symonanthus aromaticus
Anthocercis intricata
Grammosolen dixonii
Crenidium spinescens
Cyphanthera anthocercidea
Duboisia myoporoides
Anthotroche pannosa
Latua pubiflora
Lycium americanum
Nolana divaricata
Sclerophylax spinescens
Jaborosa laciniata
Atropa belladonna
Anisodus tanguticus
Atropanthe sinensis
Scopolia carniolica
Physochlaina orientalis
Przewalskia tangutica
Hyoscyamus niger
Exodeconus miersii
Schultesianthus leucanthus
Solandra grandiflora
Dyssochroma viridiflorum
Markea coccinea
Merinthopodium neuranthum
Juanulloa mexicana
Hawkesiophyton panamense
Mandragora officinarum
Nicandra physalodes
Trompettia cardenasiana
Datura ferox
Brugmansia suaveolens
Jaltomata grandiflora
Jaltomata bicolor
Solanum thelopodium
Solanum valdiviense
Solanum terminale
Solanum herculeum
Solanum laciniatum
Solanum weddellii
Solanum annuum
Solanum salamancae
Solanum sinuatirecurvum
Solanum corymbosum
Solanum uncinellum
Solanum crispum
Solanum montanum
Solanum pinnatum
Solanum evolvulifolium
Solanum trizygum
Solanum caripense
Solanum palustre
Solanum microdontum
Solanum peruvianum
Solanum pimpinellifolium
Solanum evolvuloides
Solanum polygamum
Solanum sisymbriifolium
Solanum hieronymi
Solanum barbisetum
Solanum oedipus
Solanum mahoriense
Solanum sejunctum
Solanum torvum
Solanum jamaicense
Solanum crinitum
Cuatresia harlingiana
Athenaea brasiliana
Athenaea cuspidata
Darcyanthus spruceanus
Deprea abra patriciae
Deprea pauciflora
Dunalia brachyacantha
Saracha quitensis
Vassobia breviflora
Iochroma cyaneum
Eriolarynx australis
Trozelia umbellata
K530 Solanoides dorofeevii
Oryctes nevadensis
Witheringia solanacea
Witheringia nelsonii
Tzeltalia amphitricha
Schraderanthus viscosus
Leucophysalis grandiflora
Capsicophysalis potosina
Calliphysalis carpenteri
Alkekengi officinarum
Chamaesaracha arida
Quincula lobata
Physalis peruviana
Physalis angulata
Physaliastrum heterophyllum
Withania somnifera
Mellissia begoniifolia
Discopodium penninervium
Nothocestrum latifolium
Tubocapsicum anomalum
Archiphysalis chamaesarachoides
Salpichroa tristis
Nectouxia formosa
Capsicum chacoense
Lycianthes acapulcensis
Lycianthes rantonnetii
Schwenckia americana
Melananthus guatemalensis
Heteranthia decipiens
Combera paradoxa
Pantacantha ameghinoi
Benthamiella patagonica
Salpiglossis sinuata
Reyesia chilensis
Vestia foetida
Sessea stipulata
Cestrum bracteatum
Protoschwenkia mandonii
Browallia americana
Streptosolen jamesonii

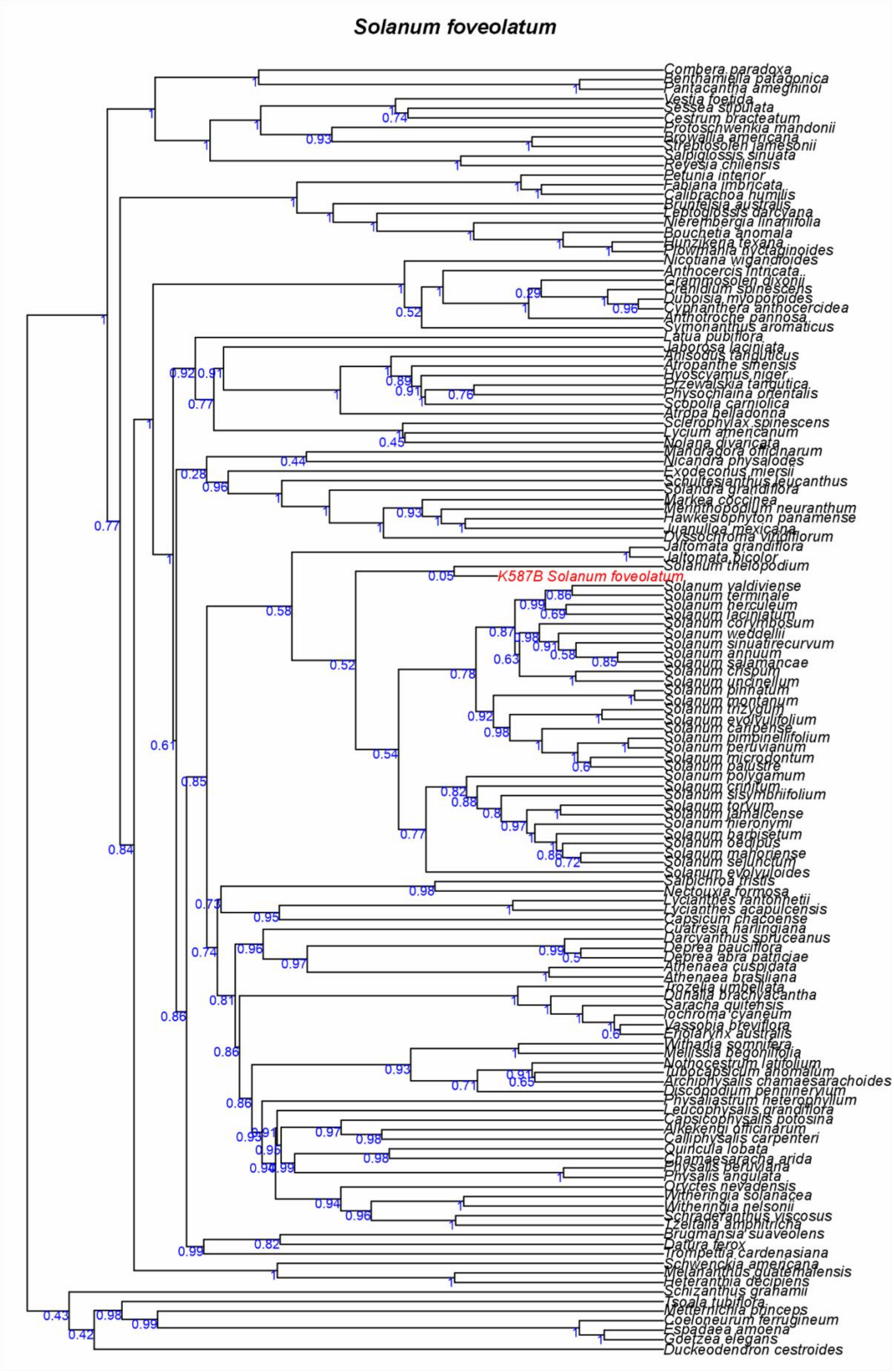
Solanum foveolatum
K587B Solanum foveolatum
Combera paradoxa
Benthamiella patagonica
Pantacantha ameghinoi
Vestia foetida
Sessea stipulata
Cestrum bracteatum
Protoschwenkia mandonii
Browallia americana
Streptosolen jamesonii
Salpiglossis sinuata
Reyesia chilensis
Petunia interior
Fabiana imbricata
Calibrachoa humilis
Brunfelsia australis
Nicotiana wigandioides
Anthocercis intricata
Symonanthus aromaticus
Latua pubiflora
Jaborosa laciniata
Anisodus tanguticus
Atropanthe sinensis
Hyoscyamus niger
Physochlaina orientalis
Scopolia carniolica
Atropa belladonna
Sclerophylax spinescens
Lycium americanum
Nolana divaricata
Mandragora officinarum
Nicandra physalodes
Exodeconus miersii
Schultesianthus leucanthus
Solandra grandiflora
Markea coccinea
Merinthopodium neuranthum
Hawkesiophyton panamense
Juanulloa mexicana
Dyssochroma viridiflorum
Jaltomata grandiflora
Jaltomata bicolor
Solanum thelopodium
Solanum valdiviense
Solanum terminale
Solanum herculeum
Solanum laciniatum
Solanum corymbosum
Solanum weddellii
Solanum sinuatirecurvum
Solanum annuum
Solanum salamancae
Solanum crispum
Solanum uncinellum
Solanum pinnatum
Solanum montanum
Solanum evolvulifolium
Solanum pimpinellifolium
Solanum peruvianum
Solanum microdontum
Solanum palustre
Solanum polygamum
Solanum crinitum
Solanum sisymbriifolium
Solanum torvum
Solanum jamaicense
Solanum hieronymi
Solanum barbisetum
Solanum oedipus
Solanum mahoriense
Solanum evolvuloides
Nectouxia formosa
Lycianthes acapulcensis
Capsicum chacoense
Cuatresia harlingiana
Darcyanthus spruceanus
Deprea pauciflora
Athenaea cuspidata
Athenaea brasiliana
Dunalia brachyacantha
Saracha quitensis
Iochroma cyaneum
Vassobia breviflora
Eriolarynx australis
Withania somnifera
Mellissia begoniifolia
Nothocestrum latifolium
Tubocapsicum anomalum
Archiphysalis chamaesarachoides
Discopodium penninervium
Physaliastrum heterophyllum
Leucophysalis grandiflora
Capsicophysalis potosina
Alkekengi officinarum
Calliphysalis carpenteri
Quincula lobata
Chamaesaracha arida
Physalis peruviana
Physalis angulata
Oryctes nevadensis
Witheringia solanacea
Witheringia nelsonii
Schraderanthus viscosus
Brugmansia suaveolens
Datura ferox
Trompettia cardenasiana
Schwenckia americana
Melananthus guatemalensis
Heteranthia decipiens
Schizanthus grahamii
Tsoala tubiflora
Metternichia princeps
Coeloneurum ferrugineum
Espadaea amoena
Goetzea elegans
Duckeodendron cestroides

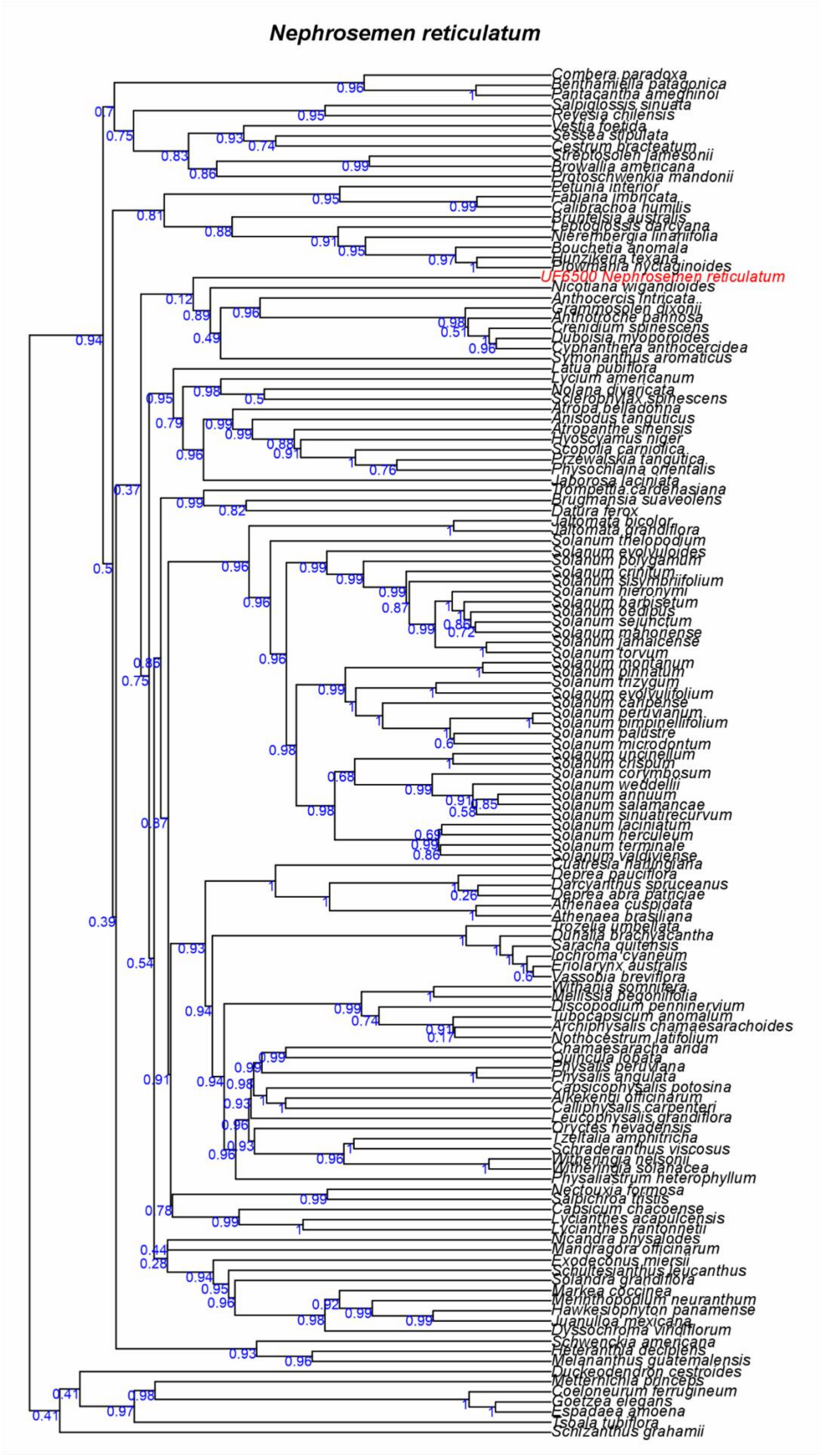
Nephrosemen reticulatum
UF6500 Nephrosemen reticulatum

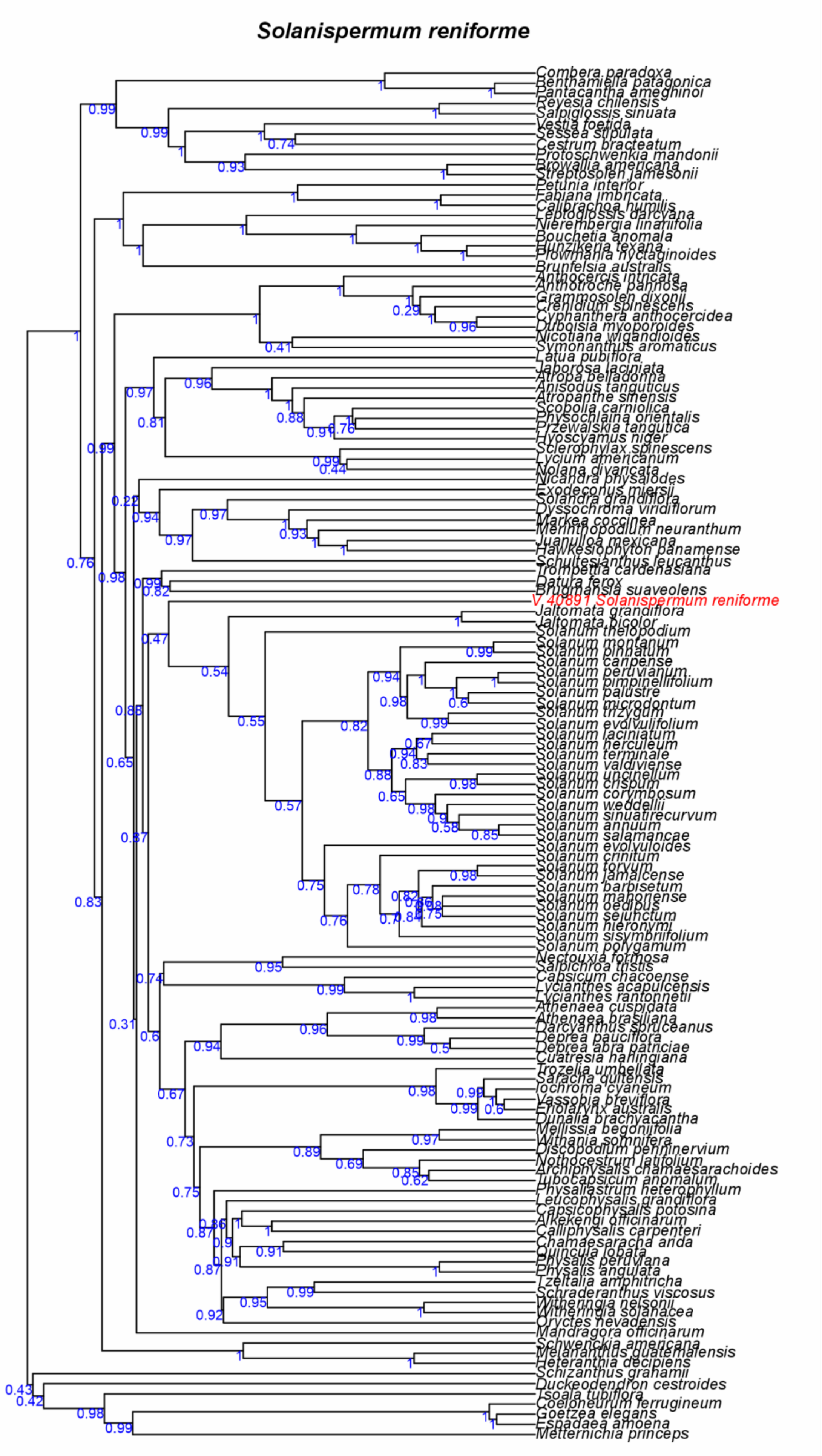

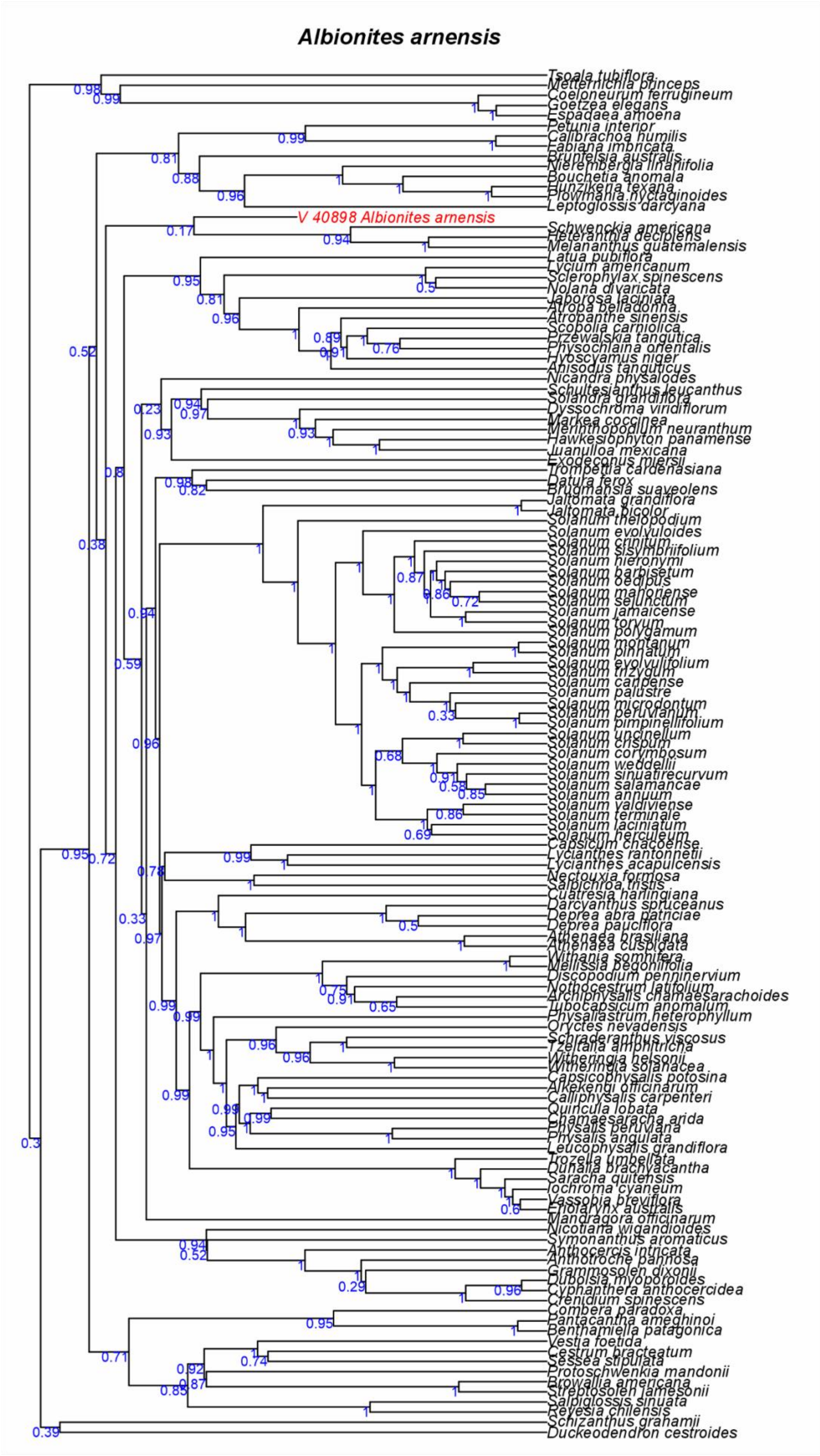
Albionites arnensis
Tsoala tubiflora
Metternichia princeps
Coeloneurum ferrugineum
Goetzea elegans
Espadaea amoena
Petunia interior
Calibrachoa humilis
Fabiana imbricata
Brunfelsia australis
Nierembergia linariifolia
Bouchetia anomala
Hunzikeria texana
Plowmania nyctaginoides
Leptoglossis darcyana
V 40898 Albionites arnensis
Schwenckia americana
Heteranthia decipiens
Melananthus guatemalensis
Latua pubiflora
Lycium americanum
Sclerophylax spinescens
Nolana divaricata
Jaborosa laciniata
Atropa belladonna
Atropanthe sinensis
Scopolia carniolica
Przewalskia tangutica
Physochlaina orientalis
Hyoscyamus niger
Anisodus tanguticus
Nicandra physalodes
Schultesianthus leucanthus
Solandra grandiflora
Dyssochroma viridiflorum
Markea coccinea
Merinthopodium neuranthum
Hawkesiophyton panamense
Juanulloa mexicana
Exodeconus miersii
Trompettia cardenasiana
Datura ferox
Brugmansia suaveolens
Jaltomata grandiflora
Jaltomata bicolor
Solanum thelopodium
Solanum evolvuloides
Solanum crinitum
Solanum sisymbriifolium
Solanum hieronymi
Solanum barbisetum
Solanum oedipus
Solanum mahoriense
Solanum sejunctum
Solanum jamaicense
Solanum torvum
Solanum polygamum
Solanum montanum
Solanum pinnatum
Solanum evolvulifolium
Solanum trizygum
Solanum carpense
Solanum palustre
Solanum microdontum
Solanum peruvianum
Solanum pimpinellifolium
Solanum uncinellum
Solanum crispum
Solanum corymbosum
Solanum weddellii
Solanum sinuatirecurvum
Solanum salamancae
Solanum annuum
Solanum valdiviense
Solanum terminale
Solanum laciniatum
Solanum herculeum
Capsicum chacoense
Lycianthes rantonnetii
Lycianthes acapulcensis
Nectouxia formosa
Salpichroa tristis
Cuatresia harlingiana
Darcyanthus spruceanus
Deprea abra patriciae
Deprea pauciflora
Athenaea brasiliana
Athenaea cuspidata
Withania somnifera
Mellissia begoniifolia
Discopodium penninervium
Nothocestrum latifolium
Archiphysalis chamaesarachoides
Tubocapsicum anomalum
Physaliastrum heterophyllum
Oryctes nevadensis
Schraderanthus viscosus
Tzeltalia amphitricha
Witheringia nelsonii
Witheringia solanacea
Capsicophysalis potosina
Alkekengi officinarum
Calliphysalis carpenteri
Quincula lobata
Chamaesaracha arida
Physalis peruviana
Physalis angulata
Leucophysalis grandiflora
Trozelia umbellata
Dunalia brachyacantha
Saracha quitensis
Iochroma cyaneum
Vassobia breviflora
Eriolarynx australis
Mandragora officinarum
Nicotiana wigandioides
Symonanthus aromaticus
Anthocercis intricata
Anthotroche pannosa
Grammosolen dixonii
Duboisia myoporoides
Cyphanthera anthocercidea
Crenidium spinescens
Combera paradoxa
Pantacantha ameghinoi
Benthamiella patagonica
Vestia foetida
Cestrum bracteatum
Sessea stipulata
Protoschwenkia mandonii
Browallia americana
Streptosolen jamesonii
Salpiglossis sinuata
Reyesia chilensis
Schizanthus grahamii
Duckeodendron cestroides

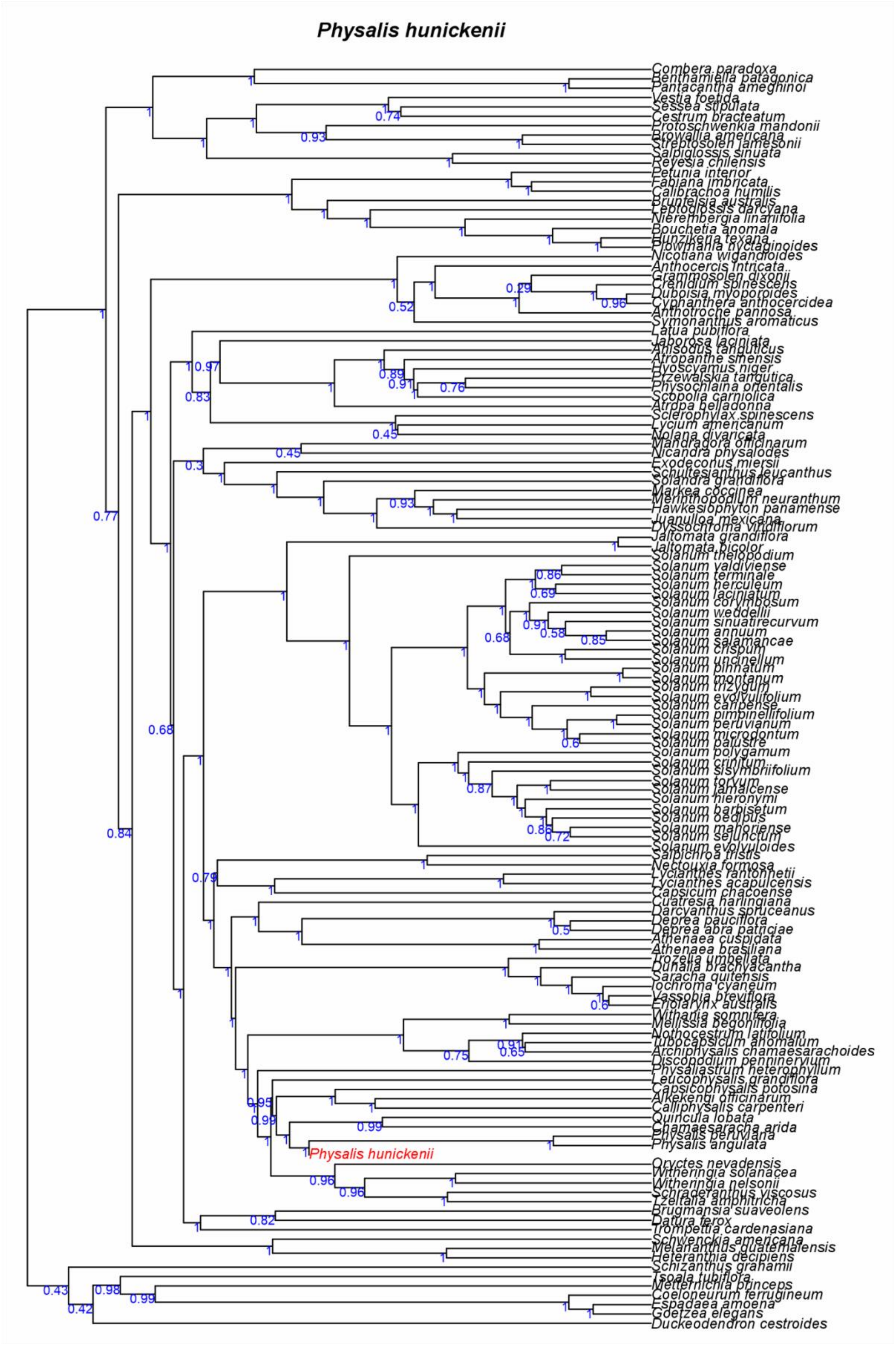
Physalis hunickenii
Combera paradoxa
Benthamiella patagonica
Pantacantha ameghinoi
Vestia foetida
Sessea stipulata
Cestrum bracteatum
Protoschwenkia mandonii
Browallia americana
Streptosolen jamesonii
Salpiglossis sinuata
Reyesia chilensis
Petunia interior
Fabiana imbricata
Calibrachoa humilis
Brunfelsia australis
Leptoglossis darcyana
Nierembergia linariifolia
Bouchetia anomala
Hunzikeria texana
Plowmania nyctaginoides
Nicotiana wigandioides
Anthocercis intricata
Grammosolen dixonii
Crenidium spinescens
Duboisia myoporoides
Cyphanthera anthocercidea
Anthotroche pannosa
Symonanthus aromaticus
Latua pubiflora
Jaborosa laciniata
Anisodus tanguticus
Atropanthe sinensis
Hyoscyamus niger
Przewalskia tangutica
Physochlaina orientalis
Scopolia carniolica
Atropa belladonna
Sclerophylax spinescens
Lycium americanum
Nolana divaricata
Mandragora officinarum
Nicandra physalodes
Exodeconus miersii
Schultesianthus leucanthus
Solandra grandiflora
Markea coccinea
Merinthopodium neuranthum
Hawkesiophyton panamense
Juanulloa mexicana
Dyssochroma viridiflorum
Jaltomata grandiflora
Jaltomata bicolor
Solanum thelopodium
Solanum valdiviense
Solanum terminale
Solanum herculeum
Solanum laciniatum
Solanum corymbosum
Solanum weddellii
Solanum sinuatirecurvum
Solanum annuum
Solanum salamancae
Solanum crispum
Solanum uncinellum
Solanum pinnatum
Solanum montanum
Solanum trizygum
Solanum evolvulifolium
Solanum caripense
Solanum pimpinellifolium
Solanum peruvianum
Solanum microdontum
Solanum palustre
Solanum polygamum
Solanum crinitum
Solanum sisymbriifolium
Solanum torvum
Solanum jamaicense
Solanum hieronymi
Solanum barbisetum
Solanum oedipus
Solanum mahoriense
Solanum sejunctum
Solanum evolvuloides
Salpichroa tristis
Nectouxia formosa
Lycianthes rantonnetii
Lycianthes acapulcensis
Capsicum chacoense
Cuatresia harlingiana
Darcyanthus spruceanus
Deprea pauciflora
Deprea abra patriciae
Athenaea cuspidata
Athenaea brasiliana
Trozelia umbellata
Dunalia brachyacantha
Saracha quitensis
Iochroma cyaneum
Vassobia breviflora
Eriolarynx australis
Withania somnifera
Mellissia begoniifolia
Nothocestrum latifolium
Tubocapsicum anomalum
Archiphysalis chamaesarachoides
Discopodium penninervium
Physaliastrum heterophyllum
Leucophysalis grandiflora
Capsicophysalis potosina
Alkekengi officinarum
Calliphysalis carpenteri
Quincula lobata
Chamaesaracha arida
Physalis peruviana
Physalis angulata
Physalis hunickenii
Oryctes nevadensis
Witheringia solanacea
Witheringia nelsonii
Schraderanthus viscosus
Tzeltalia amphitricha
Brugmansia suaveolens
Datura ferox
Trompettia cardenasiana
Schwenckia americana
Melananthus guatemalensis
Heteranthia decipiens
Schizanthus grahamii
Tsoala tubiflora
Metternichia princeps
Coeloneurum ferrugineum
Espadaea amoena
Goetzea elegans
Duckeodendron cestroides

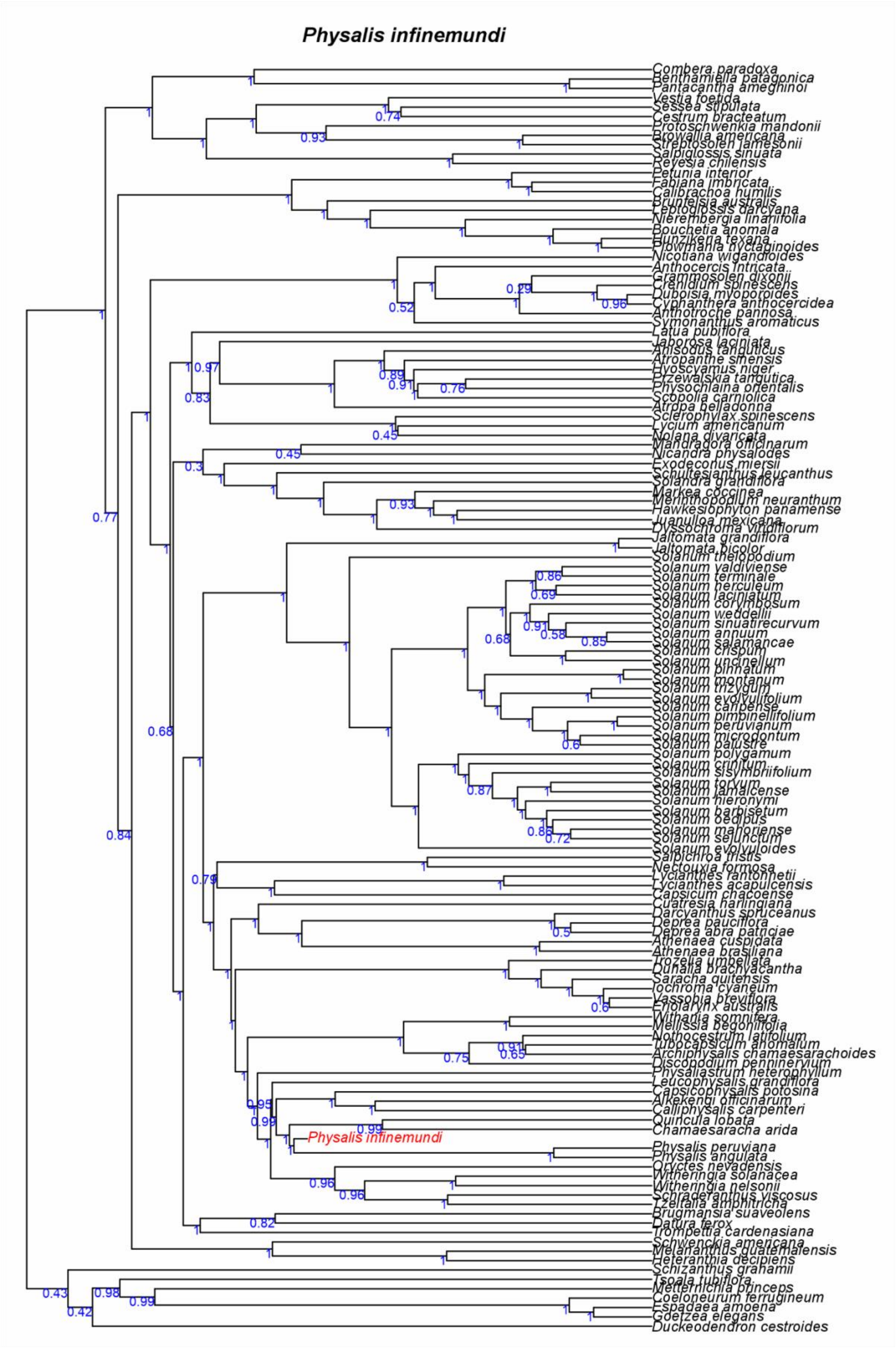
Physalis infinemundi
Combera paradoxa
Benthamiella patagonica
Pantacantha ameghinoi
Vestia foetida
Sessea stipulata
Cestrum bracteatum
Protoschwenkia mandonii
Browallia americana
Streptosolen jamesonii
Salpiglossis sinuata
Reyesia chilensis
Petunia interior
Fabiana imbricata
Calibrachoa humilis
Brunfelsia australis
Leptoglossis darcyana
Nierembergia linariifolia
Bouchetia anomala
Hunzikeria texana
Plowmania nyctaginoides
Nicotiana wigandioides
Anthocercis intricata
Grammosolen dixonii
Crenidium spinescens
Duboisia myoporoides
Cyphanthera anthocercidea
Anthotroche pannosa
Symonanthus aromaticus
Latua pubiflora
Jaborosa laciniata
Anisodus tanguticus
Atropanthe sinensis
Hyoscyamus niger
Przewalskia tangutica
Physochlaina orientalis
Scopolia carniolica
Atropa belladonna
Sclerophylax spinescens
Lycium americanum
Nolana divaricata
Mandragora officinarum
Nicandra physalodes
Exodeconus miersii
Schultesianthus leucanthus
Solandra grandiflora
Markea coccinea
Merinthopodium neuranthum
Hawkesiophyton panamense
Juanulloa mexicana
Dyssochroma viridiflorum
Jaltomata grandiflora
Jaltomata bicolor
Solanum thelopodium
Solanum valdiviense
Solanum terminale
Solanum herculeum
Solanum laciniatum
Solanum corymbosum
Solanum weddellii
Solanum sinuatirecurvum
Solanum annuum
Solanum salamancae
Solanum crispum
Solanum uncinellum
Solanum pinnatum
Solanum montanum
Solanum trizygum
Solanum evolvulifolium
Solanum caripense
Solanum pimpinellifolium
Solanum peruvianum
Solanum microdontum
Solanum palustre
Solanum polygamum
Solanum crinitum
Solanum sisymbriifolium
Solanum torvum
Solanum jamaicense
Solanum hieronymi
Solanum barbisetum
Solanum oedipus
Solanum mahoriense
Solanum sejunctum
Solanum evolvuloides
Salpichroa tristis
Nectouxia formosa
Lycianthes rantonnetii
Lycianthes acapulcensis
Capsicum chacoense
Cuatresia harlingiana
Darcyanthus spruceanus
Deprea pauciflora
Deprea abra patriciae
Athenaea cuspidata
Athenaea brasiliana
Trozelia umbellata
Dunalia brachyacantha
Saracha quitensis
Iochroma cyaneum
Vassobia breviflora
Eriolarynx australis
Withania somnifera
Mellissia begoniifolia
Nothocestrum latifolium
Tubocapsicum anomalum
Archiphysalis chamaesarachoides
Discopodium penninervium
Physaliastrum heterophyllum
Leucophysalis grandiflora
Capsicophysalis potosina
Alkekengi officinarum
Calliphysalis carpenteri
Quincula lobata
Chamaesaracha arida
Physalis infinemundi
Physalis peruviana
Physalis angulata
Oryctes nevadensis
Witheringia solanacea
Witheringia nelsonii
Schraderanthus viscosus
Tzeltalia amphitricha
Brugmansia suaveolens
Datura ferox
Trompettia cardenasiana
Schwenckia americana
Melananthus guatemalensis
Heteranthia decipiens
Schizanthus grahamii
Tsoala tubiflora
Metternichia princeps
Coeloneurum ferrugineum
Espadaea amoena
Goetzea elegans
Duckeodendron cestroides

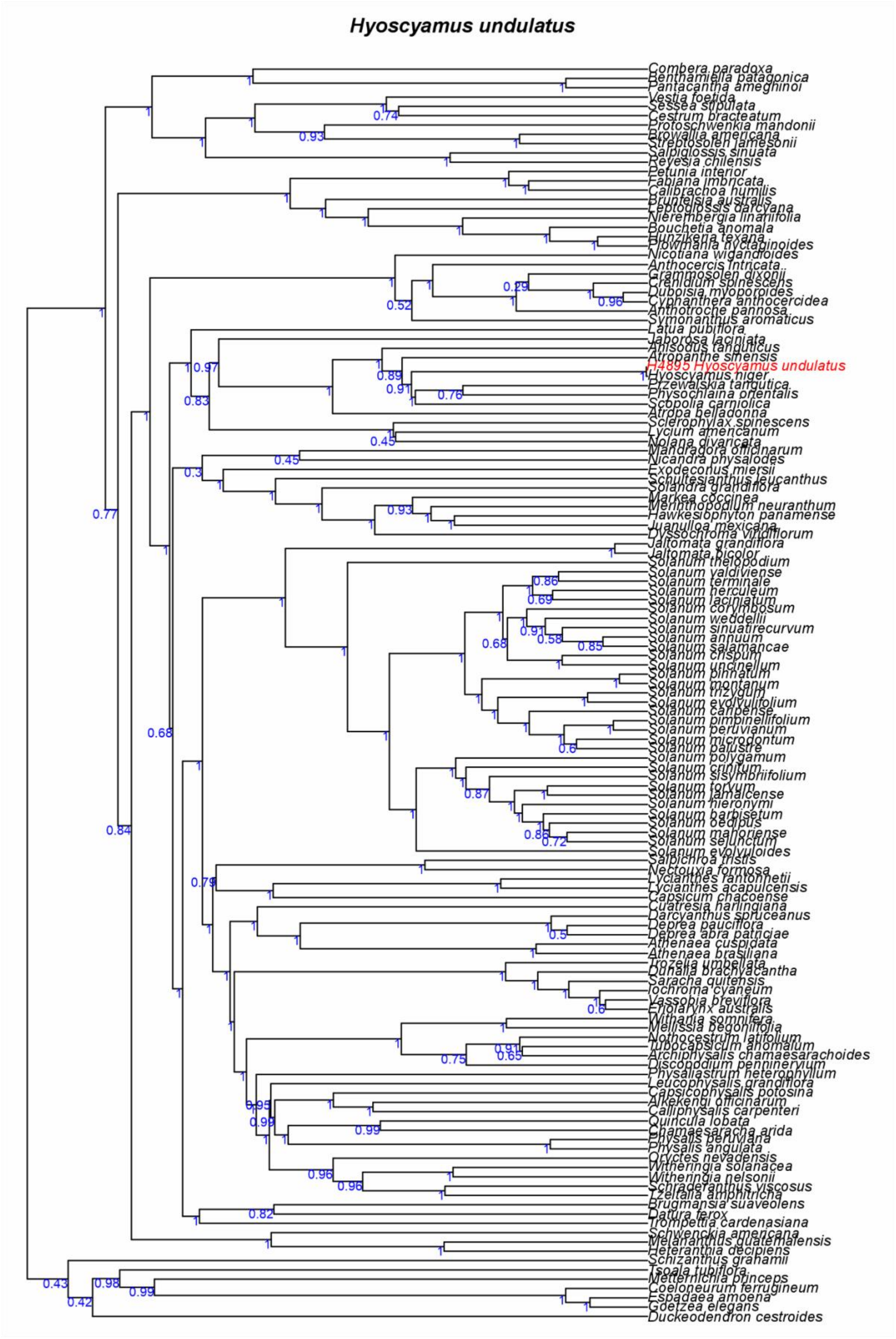
Hyoscyamus undulatus
Combera paradoxa
Benthamiella patagonica
Pantacantha ameghinoi
Vestia foetida
Sessea stipulata
Cestrum bracteatum
Protoschwenkia mandonii
Browallia americana
Streptosolen jamesonii
Salpiglossis sinuata
Reyesia chilensis
Petunia interior
Fabiana imbricata
Calibrachoa humilis
Brunfelsia australis
Leptoglossis darcyana
Nierembergia linariifolia
Bouchetia anomala
Hunzikeria texana
Plowmania nyctaginoides
Nicotiana wigandioides
Anthocercis intricata
Grammosolen dixonii
Crenidium spinescens
Duboisia myoporoides
Cyphanthera anthocercidea
Anthotroche pannosa
Symonanthus aromaticus
Latua pubiflora
Jaborosa laciniata
Anisodus tanguticus
Atropanthe sinensis
H4895 Hyoscyamus undulatus
Hyoscyamus niger
Przewalskia tangutica
Physochlaina orientalis
Scopolia carniolica
Atropa belladonna
Sclerophylax spinescens
Lycium americanum
Nolana divaricata
Mandragora officinarum
Nicandra physalodes
Exodeconus miersii
Schultesianthus leucanthus
Solandra grandiflora
Markea coccinea
Merinthopodium neuranthum
Hawkesiophyton panamense
Juanulloa mexicana
Dyssochroma viridiflorum
Jaltomata grandiflora
Jaltomata bicolor
Solanum thelopodium
Solanum valdiviense
Solanum terminale
Solanum herculeum
Solanum laciniatum
Solanum corymbosum
Solanum weddellii
Solanum sinuatirecurvum
Solanum annuum
Solanum salamancae
Solanum crispum
Solanum uncinellum
Solanum pinnatum
Solanum montanum
Solanum trizygum
Solanum evolvulifolium
Solanum caripense
Solanum pimpinellifolium
Solanum peruvianum
Solanum microdontum
Solanum palustre
Solanum polygamum
Solanum crinitum
Solanum sisymbriifolium
Solanum torvum
Solanum jamaicense
Solanum hieronymi
Solanum barbisetum
Solanum oedipus
Solanum mahoriense
Solanum sessilestellatum
Solanum evolvuloides
Salpichroa tristis
Nectouxia formosa
Lycianthes rantonnetii
Lycianthes acapulcensis
Capsicum chacoense
Cuatresia harlingiana
Darcyanthus spruceanus
Deprea pauciflora
Deprea abra patriciae
Athenaea cuspidata
Athenaea brasiliana
Trozelia umbellata
Dunalia brachyacantha
Saracha quitensis
Iochroma cyaneum
Vassobia breviflora
Eriolarynx australis
Withania somnifera
Melissia begoniifolia
Nothocestrum latifolium
Tubocapsicum anomalum
Archiphysalis chamaesarachoides
Discopodium penninervium
Physaliastrum heterophyllum
Leucophysalis grandiflora
Capsicophysalis potosina
Alkekengi officinarum
Calliphysalis carpenteri
Quincula lobata
Chamaesaracha arida
Physalis peruviana
Physalis angulata
Oryctes nevadensis
Witheringia solanacea
Witheringia nelsonii
Schraderanthus viscosus
Tzeltalia amphitricha
Brugmansia suaveolens
Datura ferox
Trompettia cardenasiana
Schwenckia americana
Melananthus guatemalensis
Heteranthia decipiens
Schizanthus grahamii
Tsoala tubiflora
Metternichia princeps
Coeloneurum ferrugineum
Espadaea amoena
Goetzea elegans
Duckeodendron cestroides

**Figure S11.** Mean age and 95%HPD estimates for extant Solanaceae based on the time-heterogeneous total evidence dating analysis. The nodes in the table correspond to the nodes in the phylogeny.

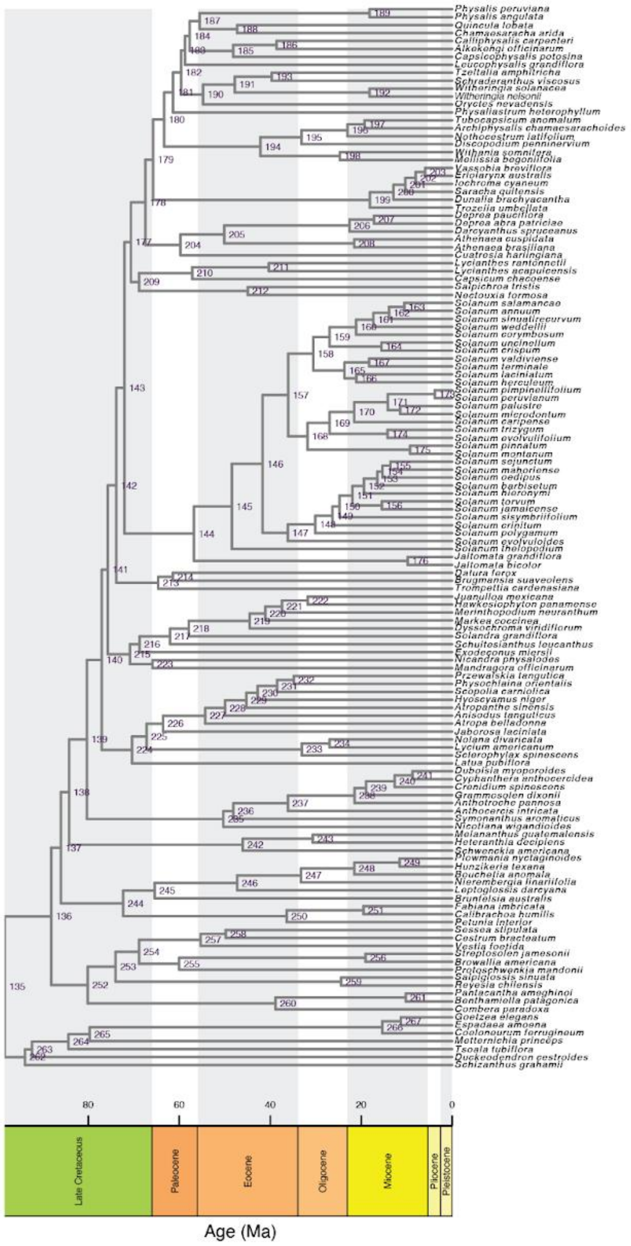


| node | mean age | minHPD | maxHPD | node | mean age | minHPD | maxHPD |
|---|---|---|---|---|---|---|---|
| 135 | 98.38 | 95.66 | 100.27 | 202 | 8.00 | 4.51 | 12.18 |
| 136 | 88.14 | 80.45 | 95.56 | 203 | 5.93 | 3.05 | 9.85 |
| 137 | 86.02 | 78.87 | 92.93 | 204 | 59.73 | 48.92 | 69.06 |
| 138 | 84.23 | 77.02 | 90.86 | 205 | 50.08 | 32.24 | 63.66 |
| 139 | 80.30 | 73.75 | 86.57 | 206 | 22.56 | 9.28 | 38.87 |
| 140 | 77.02 | 70.85 | 82.88 | 207 | 17.19 | 6.33 | 28.23 |
| 141 | 75.70 | 70.10 | 81.58 | 208 | 21.54 | 11.97 | 34.18 |
| 142 | 73.85 | 68.63 | 79.55 | 209 | 68.83 | 62.65 | 74.59 |
| 143 | 72.04 | 67.09 | 77.66 | 210 | 57.14 | 48.51 | 67.53 |
| 144 | 56.76 | 45.09 | 69.09 | 211 | 40.31 | 20.40 | 55.22 |
| 145 | 48.43 | 37.90 | 59.38 | 212 | 44.94 | 27.60 | 62.46 |
| 146 | 41.65 | 31.55 | 51.34 | 213 | 64.61 | 55.47 | 73.36 |
| 147 | 36.13 | 26.87 | 45.05 | 214 | 61.39 | 51.94 | 70.77 |
| 148 | 30.08 | 21.93 | 37.78 | 215 | 70.87 | 63.14 | 78.29 |
| 149 | 26.30 | 19.98 | 33.65 | 216 | 68.71 | 58.00 | 78.74 |
| 150 | 24.75 | 18.95 | 30.93 | 217 | 62.02 | 52.19 | 72.98 |
| 151 | 21.92 | 16.92 | 27.51 | 218 | 57.93 | 47.35 | 70.03 |
| 152 | 19.39 | 14.48 | 24.41 | 219 | 44.52 | 31.88 | 57.93 |
| 153 | 16.42 | 12.03 | 21.30 | 220 | 41.07 | 28.15 | 55.12 |
| 154 | 15.30 | 11.16 | 20.09 | 221 | 37.40 | 22.63 | 50.58 |
| 155 | 13.59 | 8.96 | 18.77 | 222 | 31.73 | 19.29 | 44.13 |
| 156 | 15.44 | 8.43 | 23.03 | 223 | 65.88 | 40.64 | 79.59 |
| 157 | 36.03 | 27.03 | 45.65 | 224 | 70.38 | 61.18 | 78.57 |
| 158 | 30.55 | 21.44 | 39.16 | 225 | 67.18 | 57.73 | 76.30 |
| 159 | 26.94 | 18.94 | 35.28 | 226 | 63.58 | 53.91 | 73.11 |
| 160 | 21.11 | 14.55 | 28.19 | 227 | 54.28 | 42.42 | 65.21 |
| 161 | 17.29 | 10.98 | 23.03 | 228 | 49.92 | 37.52 | 60.78 |
| 162 | 13.87 | 7.91 | 19.57 | 229 | 45.25 | 32.05 | 57.88 |
| 163 | 10.52 | 5.11 | 16.23 | 230 | 42.75 | 30.09 | 56.92 |
| 164 | 15.54 | 5.87 | 24.40 | 231 | 38.50 | 25.54 | 51.79 |
| 165 | 23.64 | 16.77 | 33.30 | 232 | 34.82 | 21.46 | 47.89 |
| 166 | 21.05 | 13.20 | 29.86 | 233 | 33.04 | 17.82 | 59.32 |
| 167 | 18.24 | 7.13 | 26.94 | 234 | 26.94 | 10.13 | 53.76 |
| 168 | 31.78 | 22.43 | 41.37 | 235 | 50.29 | 31.13 | 70.60 |
| 169 | 26.93 | 18.81 | 36.15 | 236 | 48.11 | 28.94 | 67.87 |
| 170 | 21.53 | 14.14 | 29.38 | 237 | 36.10 | 20.57 | 53.48 |
| 171 | 14.15 | 8.36 | 20.59 | 238 | 21.44 | 13.08 | 30.11 |
| 172 | 11.40 | 4.59 | 17.84 | 239 | 18.91 | 11.01 | 26.17 |
| 173 | 3.80 | 1.12 | 7.21 | 240 | 12.62 | 6.74 | 18.96 |
| 174 | 14.21 | 5.57 | 23.05 | 241 | 8.69 | 3.01 | 14.83 |
| 175 | 9.25 | 2.08 | 18.68 | 242 | 46.02 | 25.54 | 66.40 |
| 176 | 9.75 | 2.24 | 19.92 | 243 | 30.63 | 14.57 | 48.72 |
| 177 | 70.61 | 65.25 | 75.71 | 244 | 72.32 | 59.86 | 84.56 |
| 178 | 67.46 | 62.70 | 72.59 | 245 | 65.40 | 50.72 | 79.19 |
| 179 | 65.74 | 61.05 | 70.50 | 246 | 47.27 | 32.24 | 63.45 |
| 180 | 63.31 | 59.17 | 67.74 | 247 | 33.24 | 19.11 | 47.86 |
| 181 | 61.36 | 57.65 | 65.56 | 248 | 21.52 | 11.67 | 33.31 |
| 182 | 59.60 | 56.06 | 63.19 | 249 | 11.54 | 5.06 | 18.81 |
| 183 | 58.73 | 55.55 | 62.43 | 250 | 36.37 | 17.63 | 58.69 |
| 184 | 57.70 | 54.75 | 61.31 | 251 | 19.50 | 9.10 | 30.93 |
| 185 | 48.14 | 30.39 | 59.25 | 252 | 80.12 | 69.14 | 90.38 |
| 186 | 38.59 | 21.24 | 53.04 | 253 | 74.00 | 62.70 | 84.66 |
| 187 | 55.51 | 52.95 | 58.52 | 254 | 68.85 | 57.92 | 79.64 |
| 188 | 47.28 | 37.05 | 56.03 | 255 | 60.02 | 42.55 | 75.19 |
| 189 | 18.15 | 6.95 | 32.07 | 256 | 19.03 | 7.48 | 31.71 |
| 190 | 54.77 | 43.21 | 63.63 | 257 | 55.29 | 43.10 | 68.16 |
| 191 | 47.78 | 31.27 | 59.69 | 258 | 49.77 | 36.45 | 61.98 |
| 192 | 18.16 | 6.40 | 30.81 | 259 | 24.40 | 7.94 | 45.61 |
| 193 | 39.65 | 22.28 | 54.57 | 260 | 38.82 | 18.37 | 59.27 |
| 194 | 42.17 | 26.44 | 60.23 | 261 | 10.20 | 3.85 | 18.19 |
| 195 | 33.11 | 19.54 | 49.89 | 262 | 93.94 | 87.38 | 99.68 |
| 196 | 22.99 | 12.52 | 34.84 | 263 | 92.37 | 83.96 | 99.44 |
| 197 | 19.26 | 8.37 | 30.19 | 264 | 84.36 | 72.96 | 94.97 |
| 198 | 24.67 | 9.65 | 49.61 | 265 | 79.69 | 69.16 | 91.16 |
| 199 | 18.06 | 10.63 | 25.84 | 266 | 15.35 | 7.84 | 22.98 |
| 200 | 12.89 | 7.61 | 18.82 | 267 | 11.25 | 5.11 | 18.28 |
| 201 | 10.27 | 5.99 | 15.14 | | | | |

### SUPPLEMENTARY TABLES

**Table S1.** Previously reported divergence time estimates for Solanaceae.

| Publication | Solanaceae | | Solanales | | Angiosperms | |
|---|---|---|---|---|---|---|
| | Crown age (Ma) | Stem age (Ma) | Crown age (Ma) | Stem age (Ma) | Crown age (Ma) | Stem age (Ma) |
| Qin et al., 2014 | 156 | | | | | |
| Magallon et al., 2015 | -- | 66.65 (47.8-86.9) | 79.24 (68-96.7) | 85.91 (75.4-100.3) | 139.4 | 325.1 |
| Montoya et al., 2018 | 40* | 75* | 66-105 | 110* | 149-256 | -- |
| Li et al., 2019 | -- | 82.5 (58.7-102.9) | 86.3 (107.2-61.5) | 87.4 (108.5-63.4) | 209 (267-187) | -- |
| Ramirez Barahona et al., 2020 | 58.6 (52.2-67.6) | 86.8 (68.9-102.2) | 101.6 (90.1-115.1) | 110.9 (100.4-119.4) | | |
| Silvestro et al., 2021 | | 71.4 (55–100.2)** | -- | -- | 254.8-153.7 | -- |
| Särkinen et al., 2013 | 30 (26-34) | 49 (46-54) | -- | -- | -- | -- |
| Huang et al., 2023 | 73.3 (72-74.3) | 93.9 (95.8-103.2) | -- | -- | -- | -- |
| Zuntini et al., 2024 (young tree) | 81.99 | 114.96 | 130.86 | 137.47 | -- | -- |
| Zuntini et al., 2024 (old tree) | 109.99 | 174.33 | 204.62 | 217.89 | -- | -- |

* Inferred from the figure, not reported; ** used 'time of origin'

**Table S2.** Solanaceae seeds and fruit fossils included in this study. The geological time scale from the International Chronostratigraphic Chart v. 2024/12 was used for stratigraphic ranges (Dec 2024; updated).

| Species | Reference | Stratigraphic range as Epoch: Stage (age) | Specimen used | Occurrence of the specimen | Repository of the specimen |
|---|---|---|---|---|---|
| *Physalis infinemundi* Wilf. | Wilf et al. (2017) | 52.44–52 Ma | MPEF-Pb 6434a,b | Laguna del Hunco, Huitrera Formation, Chubut Province, Argentina. | Museo Paleontológico Egidio Feruglio, Trelew, Argentina (MPEF-Pb). |
| *Physalis hunickenii* Deanna, Wilf & Gandolfo | Deanna et al. (2020) | 52.44–52 Ma | MPEF-Pb 6443a,b | Laguna del Hunco, Huitrera Formation, Chubut Province, Argentina. | Museo Paleontológico Egidio Feruglio, Trelew, Argentina (MPEF-Pb) |
| *Eophysaloides inflata* Martínez-A. & Deanna | Deanna et al. (2023) | 47.3–33.9 Ma | STRI-SGC 36163 | Eocene Esmeraldas Formation, Colombia. | Museo Paleontologico Jose Royo y Gomez, Colombian Geological Survey (SGC), Bogota, Colombia. |
| *Lycianthoides calycina* Deanna & Manchester | Deanna et al. (2023) | 49.5–51.5 Ma | UCM 41276a,b | Claudia's Place (UCM locality 20099063), Green River Formation, Garfield County, CO, USA. | University of Colorado Museum (UCM), Paleontological Section, Boulder, CO, USA. |
| *Hyoscyosperma daturoides* Deanna & S.D.Sm. | Deanna et al. (2025) | 27.8 – 0.01 Ma | K453/133 | Rostov oblast, Northern Caucasus, Russia. | Komarov Botanical Institute, St. Petersburg, Russia. |

| | | | | | |
|---|---|---|---|---|---|
| *Solanum foveolatum* Negru | Negru (1986) | 33.9 – 0.01 Ma | MB.Pb.1998/0 434 | Bauersberg, Germany. | Museum für Naturkunde Berlin, Germany. |
| *Solanum miocenicum* Deanna & S.D.Sm. | Deanna et al. (2025) | 27.82 – 0.01 Ma | K528/50 | Omsk oblast, West Siberia, Russia. | Komarov Botanical Institute, St. Petersburg, Russia. |
| *Solanispermum reniforme* M.Chandler | Chandler (1957) | 48-0.01 Ma | V-40891 | Arne, Poole Formation Sandbanks, Poole Formation, United Kingdom. | Natural History Museum, London, United Kingdom. |
| *Hyoscyamus undulatus* Deanna & S.D.Sm. | Deanna et al. (2025) | 23.03-0.01 Ma | H4895/49 | Novosibirsk oblast, West Siberia, Russia. | Komarov Botanical Institute, St. Petersburg, Russia. |
| *Solanoides dorofeevii* Deanna & S.D.Sm. | Deanna et al. (2025) | 33.9 – 0.77 Ma | K530/21 | Krasnodar kray, Northern Caucasus, Russia. | Komarov Botanical Institute, St. Petersburg, Russia. |
| *Albionites arnensis* (M.Chandler) Deanna & S.Knapp | Chandler (1962); Deanna et al. (2025) | 48-46 Ma | V40898 | Arne, Poole Formation, United Kingdom. | Natural History Museum, London, United Kingdom. |
| *Nephrosemen reticulatum* Manchester | Manchester (1994) | 47.8-0.01 Ma | UF6500 | Nut Beds flora, Clarno Formation, north-central Oregon, United States. | Florida Museum of Natural History, University of Florida, United States. |
| *Capsicum pliocenicum* | Deanna et al. (2025) | 5.33-0.13 Ma | MRSN-P/345-CCN6065b | Bucine, Italy. | CENOFITA collection, managed by the Regional |

| | | | | | |
|---|---|---|---|---|---|
| Deanna & S.D.Sm. | | | | | Museum of Natural Sciences of Turin, Turin, Italy. |
| *Thanatosperma minutum* Deanna & S.Knapp | Deanna et al. (2025) | 5.33-0.001 Ma | MB.Pb.2003/0 073 | Oberzella a. d. Werra, Germany. | Museum für Naturkunde, Berlin, Germany. |

**Table S3.** Coding scheme for seed and fruit characters used in this study. **Bold** characters are secondary traits, calculated from primary measurements.

| Character | # | Continuous (C) / Discrete (D) | Character states |
|---|---|---|---|
| Seed length | 1 | C | -- |
| **Seed length/width ratio** | 2 | C | -- |
| Seed compression | 3 | D | 0= flattened<br>1= not flattened |
| Hilum position in the seed | 4 | D | 0= lateral or sub-lateral<br>1= terminal |
| Embryo shape | 5 | D | 0 = curved<br>1= straight<br>2 = circinate/coiled |
| Seed hilar-chalazal cavity | 6 | D | 0= absent<br>1= present |
| Exotestal cell walls | 7 | D | 0= straight<br>1= sinuate-cerebelloid |
| Seed wings | 8 | D | 0= absent<br>1= present |
| Seed elaiosomes | 9 | D | 0= absent<br>1= present |
| Fruiting calyx length | 10 | C | -- |
| **Fruiting calyx length/width ratio** | 11 | C | -- |
| Length of the longest fruiting calyx lobe | 12 | C | -- |
| **Longest fruiting calyx lobe length/ fruiting calyx length ratio** | 13 | C | -- |
| Fruiting pedicel length | 14 | C | -- |

| | | | |
|---|---|---|---|
| **Fruit length/width ratio** | 15 | C | -- |
| Fruiting calyx base invagination | 16 | D | 0= absent<br>1= present |
| Fruiting calyx angled | 17 | D | 0= absent<br>1= present |
| Fruiting calyx lobes sinus | 18 | D | 0= rounded<br>1= angular (if angular because it's broken when ripe, score it as angular)<br>2= absent |
| Fruiting calyx teeth | 19 | D | 0= absent<br>1= present |
| Fruiting calyx widest veins distinct and terminating in lobe tips | 20 | D | 0= absent<br>1= present |
| Fruiting calyx secondary veins distinct from other vein orders and emerge from the base | 21 | D | 0= absent<br>1= present |
| Fruiting calyx secondary veins fork before the lobe sinus | 22 | D | 0= absent<br>1= present |
| Fruit type | 23 | D | 0= berry<br>1= capsule<br>2= diclesium or mericarp<br>3= drupe<br>4= non-capsular dehiscent fruits<br>5= pyxidium |
| Fruiting calyx inflated | 24 | D | 0= absent<br>1= present (the calyx tube length exceeds by at least 20% the fruit diameter) |
| Fruiting calyx venation pattern | 25 | D | 0= parallelodromous<br>1= other type |

**Table S4.** Phylogenetic signal in binary morphological variables. Mean, minimum, and maximum values across 100 randomly selected trees from the posterior distribution are given for Blomberg's K statistic.

| Trait | MinK | MaxK | MeanK | MinP | MaxP | MeanP | Conclusion |
|---|---|---|---|---|---|---|---|
| seed length | 1.530 | 5.020 | 3.096 | 0.001 | 0.001 | 0.001 | Significant Phylogenetic Signal |
| seed ratio | 0.880 | 2.497 | 1.270 | 0.001 | 0.015 | 0.004 | Significant Phylogenetic Signal |
| fruiting calyx length | 0.568 | 0.749 | 0.671 | 0.001 | 0.008 | 0.003 | Significant Phylogenetic Signal |
| fruiting calyx ratio | 0.919 | 1.545 | 1.146 | 0.001 | 0.002 | 0.001 | Significant Phylogenetic Signal |
| calyx lobe length | 0.536 | 0.751 | 0.672 | 0.001 | 0.006 | 0.002 | Significant Phylogenetic Signal |
| calyx lobe ratio | 0.427 | 0.583 | 0.515 | 0.008 | 0.097 | 0.036 | Significant Phylogenetic Signal |
| fruiting pedicel | 0.535 | 0.755 | 0.666 | 0.001 | 0.007 | 0.002 | Significant Phylogenetic Signal |
| fruit ratio | 0.956 | 1.968 | 1.282 | 0.001 | 0.002 | 0.001 | Significant Phylogenetic Signal |

**Table S5.** Phylogenetic signal in binary morphological variables. Mean, minimum, and maximum values across the 100 sampled trees are given for the D-statistic. Values of D smaller than 0 are more conserved than expected under Brownian motion (light orange), and values near zero are consistent with Brownian motion (light yellow). Values of D between zero and 1 (light green) suggest a weak signal, and those around 1 or greater (light blue) are phylogenetically over-dispersed.

| | Estimated D | | | Prob. of E(D) resulting from no (random) phy. structure | | | Prob. of E(D) resulting from Brownian phy. structure | | | Interpretation |
|---|---|---|---|---|---|---|---|---|---|---|
| | MinD | MaxD | MeanD | MinPval1 | MaxPval1 | MeanPval1 | MinPval0 | MaxPval0 | MeanPval0 | |
| calyx_teeth | -5.160 | -3.033 | -4.044 | 0 | 0.001 | 1.00E-05 | 0.967 | 0.998 | 0.982 | Strong signal, consistent across trees |
| venation_pattern | -3.430 | -2.192 | -2.849 | 0 | 0.001 | 2.00E-05 | 0.954 | 0.996 | 0.977 | Strong signal, consistent across trees |
| compression | -1.565 | -1.086 | -1.333 | 0 | 0 | 0 | 0.999 | 1.000 | 1.000 | Strong signal, consistent across trees |
| hilum_position | -0.672 | -0.244 | -0.445 | 0 | 0 | 0 | 0.745 | 0.923 | 0.852 | Strong signal, consistent across trees |
| secondary_veins_fork | -0.443 | -0.133 | -0.289 | 0 | 0.000 | 0.00 | 0.666 | 0.877 | 0.786 | Strong signal, consistent across trees |
| secondary_veins | -0.077 | 0.186 | 0.050 | 0 | 0.001 | 0.00 | 0.271 | 0.617 | 0.452 | Moderate signal, consistent across trees |
| calyx_inflated | -0.043 | 0.335 | 0.121 | 0 | 0.004 | 0.0005 | 0.208 | 0.574 | 0.404 | Moderate signal, consistent across trees |
| cavity | 0.027 | 0.302 | 0.168 | 0 | 0.001 | 2.00E-05 | 0.126 | 0.480 | 0.300 | Moderate signal, consistent across trees |
| widest_veins | 0.137 | 0.485 | 0.323 | 0 | 0.014 | 0.00 | 0.080 | 0.381 | 0.202 | Moderate signal, varies across trees |
| testal_walls | 0.423 | 0.604 | 0.523 | 0 | 0.015 | 0.01 | 0.011 | 0.079 | 0.032 | Weak signal, consistent across trees |
| wings | -0.060 | 1.283 | 0.687 | 0.077 | 0.630 | 0.34 | 0.159 | 0.607 | 0.348 | Weak signal, consistent across trees |
| base_angled | 0.539 | 0.951 | 0.740 | 0.065 | 0.407 | 0.18 | 0.015 | 0.172 | 0.071 | Weak signal, varies across trees |
| base_invaginated | 0.581 | 1.083 | 0.861 | 0.069 | 0.584 | 0.31 | 0.011 | 0.132 | 0.046 | Weak signal, varies across trees |
| elaiosomes | -9.281 | 10.844 | 2.069 | 0.035 | 0.744 | 0.55 | 0.198 | 0.935 | 0.394 | No signal, consistent across trees |
| embryo | NA | NA | NA | NA | NA | NA | NA | NA | NA | NA |
| lobe_sinus | NA | NA | NA | NA | NA | NA | NA | NA | NA | NA |
| fruit_type | NA | NA | NA | NA | NA | NA | NA | NA | NA | NA |

**Table S6.** Posterior probability of sister groups for the fossils whose position was unconstrained. These posterior probabilities were calculated by analyzing the "one at a time" fossil traces. For the fossils that were retrieved in more than 10 possible clades, we report the 10 groups with the highest PP, and we show the cumulative posterior of these 10 most probable groupings.

| Fossil | Sister group clades | PP | Included in final summary tree (Fig. 2) |
|---|---|---|---|
| ***Lycianthoides calycina*** (5 different sampled sister groups) | c("Lycianthoides_calycina", "Lycianthes_acapulcensis", "Lycianthes_rantonnetii") | 0.55446043 | yes |
| | c("Capsicum_chacoense", "Lycianthoides_calycina") | 0.2071223 | |
| | c("Capsicum_chacoense", "Lycianthoides_calycina", "Lycianthes_acapulcensis", "Lycianthes_rantonnetii") | 0.11201439 | |
| | c("Lycianthoides_calycina", "Lycianthes_rantonnetii") | 0.07676259 | |
| | c("Lycianthoides_calycina", "Lycianthes_acapulcensis") | 0.04964029 | |
| | Cummulative posterior | 1 | |
| ***Eophysaloides inflata*** (150 different sister groups) Showing first 10 | c("Eophysaloides_inflata", "Withania_somnifera") | 0.08503597 | yes |
| | c("Eophysaloides_inflata", "Leucophysalis_grandiflora") | 0.07129496 | |
| | c("Calliphysalis_carpenteri", "Eophysaloides_inflata") | 0.06856115 | |
| | c("Eophysaloides_inflata", "Oryctes_nevadensis") | 0.06633094 | |
| | c("Cuatresia_harlingiana", "Eophysaloides_inflata") | 0.05942446 | |
| | c("Darcyanthus_spruceanus", "Deprea_abra_patriciae", "Deprea_pauciflora", "Eophysaloides_inflata") | 0.04971223 | |
| | c("Eophysaloides_inflata", "Nicandra_physalodes") | 0.04 | |
| | c("Athenaea_brasiliana", "Athenaea_cuspidata", "Cuatresia_harlingiana", | 0.0347482 | |

| | | | |
|---|---|---|---|
| | "Darcyanthus_spruceanus", "Deprea_abra_patriciae", "Deprea_pauciflora", "Eophysaloides_inflata") | | |
| | c("Athenaea_brasiliana", "Athenaea_cuspidata", "Darcyanthus_spruceanus", "Deprea_abra_patriciae", "Deprea_pauciflora", "Eophysaloides_inflata") | 0.03201439 | |
| | c("Eophysaloides_inflata", "Nectouxia_formosa", "Salpichroa_tristis") | 0.0281295 | |
| | Cummulative posterior in the first 10 sister groups | 0.5352518 | |
| ***Hyoscyosperma daturoides*** (201 different sampled sister groups) Showing first 10 | c("Datura_ferox", "K453A_Hyoscyosperma_daturoides") | 0.605611511 | yes |
| | c("Discopodium_penninervium", "K453A_Hyoscyosperma_daturoides") | 0.019352518 | |
| | c("Exodeconus_miersii", "K453A_Hyoscyosperma_daturoides") | 0.018201439 | |
| | c("Capsicum_chacoense", "K453A_Hyoscyosperma_daturoides") | 0.016834532 | |
| | c("Jaltomata_bicolor", "Jaltomata_grandiflora", "K453A_Hyoscyosperma_daturoides") | 0.015611511 | |
| | c("Cuatresia_harlingiana", "K453A_Hyoscyosperma_daturoides") | 0.01028777 | |
| | c("K453A_Hyoscyosperma_daturoides", "Salpichroa_tristis") | 0.009496403 | |
| | c("K453A_Hyoscyosperma_daturoides", "Solanum_jamaicense") | 0.009352518 | |
| | c("K453A_Hyoscyosperma_daturoides", "Solanum_torvum") | 0.008561151 | |
| | c("K453A_Hyoscyosperma_daturoides", "Solanum_montanum", "Solanum_pinnatum") | 0.008417266 | |
| | Cummulative posterior in the first 10 sister groups | 0.721726619 | |
| ***Thanatosperma minutum*** (220 different sampled sister groups) Showing first 10 | c("2003_0073_Thanatosperma_minutum", "Anisodus_tanguticus") | 0.15431655 | yes |
| | c("2003_0073_Thanatosperma_minutum", "Atropa_belladonna") | 0.09964029 | |
| | c("2003_0073_Thanatosperma_minutum", "Solanum_caripense") | 0.08748201 | |
| | c("2003_0073_Thanatosperma_minutum", "Hyoscyamus_niger") | 0.05482014 | |
| | c("2003_0073_Thanatosperma_minutum", "Solanum_microdontum") | 0.04676259 | |
| | c("2003_0073_Thanatosperma_minutum", "Solanum_palustre") | 0.04294964 | |
| | c("2003_0073_Thanatosperma_minutum", "Przewalskia_tangutica") | 0.03323741 | |

| | | | |
|---|---|---|---|
| | c("2003_0073_Thanatosperma_minutum", "Lycianthes_rantonnetii") | 0.02992806 | |
| | c("2003_0073_Thanatosperma_minutum", "Atropanthe_sinensis") | 0.02856115 | |
| | c("2003_0073_Thanatosperma_minutum", "Scopolia_carniolica") | 0.02553957 | |
| | Cummulative posterior in the first 10 sister groups | 0.60323741 | |
| ***Solanoides dorofeevii*** (154 different sister groups) Showing first 10 | c("Calliphysalis_carpenteri", "K530_Solanoides_dorofeevii") | 0.06330935 | yes |
| | c("K530_Solanoides_dorofeevii", "Physaliastrum_heterophyllum") | 0.05625899 | |
| | c("Exodeconus_miersii", "K530_Solanoides_dorofeevii") | 0.04690647 | |
| | c("Darcyanthus_spruceanus", "Deprea_abra_patriciae", "Deprea_pauciflora", "K530_Solanoides_dorofeevii") | 0.04676259 | |
| | c("Capsicum_chacoense", "K530_Solanoides_dorofeevii") | 0.04136691 | |
| | c("Alkekengi_officinarum", "K530_Solanoides_dorofeevii") | 0.03496403 | |
| | c("Capsicophysalis_potosina", "K530_Solanoides_dorofeevii") | 0.03345324 | |
| | c("Athenaea_brasiliana", "Athenaea_cuspidata", "K530_Solanoides_dorofeevii") | 0.03201439 | |
| | c("Chamaesaracha_arida", "K530_Solanoides_dorofeevii") | 0.02935252 | |
| | c("Cuatresia_harlingiana", "K530_Solanoides_dorofeevii") | 0.02848921 | |
| | Cummulative posterior in the first 10 sister groups | 0.4128777 | |
| ***Solanum miocenicum*** (214 different sister groups) Showing first 10 | c("Jaltomata_bicolor", "Jaltomata_grandiflora", "K528_Solanum_miocenicum") | 0.05129496 | no |
| | c("K528_Solanum_miocenicum", "Withania_somnifera") | 0.05 | |
| | c("Exodeconus_miersii", "K528_Solanum_miocenicum") | 0.03769784 | |
| | c("Calliphysalis_carpenteri", "K528_Solanum_miocenicum") | 0.03604317 | |
| | c("K528_Solanum_miocenicum", "Solanum_corymbosum") | 0.03453237 | |
| | c("Archiphysalis_chamaesarachoides", "K528_Solanum_miocenicum") | 0.03417266 | |
| | c("K528_Solanum_miocenicum", "Lycium_americanum") | 0.03410072 | |
| | c("K528_Solanum_miocenicum", "Salpichroa_tristis") | 0.02755396 | |
| | c("K528_Solanum_miocenicum", "Solanum_jamaicense") | 0.02395683 | |
| | c("Capsicophysalis_potosina", "K528_Solanum_miocenicum") | 0.02151079 | |

| | | | |
|---|---|---|---|
| | Cummulative posterior in the first 10 sister groups | 0.3508633 | |
| ***Albionites arnensis*** (133 different sister groups) Showing first 10 | c("Heteranthia_decipiens", "Melananthus_guatemalensis", "Schwenckia_americana", "V_40898_Albionites_arnensis") | 0.11208633 | yes |
| | c("Benthamiella_patagonica", "Combera_paradoxa", "Pantacantha_ameghinoi", "V_40898_Albionites_arnensis") | 0.08705036 | |
| | c("Reyesia_chilensis", "Salpiglossis_sinuata", "V_40898_Albionites_arnensis") | 0.0642446 | |
| | c("Calibrachoa_humilis", "Fabiana_imbricata", "Petunia_interior", "V_40898_Albionites_arnensis") | 0.0528777 | |
| | c("Schizanthus_grahamii", "V_40898_Albionites_arnensis") | 0.04748201 | |
| | c("Bouchetia_anomala", "Hunzikeria_texana", "Leptoglossis_darcyana", "Nierembergia_linariifolia", "Plowmania_nyctaginoides", "V_40898_Albionites_arnensis") | 0.04582734 | |
| | c("Protoschwenkia_mandonii", "V_40898_Albionites_arnensis") | 0.04280576 | |
| | c("Nicotiana_wigandioides", "V_40898_Albionites_arnensis") | 0.04158273 | |
| | c("Heteranthia_decipiens", "Melananthus_guatemalensis", "V_40898_Albionites_arnensis") | 0.03035971 | |
| | c("Schultesianthus_leucanthus", "V_40898_Albionites_arnensis") | 0.02964029 | |
| | Cummulative posterior in the first 10 sister groups | 0.55395683 | |
| ***Nephrosemen reticulatum*** (237 different sister groups) Showing first 10 | c("UF6500_Nephrosemen_reticulatum", "Vestia_foetida") | 0.06266187 | no |
| | c("Protoschwenkia_mandonii", "UF6500_Nephrosemen_reticulatum") | 0.04661871 | |
| | c("Schizanthus_grahamii", "UF6500_Nephrosemen_reticulatum") | 0.04446043 | |
| | c("Nicotiana_wigandioides", "UF6500_Nephrosemen_reticulatum") | 0.0394964 | |
| | c("Reyesia_chilensis", "UF6500_Nephrosemen_reticulatum") | 0.03453237 | |
| | c("Leptoglossis_darcyana", "UF6500_Nephrosemen_reticulatum") | 0.03359712 | |
| | c("Reyesia_chilensis", "Salpiglossis_sinuata", "UF6500_Nephrosemen_reticulatum") | 0.03338129 | |
| | c("Schwenckia_americana", "UF6500_Nephrosemen_reticulatum") | 0.02906475 | |
| | c("Browallia_americana", "Streptosolen_jamesonii", "UF6500_Nephrosemen_reticulatum") | 0.0271223 | |

| | | | |
|---|---|---|---|
| | c("Symonanthus_aromaticus", "UF6500_Nephrosemen_reticulatum") | 0.02561151 | |
| | Cummulative posterior in the first 10 sister groups | 0.37654675 | |
| ***Capsicum pliocenicum*** (227 different sister groups) Showing first 10 | c("RGM792900_Capsicum_pliocenicum", "Chamaesaracha_arida") | 0.09935252 | no |
| | c("RGM792900_Capsicum_pliocenicum", "Nicandra_physalodes") | 0.05071942 | |
| | c("RGM792900_Capsicum_pliocenicum", "Witheringia_solanacea") | 0.04863309 | |
| | c("RGM792900_Capsicum_pliocenicum", "Nothocestrum_latifolium") | 0.04827338 | |
| | c("RGM792900_Capsicum_pliocenicum", "Solanum_pinnatum") | 0.0457554 | |
| | c("RGM792900_Capsicum_pliocenicum", "Solanum_hieronymi") | 0.04179856 | |
| | c("RGM792900_Capsicum_pliocenicum", "Tubocapsicum_anomalum") | 0.02697842 | |
| | c("Archiphysalis_chamaesarachoides", "RGM792900_Capsicum_pliocenicum") | 0.02467626 | |
| | c("Athenaea_brasiliana", "Athenaea_cuspidata", "RGM792900_Capsicum_pliocenicum") | 0.02381295 | |
| | c("RGM792900_Capsicum_pliocenicum", "Schraderanthus_viscosus") | 0.02165468 | |
| | Cummulative posterior in the first 10 sister groups | 0.43165468 | |
| ***Solanum foveolatum*** (167 different sister groups) Showing first 10 | c("Jaborosa_laciniata", "K587B_Solanum_foveolatum") | 0.05920863 | no |
| | c("Jaltomata_bicolor", "Jaltomata_grandiflora", "K587B_Solanum_foveolatum") | 0.05705036 | |
| | c("K587B_Solanum_foveolatum", "Solanum_thelopodium") | 0.04503597 | |
| | c("Capsicum_chacoense", "K587B_Solanum_foveolatum") | 0.04057554 | |
| | c("K587B_Solanum_foveolatum", "Solanum_barbisetum", "Solanum_crinitum", "Solanum_hieronymi", "Solanum_jamaicense", "Solanum_mahoriense", "Solanum_oedipus", "Solanum_sejunctum", "Solanum_sisymbriifolium", "Solanum_torvum") | 0.03582734 | |
| | c("K587B_Solanum_foveolatum", "Solanum_montanum", "Solanum_pinnatum") | 0.0352518 | |
| | c("K587B_Solanum_foveolatum", "Solanum_barbisetum", "Solanum_hieronymi", "Solanum_jamaicense", "Solanum_mahoriense", "Solanum_oedipus", "Solanum_sejunctum", "Solanum_torvum") | 0.03230216 | |

| | | | |
|---|---|---|---|
| | c("K587B_Solanum_foveolatum", "Solanum_evolvuloides") | 0.02978417 | |
| | c("K587B_Solanum_foveolatum", "Solanum_annuum", "Solanum_corymbosum", "Solanum_salamancae", "Solanum_sinuatirecurvum", "Solanum_weddellii") | 0.02884892 | |
| | c("Exodeconus_miersii", "K587B_Solanum_foveolatum") | 0.02863309 | |
| | Cummulative posterior in the first 10 sister groups | 0.39251798 | |
| ***Solanispermum reniforme*** (228 different sister groups) Showing first 10 | c("Quincula_lobata", "V_40891_Solanispermum_reniforme") | 0.06676259 | no |
| | c("Solanum_sejunctum", "V_40891_Solanispermum_reniforme") | 0.05741007 | |
| | c("Solanum_mahoriense", "V_40891_Solanispermum_reniforme") | 0.05151079 | |
| | c("Tubocapsicum_anomalum", "V_40891_Solanispermum_reniforme") | 0.03683453 | |
| | c("Nectouxia_formosa", "V_40891_Solanispermum_reniforme") | 0.03338129 | |
| | c("Solandra_grandiflora", "V_40891_Solanispermum_reniforme") | 0.02856115 | |
| | c("Exodeconus_miersii", "V_40891_Solanispermum_reniforme") | 0.02539568 | |
| | c("Solanum_montanum", "Solanum_pinnatum", "V_40891_Solanispermum_reniforme") | 0.02323741 | |
| | c("Oryctes_nevadensis", "V_40891_Solanispermum_reniforme") | 0.0218705 | |
| | c("Solanum_thelopodium", "V_40891_Solanispermum_reniforme") | 0.02151079 | |
| | Cummulative posterior in the first 10 sister groups | 0.3664748 | |

### LITERATURE CITED

**Chandler, M.E.J.** 1957. The Oligocene flora of the Bovey Tracey lake basin, Devonshire. Bull. Brit. Mus. (Nat. Hist.), Geol. 3: 71–123. https://doi.org/10.5962/p.313850

**Chandler, M.E.J.** 1962. The Lower Tertiary floras of Southern England II. Flora of the Pipe-Clay Series of Dorset (Lower Bagshot). London: British Museum (Natural History).

**Deanna, R., Martínez, C., Manchester, S., Wilf, P., Campos, A., Knapp, S., Chiarini, F.E., Barboza, G.E., Bernardello, G., Sauquet, H., Dean, E., Orejuela, A. & Smith, S.D.** 2023. Fossil berries reveal global radiation of the nightshade family by the early Cenozoic. *New Phytol.* 238: 2685–2697. https://doi.org/10.1111/nph.18904

**Deanna, R., Wilf, P. & Gandolfo, M.A.** 2020. New physaloid fruit-fossil species from early Eocene South America. *Am. J. Bot.* 107: 1749–1762. https://doi.org/10.1002/ajb2.1565

**Deanna, R., Hvalj, A.V., Martinetto, E., Knapp, S., Sadowski E.-M., Manchester, S., Campos, A., Fernandez, V., Barboza, G.E., Sauquet, H., Dean, E., Särkinen, T., Chiarini, F.E., Bernardello, G., Smith, S.D.** 2025. Seed fossil record of Solanaceae revisited. Taxon, submitted. bioRxiv https://www.biorxiv.org/content/10.1101/2025.07.03.662944v1

**Manchester, S.R.** 1994. Fruits and seeds of the Middle Eocene Nut Beds flora, Clarno Formation, Oregon. *Paleontogr. Am.* 58: 1-205. https://biostor.org/reference/240376

**Negru A.G.** 1986. Meoticheskaya flora severo-zapadnogo Prichernomor'ya [Maeotic flora of the north-western Black Sea region]. Chișinău: Știința.

**Wilf. P., Carvalho, M.R., Gandolfo, M.A. & Cúneo, N.R.** 2017. Eocene lantern fruits from Gondwanan Patagonia and the early origins of Solanaceae. *Science* 355: 71–75. https://doi.org/10.1126/science.aag2737
